# Mapping the landscape of social behavior

**DOI:** 10.1101/2024.09.27.615451

**Authors:** Ugne Klibaite, Tianqing Li, Diego Aldarondo, Jumana F. Akoad, Bence P. Ölveczky, Timothy W. Dunn

**Author notes:** These authors contributed equally (Ugne Klibaite, Tianqing Li).

## Abstract

Social interaction is integral to animal behavior. However, we lack tools to describe it with quantitative rigor, limiting our understanding of its principles and neuropsychiatric disorders, like autism, that perturb it. Here, we present a technique for high-resolution 3D tracking of postural dynamics and social touch in freely interacting animals, solving the challenging subject occlusion and part assignment problems using 3D geometric reasoning, graph neural networks, and semi-supervised learning. We collected over 140 million 3D postures in interacting rodents, featuring new monogenic autism rat lines lacking reports of social behavioral phenotypes. Using a novel multi-scale embedding approach, we identified a rich landscape of stereotyped actions, interactions, synchrony, and body contact. This enhanced phenotyping revealed a spectrum of changes in autism models and in response to amphetamine that were inaccessible to conventional measurements. Our framework and large library of interactions will greatly facilitate studies of social behaviors and their neurobiological underpinnings.

## Introduction

The study of social interactions is essential for understanding evolutionary, ecological, and neurobiological principles of animal behavior. While social behavior is multifaceted, much of it is expressed through body movements that reflect internal states, goals, and inter-animal communication. These gestures are often subtle, necessitating a method for capturing the precise 3D kinematics of interacting animals. Such measurements could then be analyzed to define and quantify social behaviors in rigorous and reproducible ways, enabling sophisticated interrogations of social dynamics and their biological underpinnings. Further, an automated and scalable method would permit comprehensive behavioral screens to characterize the full variety of social phenotypes, including those occurring in autism and other neurodevelopmental disorders.

These goals have yet to be met. Today, social phenotyping is typically done by manually scoring video recordings or, increasingly, by automatically tracking the position and orientation of interacting animals. For instance, in studies of rodent models of autism, metrics of sociality are commonly derived from the fraction of time an animal spends in the proximity of a caged conspecific^1^. While some studies expand these metrics to include the occurrence of a limited set of social action types^2^, the manual annotation that goes into this is laborious, inconsistent, and limited to predefined categories. For autism and other conditions diagnosed using only behavioral criteria, such coarseness makes it difficult to establish animal model face validity, and also to translate findings out of the lab^3^.

To reliably discern and describe subtle variations in social phenotypes^4^, we need an approach that can capture the diversity and nuances of social behavior. That means precisely tracking how interacting animals move, synchronize, and make physical contact, plus analytics for classifying, interpreting, and comparing these elements of interaction. Capturing the nature of physical contact between social partners is particularly important given recent interest in social touch as a rewarding stimulus and its impairment in autism spectrum disorders (ASD)^5,6^. Further, these analytics must address multiple timescales of social interactions, from short, stereotyped movements and engagements to behavioral patterns that evolve over longer times, such as group coordination, instigation, communication, and arousal^7^.

Recent advances in video-based behavioral quantification have enabled more granular and scalable measurements of social interactions, but these methods are still fundamentally limited in resolution. Instead of the coarse position and orientation tracking afforded by classical computer vision, convolutional neural networks (CNNs) can now track anatomical keypoints of interacting animals in 2D^8,9^. With these 2D kinematic descriptions, classifiers can be trained to detect different types of social interactions once they are enumerated and labeled by humans^10^. Tracked 2D kinematics can also be clustered without explicit human supervision, facilitating the discovery and annotation of novel social action patterns^11–13^. Nevertheless, 2D measurements are inherently limited in terms of the body parts that can be tracked, and perspective ambiguities make it difficult to derive reliable body kinematics from a single view. Thus, while these 2D innovations are significant, they are not yet precise enough to provide comprehensive descriptions of social behavior.

Reliable quantitative descriptions of social interactions would be greatly facilitated by high-resolution 3D tracking of animal pose – that is, the positions of actuatable body parts, including limbs – which would permit precise spatiotemporal profiling of coordinated kinematics and body contact. While such 3D pose tracking methods have been developed for single animals^14–19^, extending them to social contexts is far from trivial. Single-animal methods are not equipped to deal with the complexities of multi-animal environments, where animal identities must be tracked consistently and bodies are often hidden from view. Depth imaging has been used for social tracking but cannot faithfully track limb movement^20–22^. Tracking animals as single 3D points within groups has been foundational for studies of social and collective behavior^23,24^, but cannot and does not aim to measure detailed body movements^25^. 2D whole-body pose tracking methods could, in principle, be extended to 3D via triangulation across multiple camera viewpoints^26^, but inevitable and ubiquitous animal-animal occlusions make 2D-to-3D triangulation during interaction particularly challenging. Large marker-based 3D training datasets can be used to address occlusion difficulties, but markers themselves are often occluded on limbs and during close interaction, limiting these datasets to reduced keypoint sets on the head and trunk^27–29^. Thus, reliable whole-body 3D tracking of social behavior has yet to be achieved.

To address these problems, we developed a method for tracking highly resolved 3D postural kinematics in interacting animals using commercially available video cameras. Our 3D tracking approach, social-DANNCE (s-DANNCE), builds off of DANNCE, a deep neural network for high-resolution 3D markerless tracking in single animals^17^. s-DANNCE leverages semi-supervised learning, anatomical constraints, and a graph neural network to resolve animal-animal occlusions and reliably track pairs of interacting rats with performance surpassing previous approaches^26,27^ and rivaling that of human labelers. s-DANNCE is also flexible, supporting measurements across a range of experimental settings, including in larger groups of animals, and generalizes across species from rats to mice.

To parse high-resolution tracking data into comprehensive and quantitative descriptions of social behavior across spatiotemporal scales, we developed a new suite of computational tools for analyzing dyadic interactions (**Fig. 1a**), including the automatic identification of recurring and stereotyped social interaction motifs. We further developed a method to automatically identify instances of social touch by fitting a volumetric body model to 3D poses, thus providing quantitative access to a salient mode of social exchange^30^.

**Figure 1:**
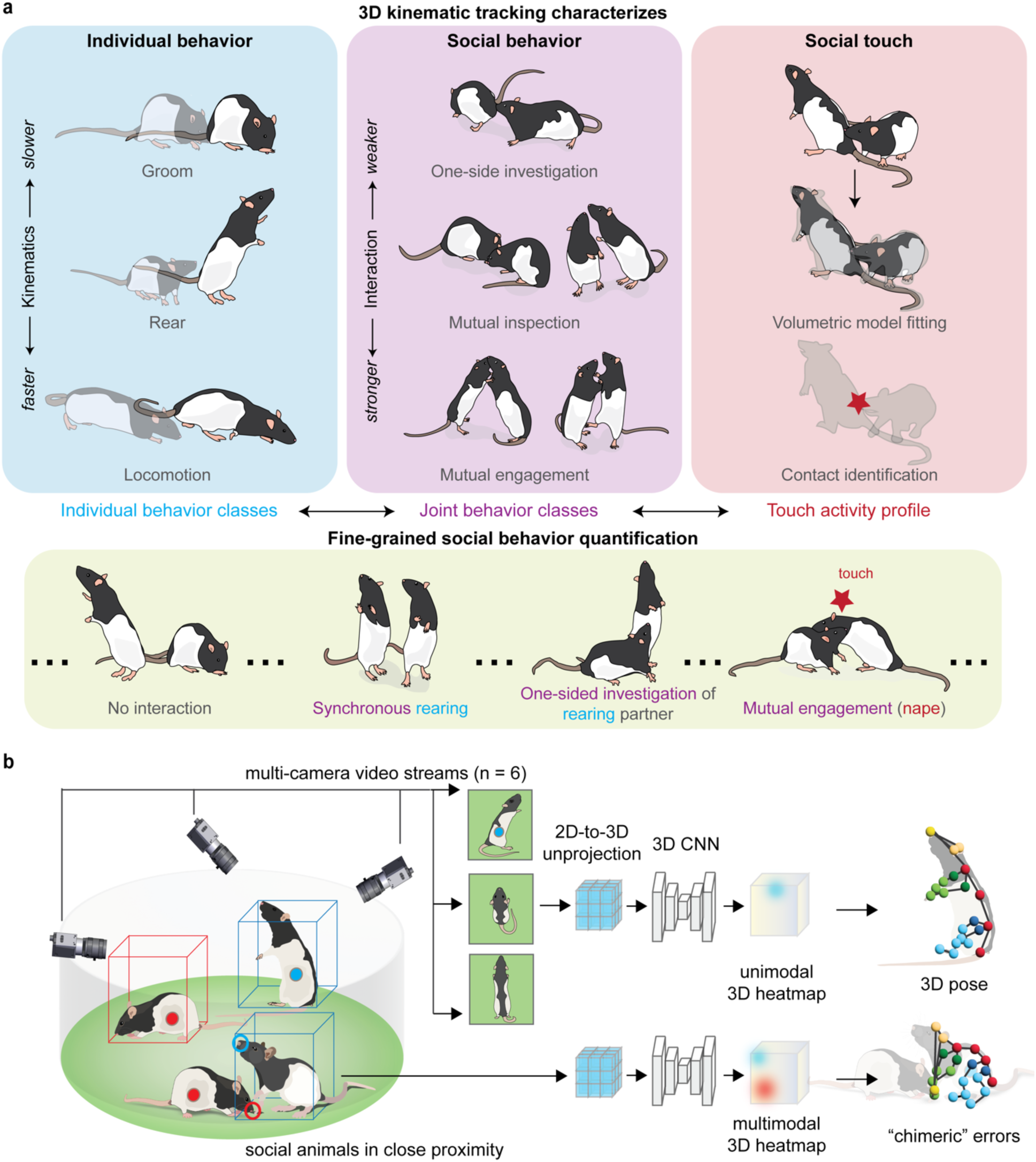
High-resolution 3D kinematic tracking enables fine-grained quantification of social behaviors. **a**, We introduce three levels of behavioral profiling based on the tracked 3D keypoints: individual behaviors (top left), social interaction behaviors which occur in dyads (top middle), and a method to profile how animals touch during interaction by using a 3D volumetric body representation (top right). Together, these levels of behavioral classification allow fine- grained curation and quantification of social behaviors in freely interacting animals (bottom). **b**, Schematic of the recording arena and the baseline DANNCE implementation, in which single animal pose estimation is separately performed within 3D volumes encapsulating different individuals. Top, six synchronized camera views capture behaviors of animals freely interacting in the circular arena. 3D volumetric inputs are constructed from simultaneously recorded multi- view images via projective geometry (**Method**) and processed by a 3D CNN to directly infer the 3D landmark positions. Bottom, failure cases for tracking social animals in close proximity, in which landmarks associated with different animals give rise to multimodal activations in the predicted 3D heatmaps and thus result in chimeric associations in the estimated 3D social postures. The usage of rat illustrations in **b** is under an [MIT license](https://github.com/eackermann/ratpack/blob/master/LICENSE).

To validate whether our framework enhanced social phenotyping, we administered amphetamine, a drug with clear and demonstrable social effects in humans but not in rodent models, to rats. In addition to inducing overall hyperactivity as expected, drug administration disrupted behavioral synchrony, as well as other canonical interaction motifs, and changed the distributions of touch across the bodies of animal pairs. In mice, social behavior mapping identified new differences between mouse strains commonly used as models of low and high sociability.

Our novel analysis approach also revealed subtle deviations from typical social behavior in seven rat models of autism. We found that rats from each of the models were impacted in distinct ways, with four of the seven demonstrating widespread shifts in social behavior. Indeed, our comprehensive behavioral phenotyping returned several phenotypically distinct candidate models of face-valid social phenotypes in rats. Beyond advancing the state-of-the art in social tracking, our study resulted in a large database of over 160 million rat 3D poses in lone and social contexts. We provide these data, together with our machine learning framework and detailed phenotyping results, as a new resource for the community to mine, model, and reference in the pursuit of a deeper understanding of social behavior.

## Results

### Animal-animal occlusions challenge 3D pose tracking during social behaviors

The state of the art for tracking animal movement during social behaviors is to use CNNs to locate 2D anatomical features, or keypoints, on bodies of interacting subjects, typically from videos recorded by a single top-down or bottom-up camera^8–10^. However, quantifying behavior in 2D pixel space, instead of descriptions of true 3D movements (in millimeters), introduces kinematic artifacts due to perspective and depth ambiguities^15,31^. Single camera measurements are sensitive to body occlusions, which increase in frequency during social interaction (**Supp.** Fig. 2a,b). In a top-down view, the animal’s own body occludes its appendages 76% of the time, while occluding the head and trunk for 5% and 10% of the time respectively (**Supp.** Fig. 2c). In a bottom-up view, the top of the body is always occluded, as are the forelimbs and head in elevated postures, such as rearing (**Supp.** Fig. 2d). Occlusions are even more pronounced during close social interaction, when one animal often obstructs the view of the other (approximately 3-fold and 1.5-fold more occlusions of the head and trunk, respectively, compared to when the animal is alone; **Supp.** Fig. 2c,d).

In principle, such occlusions and kinematic artifacts can be addressed via 3D tracking of body keypoints in multi-camera videos, as done in single animals^14–17,27^. To test the feasibility of this, we built a setup consisting of six synchronized 50 Hz color video cameras for recording rat dyadic interactions in a circular arena (**Fig. 1b, Supp.** Fig. 1**, Supp. Video 1**). We previously applied DANNCE, a 3D tracking method built on volumetric neural networks, to multiple animals^17,27^. This multi-animal implementation exploited the “top down” structure of DANNCE in which animals are first identified and then compartmentalized into 3D bounding boxes to make 3D pose estimates. While DANNCE reliably tracked 3D pose when animals were far apart, performance degraded significantly during close interactions. For example, we frequently observed ‘chimeric’ errors, where the inferred 3D pose incorrectly integrated body elements from its interaction partner (**Supp.** Fig. 3). This failure mode is due to animals being enclosed, at least partially, in the same 3D volume, leading to multiple candidate keypoint locations for individual body parts (**Fig. 1b *bottom*)**.

### social-DANNCE enables 3D tracking of social behaviors

To overcome the challenges of consistently and precisely tracking whole-body 3D kinematics during social behaviors, we developed s-DANNCE, which extends the precision of volumetric deep neural networks to closely interacting animals (**Fig. 2a, Supp.** Fig. 4). s-DANNCE combats pervasive chimeric prediction errors and other observed anatomical inconsistencies (collapsed poses and errant body segment lengths) by imposing explicit constraints on body anatomical structure during training, plus a graph neural network (GNN) module^32^ that enables skeleton-aware processing of extracted 3D image features. Further, s-DANNCE is able to leverage both labeled and unlabeled data during training, allowing the network to learn from large, diverse datasets that have not been annotated by users^19,33^.

**Figure 2:**
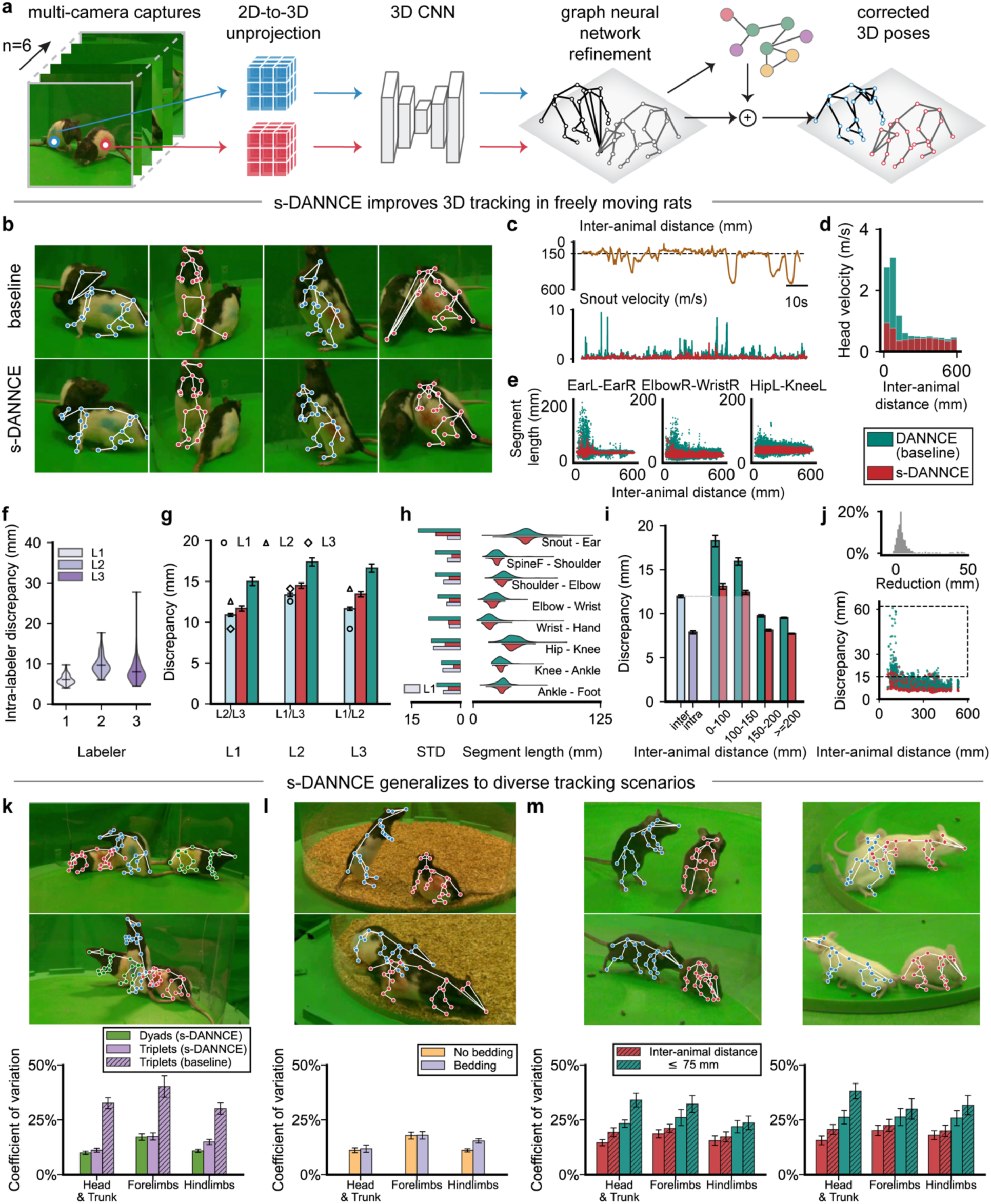
s-DANNCE improves 3D tracking of freely interacting animals. **a**, Schematic of the s- DANNCE pipeline, in which 3D volumes were separately constructed around each animal’s centroid from multi-camera (n = 6) images via the unprojection operation and then jointly processed by a 3D CNN and a graph neural network-based (GNN) refinement module (see model architecture details in **Supp.** Fig. 4a). **b,** Qualitative visualization of how chimeric errors during close animal interaction present in baseline (“multi-animal DANNCE”) predictions are resolved by s-DANNCE. **c-e** provide an examination of s-DANNCE’s tracking performance over an unannotated test video sequence (n = 90000 frames) in one rat engaging in dyadic interactions. **c**, Kinematic traces (t = 80 seconds) showing oscillations in the velocity of the animal snout, as derived from both models’ predictions (bottom). The top trace denotes distances between the paired animals. The same color scheme for the indicated methods is used in the remaining subplots. **d**, Head movement velocity as a function of inter-animal distance on 95% outliers, with a bin size of 50 mm. **e**, Scatter plots of body segment lengths as a function of inter-animal distance. **f-h,** Three human labelers independently annotated the same set of frames (n = 40 frames, 4 animals) with close animal social interactions twice to test inter-labeler variability. **f**, Violin plots of the intra-labeler annotation discrepancies, showing the means with minimum and maximum extrema. **g**, Bar plots of inter-labeling discrepancies (both human and machine annotations) against different sets of human ground truth (Labeler 1, 2, 3). Discrepancies for human annotations are merged for simplicity in each group. Error bars are 95% confidence intervals (CIs). **h**, Right, distributions of body segment lengths derived from different model predictions. Laterally symmetric body segments are merged for conciseness. Left, standard deviation (STD) of segment lengths derived from model predictions (baseline DANNCE, s-DANNCE) and average human annotations (Labeler 1-3). **i**, Bar plots of landmark localization discrepancy over a larger annotated dataset (n = 1373 frames, 2 animals, ground truth annotated by Labeler 1) withheld from model training and validation. Frames are grouped by the inter-animal distances (0-100, 100-150, 150-200, >=200 mm). Error bars are 95% CIs. The average intra- and inter- discrepancies of the three human labelers (light blue and purple bars, respectively), as computed from **f** and **g,** are included for comparison. **j**, Bottom, scatter plot of localization discrepancy as a function of inter-animal distance over the same set of samples in **i**. Top, discrepancy reduction achieved by s-DANNCE, for frames where the baseline predictions yielded Euclidean errors greater than 15 mm (n = 261 samples in the dashed box region). **k-m,** Demonstration of s-DANNCE’s generalization capacities to diverse tracking scenarios, respectively in triplets of rats, in arenas with bedding, and in two strains of mice. Top row, examples of s-DANNCE predictions. Bottom row, coefficients of variation of segment lengths derived from different model predictions across n = 12 tracked instances (n = 4 triplet recordings or n = 6 dyad recordings). Segments specified in **h** are further grouped by their underlying body regions (head, trunk, forelimbs, hindlimbs) for conciseness.

To test whether s-DANNCE mitigated naive DANNCE baseline tracking issues, we trained both on the same set of 6-camera video frames of freely interacting rat pairs (n = 423 manually annotated 23-keypoint 3D poses and n = 910 unlabeled frames) and tested the models on videos of animals not used for training. We found that s-DANNCE, in contrast to baseline DANNCE, produced robust tracking without chimeric errors (**Fig. 2b, Supp.** Fig. 3b-e**, Supp.** Fig. 5). Baseline DANNCE exhibited large fluctuations in keypoint velocities (**Fig. 2c,d**) and body segment lengths (**Fig. 2e**) when animals were close to each other (a threshold of 150 mm inter-animal distance, or approximately 2/3 of rat body length), indicating frequent social tracking errors. These errors were resolved by s-DANNCE (65% improvement to head velocity stability and 60% improvement in limb segment length stability for s-DANNCE over DANNCE, **Fig. 2d,e**).

To more directly assess s-DANNCE performance, we quantified how close tracked 3D keypoints were to human annotations on a test set of close social interaction frames. We had three different labelers annotate the same test set for these comparisons (n = 80 3D poses, with poses annotated twice by each labeler for measuring intra-labeler consistency), allowing us to account for human labeling biases when interpreting model performance metrics (**Fig. 2f**). We found s-DANNCE rivaled human labeler precision, with s-DANNCE placing keypoints closer to human annotations than one of the labelers in the group (relative to Labeler 1 and Labeler 3, s- DANNCE 12.56 mm vs. Labeler 2 13.33 mm discrepancy) and nearing inter-labeler precision on average (**Fig. 2g**; s-DANNCE 13.20 ± 0.19 mm discrepancy relative to all labelers; inter-labeler 11.95 ± 0.16 mm discrepancy over all labeler pairs; mean ± 95% CI). In contrast, baseline DANNCE, as well as a recent image masking approach for 3D social tracking^26^, yielded consistently larger discrepancies relative to human annotators (baseline DANNCE 16.34 ± 0.29 mm, **Fig. 2g**; image masking 18.88 ± 0.24 mm, **Supp.** Fig. 7). s-DANNCE also produced 3D pose estimates that were more anatomically plausible, evidenced by consistent body segment lengths across postures (s-DANNCE 0.14 ± 0.022, human labeler 0.17 ± 0.019, baseline DANNCE 0.30 ± 0.057, coefficient of variation ± 95% CI, **Fig. 2h**). We further examined the performance of s-DANNCE over a broader variety of social interactions in a larger test recording (n = 2746 3D postures in one pair of rats, annotated by Labeler 1). As anticipated, the largest improvements were found when animals were close and occluding each other during the most interactive social behaviors; specifically, s-DANNCE yielded 6.7-fold better precision relative to baseline DANNCE, when adjusting for human inter-labeling uncertainty (0.72 ± 0.20 mm vs. 4.84 ± 0.36 mm, **Fig. 2i**; 12.67 ± 0.19 mm vs. 16.80 ± 0.35 mm without adjustments). s- DANNCE outperformed baseline in 94% of the test frames with the remaining 6% within the inter-labeler margin of error (**Fig. 2j**).

s-DANNCE extended to a wide range of experimental contexts, including to recordings of interacting mice. While training frames came from just one strain of rats recorded in one arena, s-DANNCE generalized immediately to new arenas, rat strains, and larger social groups without additional training (**Fig. 2k, Supp.** Fig. 5). s-DANNCE also accurately tracked social behavior in recordings with bedding when a small number of bedding frames were added for training (**Fig. 2l**). Given a small number of labeled frames for fine-tuning, s-DANNCE could track two different and visually distinct strains of freely interacting mice (**Fig. 2m, Supp.** Fig. 6). Our approach thus broadly supports high-resolution quantification of social behavior without introducing heavy annotation burdens.

### 3D kinematic profiling reveals novel social phenotypes

High-resolution and continuous whole-body 3D kinematic measurements provide an opportunity to quantitatively map social behavioral repertoires across animals and experimental conditions. To characterize social behavior, we first examined how individual repertoires changed in lone vs. social contexts using a method that clusters behavior into stereotyped motifs based on an individual subject’s 3D postural dynamics, similar to previous works^34–36^. Briefly, for each movie we derived a time series of high-dimensional features from 3D pose dynamics and applied nonlinear dimensionality reduction to generate a 2D behavior map. We performed spatial clustering of this map to identify stereotyped action classes which were used to assign a behavioral label at each time point of tracked movement (**Fig. 3a,e**). We manually reviewed and annotated behavioral snippets sampled from each cluster on the behavior map, revealing an action space organized into regions representing 9 human-annotated high-level action classes (HLACs) and 162 low-level action clusters (LLACs) distinguishable by 3D kinematics (**Supp.** Fig. 9, **Supp. Table 2, Supp. Videos 3-4**, **Methods**). This allowed us to compare behavior at the level of fine stereotyped actions (LLACs) or with coarser, more interpretable operational definitions (HLACs). When we mapped full-body 3D kinematics from both single- and multi-animal recordings from wild-type Long Evans male rats (N = 6 animals, N = 24 pairings, N=24 lone recordings) into the behavioral space, we captured the behavioral shift associated with social context, revealing 116 LLACs (of 162 total) that exhibited significant changes in how often they were expressed (their behavioral ‘occupancies’) during social sessions (**Fig. 3a, *right***).

**Figure 3:**
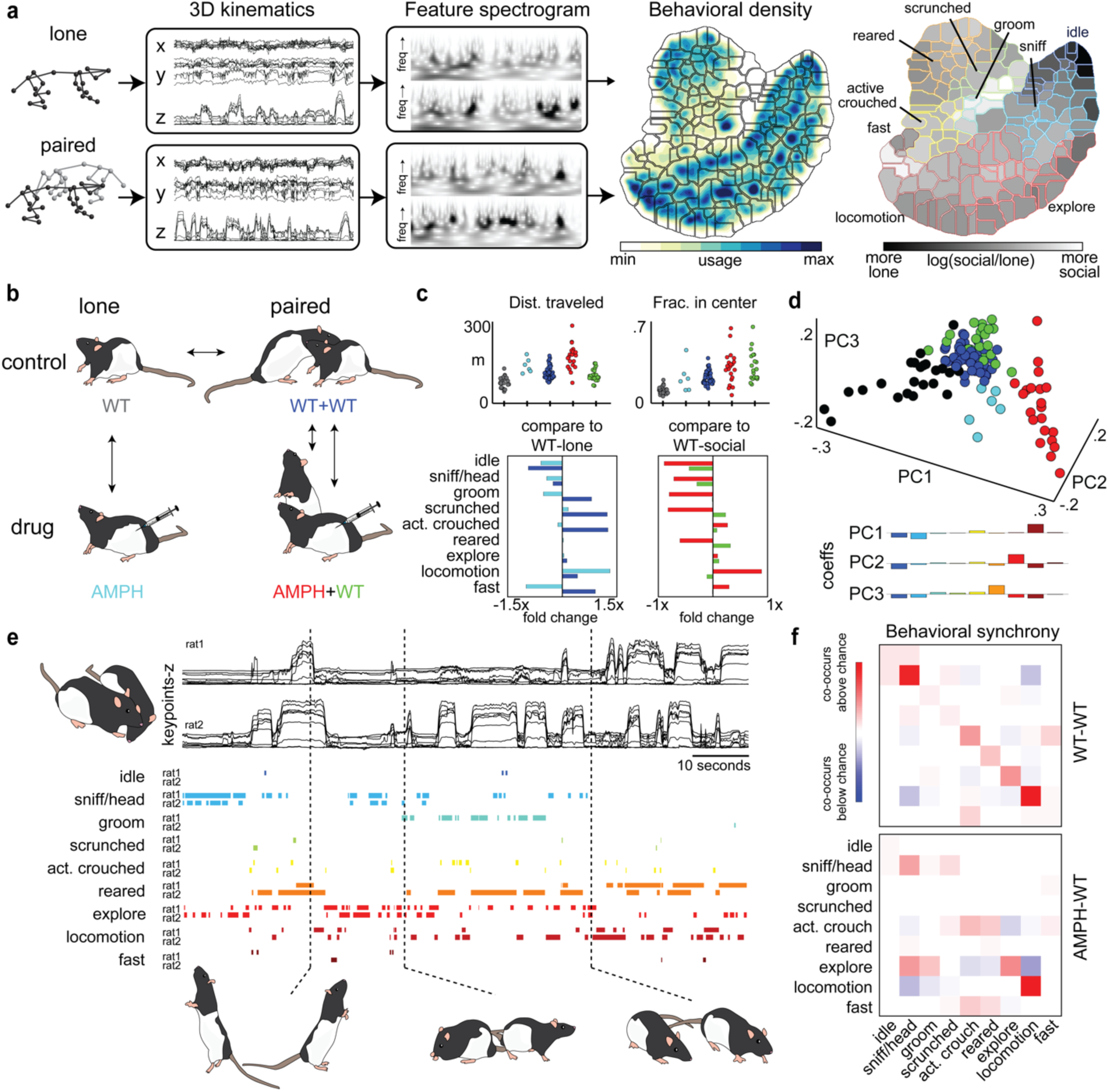
Spatio-temporal clustering demonstrates kinematic changes in response to social context and administration of a stimulant. **a**, Kinematic data from s-DANNCE is transformed to a high-dimensional signal which captures postural and movement features (wavelet decomposition of 15 postural PCs at 25 dyadically spaced frequencies .5-20 Hz, keypoint height, and keypoint speed), and then clustered to reveal a 2D behavioral map where behavioral clusters are organized into nine high-level hand-annotated behavioral descriptions. On the right, each low-level action cluster (LLAC) is outlined in the color representing the corresponding high-level action cluster (HLAC), and the fill color corresponds to how frequently the behavior occurred in the social context. **b**, Experimental setup: behavioral contexts tested here are WT-lone, WT-social, AMPH-lone, AMPH-social, and the social partner to the amphetamine-dosed animal (WT-partner) **c**, Coarse metrics of distance traveled over each 30-minute recording (top left) and the fraction of time spent in the center of the arena (top right) derived from the animal’s center of mass are shown for each experimental group. Coarse label behavioral shifts across experimental groups are shown in comparison to the WT-lone condition (bottom left) and the WT-social condition (bottom right), **d**, PCA reveals context-specific clustering across individual experiments. **e**, Time series traces of kinematics (z-axis only) and the assigned behavior classes over time show temporal structure in paired behavior. Bottom, snapshots of synchronized moments from analyzed data. **f,** Synchrony of paired behavioral time series, calculated as the likelihood of a pair of behaviors occurring together above chance, reveals asymmetry in paired behavior when one animal is administered amphetamine prior to social interaction.

These quantitative descriptions enable new, comprehensive, and automated ways of characterizing how drugs and diseases affect social behavior. To demonstrate the the power of this approach, we tracked and profiled rat behavior in lone and social contexts with and without the administration of amphetamine (n = 75 recordings across social pairings and experimental conditions from N = 6 animals; total tracked 3D poses used for analysis n = 10.8 million) (**Fig. 3b, Supp. Video 2**). While it has been shown that amphetamine affects rodent social behavior, these studies relied on manual scoring of a limited set of behavioral categories (e.g., grooming, fighting, overall movement speed) and did not dissociate social effects from amphetamine’s general effects on behavior^37,38^, which are substantial (cf. **Fig. 3a, Supp.** Fig. 18). Our quantitative profiling automatically identified a variety of behavioral changes in response to amphetamine that were specific to social contexts and not readily detectable using coarser measures (**Fig. 3c,d**). For instance, while amphetamine did not affect rearing behaviors in lone recordings, in social recordings it produced a pronounced decrease in frequency of rearing relative to WT controls (lone .04 fold increase from controls, social .6 fold decrease from controls, **Fig. 3c**). Overall, large shifts in the behavioral repertoire were induced by the social context and further modulated by amphetamine. These shifts were consistent across both animals and experimental sessions (**Fig. 3d, Supp.** Fig. 18). In contrast, conventional measures, such as distance traveled, were too coarse to distinguish between experimental groups (**Fig. 3c**).

Labeling the HLACs expressed simultaneously by social partners allowed us to analyze behavioral synchrony (**Fig. 3f**), a hallmark of social behavior that has been difficult to characterize or quantify without manual human annotation^39,40^. Paired animals have been shown to display behavioral alignment at short time scales as well as in their behavioral usage across the length of interactions^11^. We calculated how likely each combination of HLACs was to co-occur between partners above what was expected by chance given each individual’s occupancy across HLACs (**Fig. 3f, Methods**). In wild-type pairings, the highest synchrony values were observed between behavioral classes of the same type (particularly simultaneous *sniffing* and *locomotion*), providing new insights into how rats synchronize their actions during social engagement.

In addition to altering the frequency of behavior expression in individual animals within social contexts (cf **Fig. 3c**), amphetamine administered to one animal in a social pairing reduced overall behavioral synchrony, although simultaneous locomotion was preserved. The HLACs paired animals used simultaneously when an amphetamine animal was present were often from different high-level classes (**Fig. 3f**). While these asymmetries could reflect one-sided aggressive behaviors, which are common human responses to the drug^41^, disambiguating these from less assertive one-sided interactions requires analysis of additional behavioral variables.

### Dyadic embedding identifies stereotyped motifs of social interaction across species

While revealing, the kinematic profiles of individual animals do not capture the coordination of body movements unique to social interaction. To identify recurring and stereotyped social action motifs, we developed dyadic embedding, a novel classification method based on shared inter-animal features (**Fig. 4a, Supp.** Fig. 9). Broadly, our method uses the posture, dynamics, and inter-animal features from 3D kinematic tracking of two animals to produce a behavioral map that can parse social behavior into stereotyped joint classes or motifs. The method effectively treats the two animals as a single system and exploits the fact that distinct social motifs manifest as identifiable patterns in the relative positions and orientations of 3D poses between animals. For example, when one animal is sniffing and following the other from behind, the animals are consistently oriented in the same direction with heads maintained at a characteristic distance. Alternatively, when animals are investigating each other nose-to-nose, they are oriented in opposite directions with heads close together. We automatically discover each of these and numerous other social motifs from the joint embedding as in the single animal case (see **Methods**). Crucially, social motifs should be sensitive to the egocentric perspectives of individual animals, preserving each individual’s identity and context, as the meaning of an interaction is actor-dependent **(Supp.** Fig. 9).

**Figure 4:**
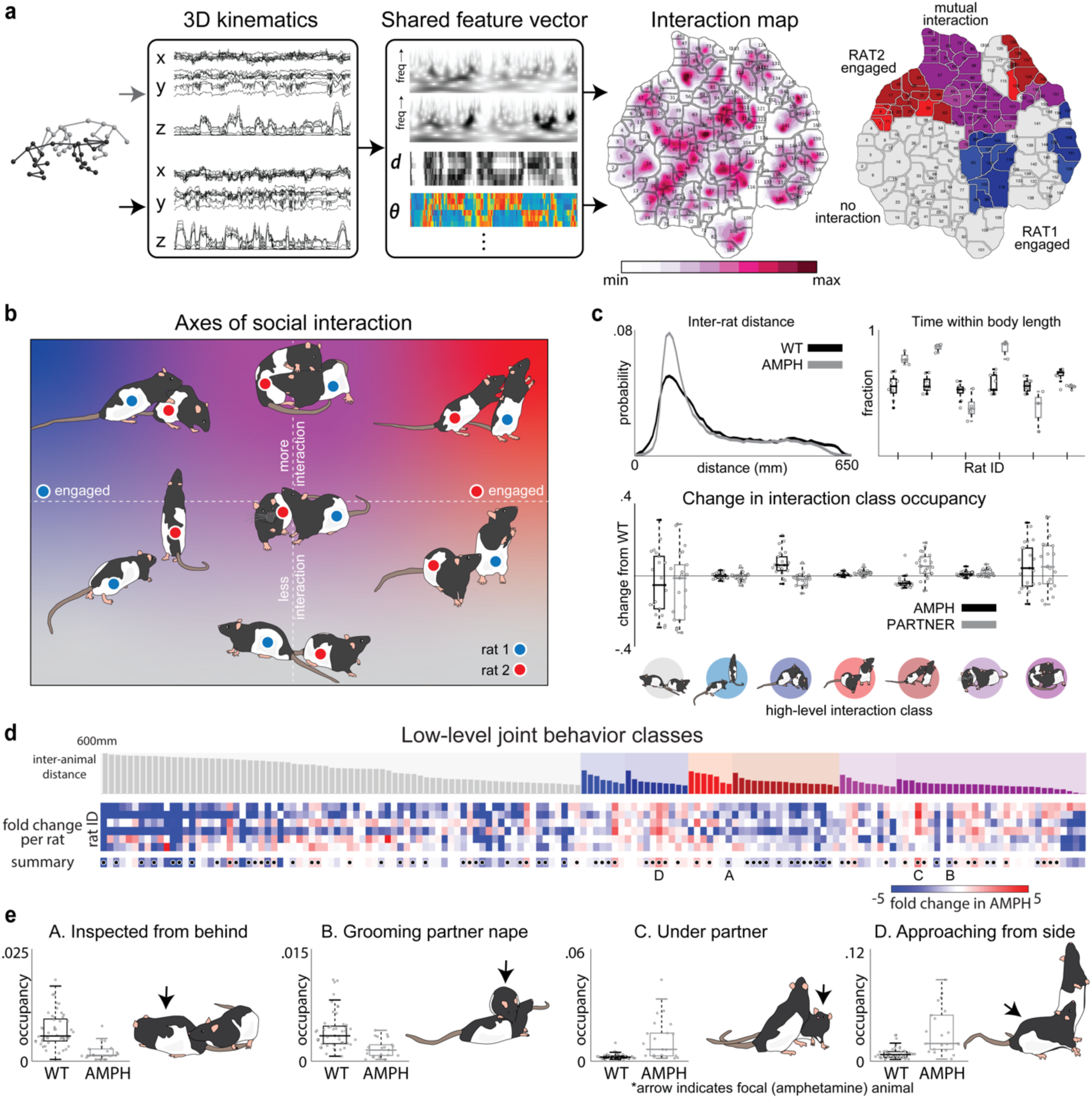
Dyadic feature embedding reveals social differences when amphetamine is administered to an animal before social interaction. **a**, Social embedding incorporates kinematic time-series from interacting animals, and features derived from these (wavelet decomposition of six PCs describing the shared configuration of both animals at 25 dyadically spaced frequencies .5-20 Hz, inter-animal distances and heading deflections from the snout and two points along the spine), to produce social interaction clusters in an unsupervised manner. **b**, Snapshots from representative behaviors from each of the seven high-level social interaction categories (HLJCs) which emerge from dyadic feature embedding are placed along axes describing which rat is engaged in an interaction, and the perceived vigor or intensity of the interaction. **c**, Probability density of inter-animal distance during social recordings for the WT- WT and AMPH-PART pairs (top left) and fraction of time spent within a body length for each individual rat with and without amphetamine given an AMPH-WT interaction (top right). The change in HLJC usage for animals in the AMPH or PART condition as compared to their usage when part of a WT-WT dyad (bottom). **d**, Average difference in occupancy in each of the 156 low-level states for each rat depending on whether it has received amphetamine in a given session. Clusters are sorted by interaction type and distance between animals. At bottom, the average fold change between WT-social and AMPH-social conditions is shown, with dots indicating p < 0.05 (hierarchical mixed effects) Benjamini-Hochberg False Discovery Rate (BHFDR) correction for multiple comparisons (**Methods**). **e**, Four example social classes (LLJCs) demonstrating behavioral differences induced by amphetamine in the social context.

Inspecting and annotating the classes that resulted from mapping and clustering multi-animal trajectories revealed a range of highly specific stereotyped and recurring social interactions defined by shared spatiotemporal properties (**Fig. 4a, Supp. Video 4**). We found two axes of description particularly informative in interpreting these clusters: the strength of interaction and the symmetry of the interaction, i.e. how much each animal appeared to be engaged in or driving the interaction (**Fig. 4b**). Based on manual review of movies sampled randomly from the 156 low-level joint behavior clusters (LLJCs), each cluster was assigned a descriptive label (e.g., ‘*rat1 following rat2, body length apart*’ or ‘*both rearing, rat2 oriented towards rat1*’) (**Supp. Table 3**). We then assigned each of the 156 low-level joint behavior classes (LLJCs) to one of seven high-level joint behavior classes (HLJCs): *no interaction*, *lightly engaged*, *engaged*, *partner lightly engaged*, *light mutual interaction*, and *mutual interaction*. Together with the action clustering, this process yielded a rich annotated social dataset where at each timepoint an individual is assigned to both an individual (animal-autonomous LLAC) behavioral class and a joint (social interaction LLJC) class (**Supp.** Fig. 11**, Supp. Video 5**). Each of these low-level classes also belongs to one of a smaller set of high-level classes (HLACs and HLJCs, respectively), providing multiple levels of description.

Using the dyadic embedding, we were able to resolve several novel aspects of how acute amphetamine affects social behavior in rats (**Fig. 4b-e**). Past studies, relying on human annotation of broad behavioral categories, have been inconsistent, with some reporting no effect or effects specific to social play in juveniles^42,43^. Our method revealed statistically significant effects that were undetectable using coarse metrics, such as center-of-mass movement or the distance between interacting animals (**Fig. 3c**, **Fig. 4c**). For instance, variance in the inter-animal distance after drug administration increased, but the mean across animals remained unchanged from controls (**Fig. 2c**). However, when we compared the HLACs between amphetamine and control conditions, we could identify social behavioral types that were consistently modulated across the cohort (**Fig 4e**). Specifically, amphetamine increased social engagement and investigatory behaviors while undosed social partners showed less of these behaviors compared to control experiments in which neither animal had received amphetamine (.68 and -.46 fold change in engagement behaviors relative to baseline controls, for amphetamine dosed and undosed partners, respectively, **Fig. 4c *bottom***). Further, this finding belied a more complex set of changes at the finer descriptive level of LLJCs, with, for example, different classes of *engagement-related behaviors* not being uniformly upregulated. For instance, the LLJCs which capture an animal touching a rearing partner from the front or side exhibited a several-fold increase after amphetamine administration (2.7-fold and 2.2-fold, respectively, **Fig. 4d,e**). On the other hand, investigation and grooming of a partner from above exhibited the largest decrease (2.0-fold, **Fig. 4d,e**). Some HLJCs such as *mutual interaction* appeared unaffected as a whole, but we found that this was due to divergent responses across the LLJCs comprising this high-level class (10 mutual interaction LLJCs were significantly upregulated and 5 downregulated (**Fig. 4d, Supp. Table 4**), reiterating the value of our high-resolution, multilevel analysis.

To validate the general, multi-species utility of dyadic embedding, we applied the same framework to two common laboratory strains of mice. We compared BALB/c to C57BL/6, two inbred laboratory strains that have been proposed as models of low and high sociability, respectively^44,45^. Understanding the nature of social behavioral differences in these strains is particularly important for ASD research, as ASD mouse models developed on different backgrounds show pronounced phenotypic differences ^46,47^. When analyzing LLACs, we found behavior in social contexts to be highly consistent across animals of the same strain but observed extensive differences between strains, with C57BL/6 expressing a broader suite of higher velocity behaviors (**Supp.** Fig. 13). These differences persisted regardless of partner animal background, with LLAC usage remaining unchanged between same-strain and mixed-strain pairings. Dyadic embedding, however, revealed phenotypic differences missed when analyzing individual animals. In mixed-strain pairings, asymmetric engagement LLJCs were upregulated relative to same-strain pairings, with BALB/c mice approaching and inspecting C57BL/6 more often than BALB/c partners, agreeing with older results from three-chamber tests showing BALB/c mice prefer social novelty^48^. These results demonstrate how our system enables granular investigation of social behavior across species and genetic backgrounds.

### Tactograms capture body-wide patterns of social touch

In addition to behavioral classification, multi-animal pose estimation also facilitates the quantification of social touch, an important facet of social interaction often affected in autism^49^. Despite its clinical and behavioral relevance, social touch is relatively understudied, as doing so traditionally requires subjective and laborious manual annotation. To quantify social touch, we fit a previously developed volumetric model of a rat^50^ to the 3D keypoints of socially interacting animals and automatically identified points of contact between volumetric models (**Fig. 5a-c, Methods**). Each volumetric model represented the rat body surface as a deformable triangular mesh (6880 vertices) whose pose and shape were determined by the keypoint-derived rat body structure in each frame. To quantify body contacts, we counted the number and locations of mesh intersections between social partners for a given recording, producing a ‘tactogram’, or the distribution of observed social touches over the entire body over time (**Fig. 5d**). To aid visualization and analysis, we also computed coarser tactogram summaries by binning contact counts over the mesh faces comprising 7 gross body regions (**Fig. 5d**, *inset*). We validated social contact detection by comparing mesh-derived contacts to three human annotators on 1100 frames chosen randomly from timepoints where animals were closely interacting. This pipeline achieved human-level accuracy when validated against manual annotations (**Supp.** Fig. 14).

**Figure 5:**
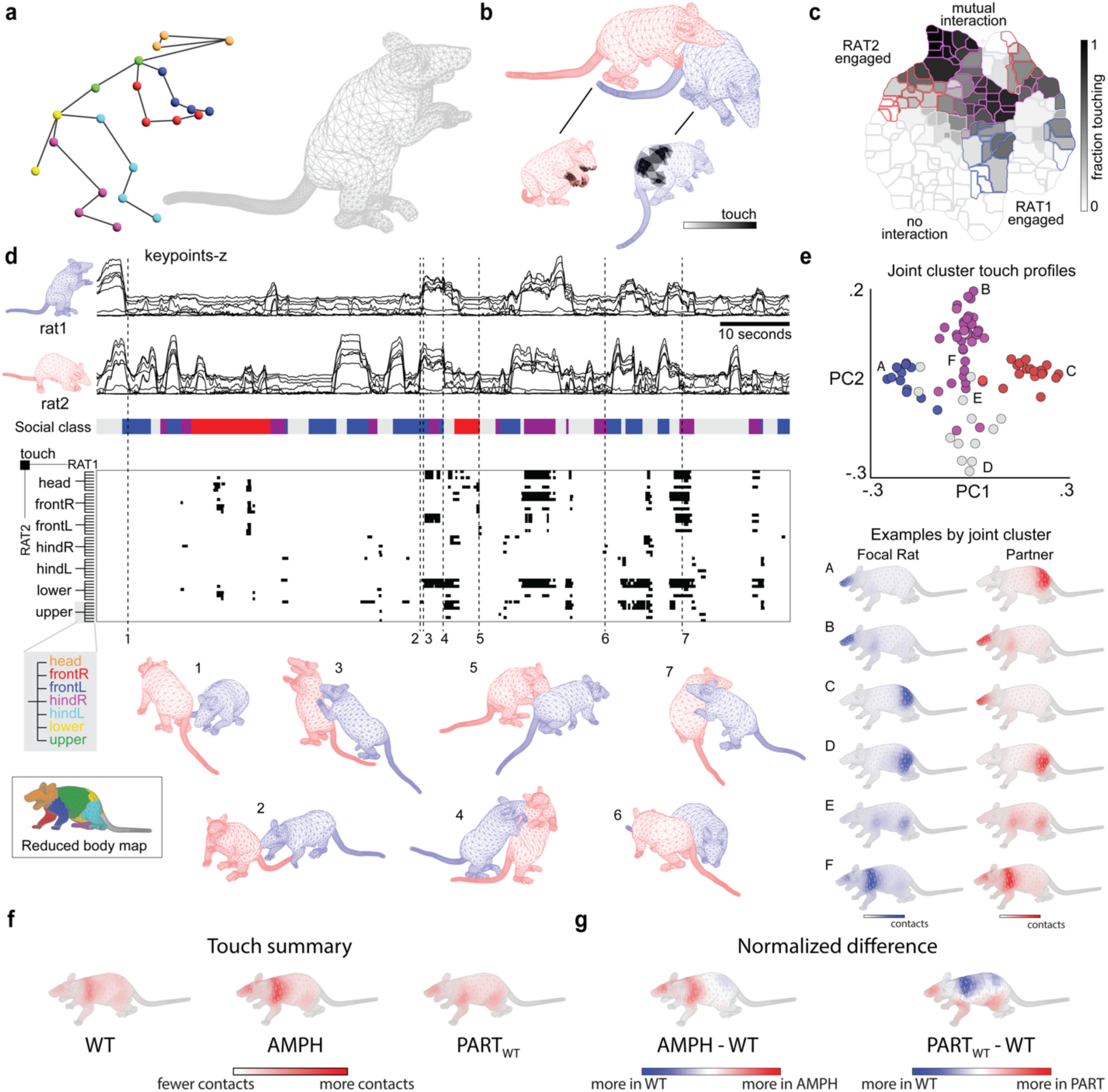
Volumetric body modeling from keypoints enables social touch profiling. **a**, 3D keypoints for each timepoint are used to fit a skin mesh model. **b**, Example of touch detection in a single frame, black fill indicates mesh faces that touch **c**, Fraction of time spent touching is shown for each high-level joint social behavior class (HLJC). **d**, Top, time series of keypoint z-axis coordinates for each animal in a representative recording and, underneath, the associated social class (HLJC) at each time point (blue, Rat 1 engaged; red, Rat 2 engaged; purple, mutual interaction; gray, no interaction). HLJCs were assigned using 3D keypoint analysis only (c.f. Fig. 4). Below, a raster plot showing contact events for the simplified tactogram where contacts are binned into seven major body regions (Reduced body map *inset*). The raster plot contains 49 rows, with each contiguous set of seven rows corresponding to one specific Rat 1 body region and the seven possible Rat 2 body regions it could contact. Body meshes are plotted for seven timepoints (dashed lines) when animals do not touch (1) and when they are touching in different configurations (2-7). Colors of example meshes denote animal identity, not HLJC. **e**, Scatter plot showing of PC scores for the mean touch profile within each low-level joint social behavior class (LLJC) for n = 396 paired recordings, 42 animals. Each dot represents a different LLJC, colored according to its associated HLJC (blue, Rat 1 engaged; red, Rat 2 engaged; purple, mutual interaction; gray, no interaction). All LLJCs and HLJC data groupings are derived from 3D keypoints only, not touch data, as described previously (c.f. Fig. 4). Below, example touch profiles are shown for the six LLJCs marked (A-F) in the PC scatter plot. Touch profiles are shown as densities over the mesh model. Hues of touch profiles denote animal identity, not HLJC. **f**, Touch profiles for WT, AMPH, and AMPH partner animals. **g**, Touch profile differences between the indicated experimental conditions.

We calculated tactograms for each LLJC to examine touch profiles across social motifs. Social touch patterns differed across LLJCs in both fraction of time spent touching and the patterns of body contact (**Fig. 5c,d**). Classes within the *mutual interaction* LLJC displayed a particularly diverse array of touch patterns, from nose-to-nose touching to widespread body contact, each of which could relay a distinct, ethologically-relevant social signal during interaction^4,51,52^. While touch profiles were diverse, they typically fell somewhere along two principal axes capturing (PC1) the symmetry of contact between social partners and (PC2) the anterior-posterior contact location (**Fig. 5e, Supp.** Fig. 15b,c). Fraction of time spent touching was also variable across LLJCs, ranging from short touches as animals brushed against each other, to prolonged contact when they were investigating (nose-to-body), allogrooming (head and upper limbs to nape), or huddling (sides of body or heads in contact). *Engagement* and *partner engagement* classes tended to cluster at opposite ends of the partner symmetry axis, with *mutual interaction* in between, serving as an independent validation of our HLJC annotations from touch profiling (cf **Fig. 4a**).

Tactograms capture dimensions of social interaction not resolved by 3D pose tracking alone, enabling deeper phenotyping in terms of a principal, yet historically understudied, social modality^30,53^. For instance, amphetamine had a spatially localized effect on social touch patterns, with dosed animals displaying a preference for touching the limbs and underside of their undosed social partners (**Fig. 5f**). These touch patterns could reflect the observed upregulation, in dosed animals, of social interaction types characterized by engagement with the front and underside of the undosed partner (**Fig. 4e, Supp. Table 3**).

### Multi-scale embedding reveals patterns of behavior across ASD models

Given that our methods identified novel amphetamine-induced changes to individual and social behavioral phenotypes, we asked whether they could also shed new light on how genes affect social behavior by profiling seven rat genetic models of autism: loss of function knockouts (KOs) of *ARID1B, CHD8, CNTNAP2, FMR1, GRIN2B, NRXN1 and SCN2A*^54^ (**Fig. 6a**). While these ASD models have been behaviorally phenotyped in mice, their social behavior has either not yet been studied (*ARID1B, CHD8, GRIN2B, SCN2A*) or not yet studied comprehensively (*CNTNAP2*^55^*, FMR1*^56^*, NRXN1*^57^) in rats, which are far more social animals^58,59^. Furthermore, reported mouse behavioral phenotypes have been inconsistent and often fail to recapitulate social effects observed in human autism^60–62^, casting doubt on the utility and validity of mice as ASD models. To probe for social phenotypes in rats and test their validity as models with consistent and relevant social deficits, we recorded and analyzed movies from 154.5 hours (27.8 million frames) of lone behavior and 304 hours (54.7 million frames; 109.4 million total 3D poses across both animals in a dyad) of social interaction across experimental groups (*ARID1B* N = 4 KOs, 4 WT littermates; *CHD8* N = 8,8; *CNTNAP* N=2,3; *FMR1* N = 5,3; *GRIN2B* N = 3,3; *NRXN1* N = 4,4; *SCN2A* N = 3,3). Social behavior was recorded in rat dyads in 30-minute sessions, with animals paired within experimental groups in an all-to-all round-robin design (**Supp. Table 1, Methods**).

**Figure 6:**
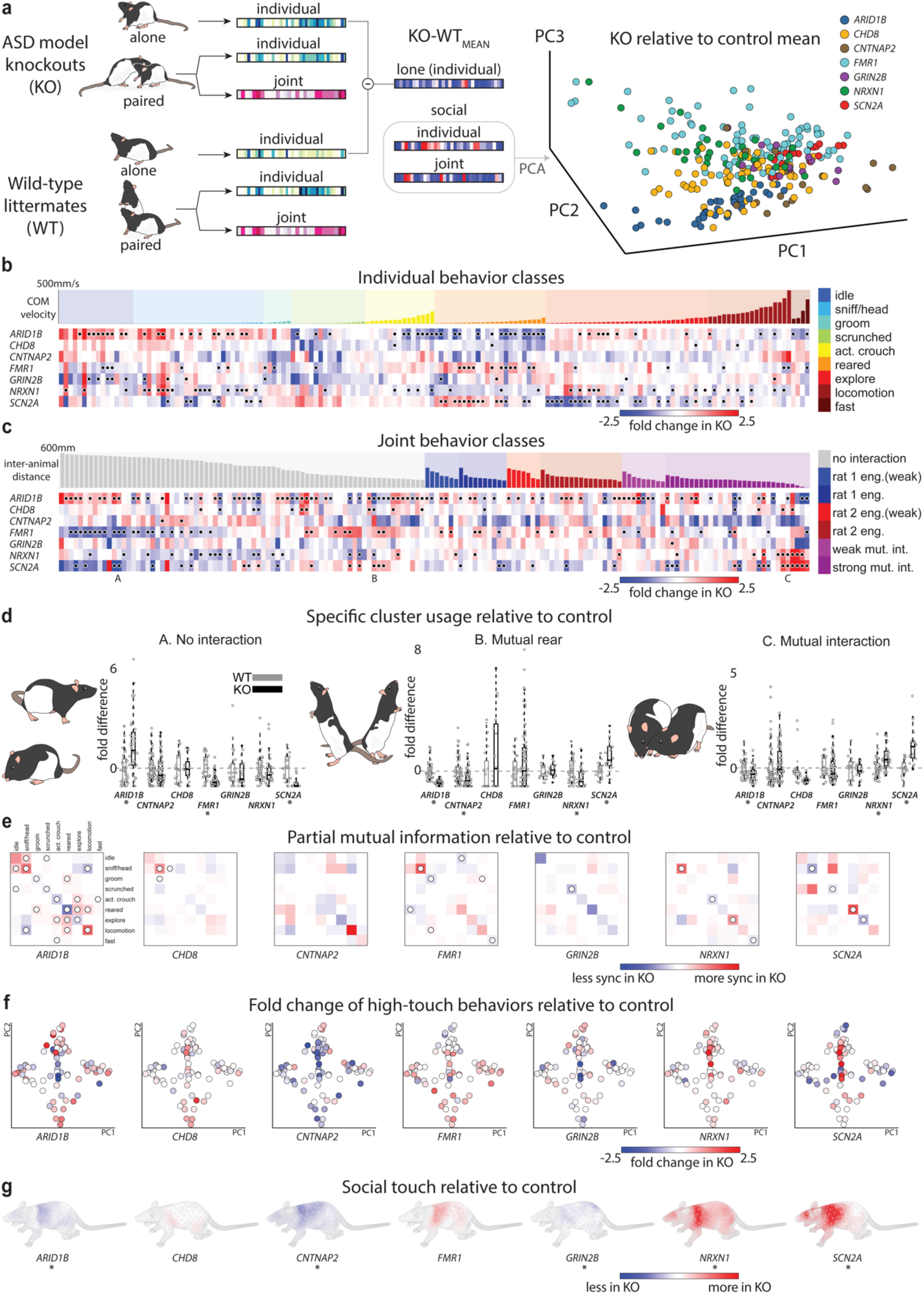
A comparison of social behavior across seven ASD model strains. **a**, Comparisons between knockout rats and their wildtype littermates are performed on the lone-individual, social-individual, and social-joint class occupancy vectors. Comparisons are always between animals from the same cohort to control for background, age, and environmental differences. Right, PCA of the combined individual and joint embedding vector difference from controls during social interaction reveals distinct clusters by knockout. **b**, Shifts in LLAC usage during social interaction are visualized for each knockout strain as a log fold change from the behavior in the respective wild-type littermate controls. Clusters that withstand a test of significance and FDR correction at p<.05 are marked with dots. **c**, Shifts in the knockout LLJC usage are visualized as in **b**. **d**, Individual occupancy shifts for each KO-KO social recording are plotted as a fold change from the respective wild-type controls for several example social clusters where at least two strains show a significant difference (indicated under strain name). Left, a non-interactive behavior where animals sit far apart was elevated in ARID1B knockouts and reduced in SCN2A knockouts in comparison to their respective controls. Middle, simultaneous rearing was elevated in SCN2A knockouts and reduced in ARID1B and NRXN1 models relative to their controls. Right, close head and body contact was increased in NRXN1 and SCN2A knockouts and reduced in ARID1B knockouts with respect to their controls. **e**, Partial mutual information was calculated for instantaneous usage of LLACs of paired animals in WT-WT and KO-KO pairings. The average difference between the two is shown, with dots indicating paired behaviors that withstand a test of significance and FDR correction at p<.05. **f**, Shifts in social clusters (as in c) are shown only for clusters where touching was detected for at least 10% of total frames, and on the same axes that delineate type of touch (as in Fig. 5e). **g**, Total touch over KO-KO experiments for each model strain was compared to the respective WT-WT control group, with asterisks indicating model strains where touch in at least one body part (as defined in Fig. 5d) was significantly different.

To map changes in social behavior across rat models of ASD, we quantified and cataloged behavior from tracked 3D kinematics using our multi-scale embedding approach. Additionally, we used our system to probe for and quantify irregularities in social touch, a core symptom of human ASD^63^ that, due to methodological limitations, is rarely studied in animal models^64^. Together, these revealed rich animal-autonomous and joint social behavioral repertoires that were largely consistent within genotypes but distinct across them, with KOs distinguishable from their littermates in their unique patterns of LLAC, LLJC, and touch usage (**Fig. 6, Supp.** Fig. 19**- 25**). Animal-autonomous behavioral profiling of interacting KO animals recapitulated several published mouse phenotypes while revealing novel characteristics not resolved by coarse metrics. For instance, while *SCN2A* KO rats exhibited an increase in rearing behaviors, as in mice^65^, we found that these increases were specific to low-velocity rearing types. Additionally, *SCN2A* KO rats were more likely than wild-types to perform synchronized rears (**Fig. 6c-e**). In contrast, the *ARID1B* KOs presented with a decrease in rearing, but particularly high-velocity rears, effects not previously reported. Using dyadic embedding, we found that paired animals across all groups engaged in many distinct interactions (e.g., allogrooming, chasing, mutual sniffing, or non-interactive idling) and overlapping but distinct subsets of these interactions were differentially impacted by genotype (**Fig. 6c-d, Supp. Table 3**). As each social motif is associated with a type of body contact (e.g., nose-to-nose sniffing results in head contact), differences in body-wide touch patterns covaried with strain-specific usage of social clusters (**Fig. 6f,g**).

The heterogeneity of autism was reflected in rat phenotypes. *ARID1B*, *CNTNAP2*, and *GRIN2B* KOs showed a tendency for reduced contact, driven by a reduction in close interactions. On the other hand, *NRXN1* and *SCN2A* KOs were highly interactive and touch-seeking, resulting in increased contact. *FMR1* KOs had a unique presentation, with robust differences from wild types in the usage of a majority of behavioral clusters where animals were not engaged in interaction. *CHD8* KOs were unique in having particularly high inter-animal variability, a hallmark of autism. These outcomes offer a choice of potential models for studying social deficits, sensory differences, variable penetrance, and non-convergent phenotypes^66^. Together, our analyses demonstrate an extension of ASD model phenotyping far beyond the status quo and provide a catalog of behaviors across models that can inform mechanistic models of social behavior and help formulate hypotheses about the neural mechanisms that underlie model-dependent differences.

## Discussion

We present s-DANNCE, a video-based technology for precise whole-body 3D tracking of interacting animals together with novel analytics for describing structure in social behavior. The platform captures behavior on multiple scales, spanning detailed, precise kinematics, individual and joint action expression, and body contact patterns, delivering a new and comprehensive approach for studying social interactions. To illustrate the power of our platform, we profiled interactions in rodents, uncovering novel effects of drugs, genes, and disease, including a spectrum of social changes across seven monogenic rat models of autism.

### s-DANNCE accelerates research and enables new lines of inquiry

By enabling more objective and sensitive studies of social behavior that scale to capture millions of behavioral events, s-DANNCE can accelerate research into the neural, genetic, developmental, and environmental underpinnings of the social brain and its disorders, as well as screen for novel therapeutics. Because our platform provides automated and quantitative measurements, which we showed were robust across animals, arenas, and sessions, it also lays a foundation for reproducible social behavioral studies across labs. To facilitate reproducibility and large-scale community collaborations, future work will need to explore the question of behavioral re-identification in other labs and explicitly establish standard behavioral definitions and quantitative metrics. To fully capture the complex and multi-modal nature of social behavior, s-DANNCE will need to integrate additional measurement modalities, including vocalizations^7^ and relevant physiological variables, such as heart rate and breathing rate. Similarly, to investigate neural underpinnings of social behavior, s-DANNCE must have the capability to synchronize with chronic neural recordings. Relationships between neural, physiological, and behavioral variables could then be probed via recent techniques that map neural activity onto 3D movement features^67,68^, or, more flexibly, via methods that associate brain and behavioral variables via multimodal fusion^69–71^.

### A better understanding of autism through animal models

Rodent models have delivered key mechanistic and therapeutic insights for many human diseases, but a glaring lack of face validity has muted their impact on autism research. At issue is whether rodent autism models exhibit social deficits, the defining symptom of human autism. In mice, the readouts from social assays have been inconsistent^3^, potentially reflecting the stark differences between mouse and human social repertoires. In contrast, rats readily and robustly express many behaviors affected in human ASD, including pro-sociality, cooperaction, and age-specific play^72,73^. High-resolution behavioral assessments will allow us to further probe the utility and face validity of both rat and mouse models. As evidence, we used our precise social phenotyping method to highlight specific social behavioral differences in genetic rat models of ASD and showed that several of the ASD genotypes are associated with strong and consistent deviations from their wild-type littermates. Importantly, we found evidence of ASD models with both increases and decreases in sociality, as well as genotype-specific differences in social variability across individuals, emphasizing the heterogeneous presentation of these models and the importance of deep phenotyping. In addition, we found divergent body-wide touch patterns across ASD strains, indicating that social deficits can present as both touch avoidance as well as touch-seeking behaviors, raising questions about whether social motivation can explain autism phenotypes^74^. Our approach also extended to mouse social behavior, and allowed us to identify consistent social differences between WT strains, including an inbred ASD model (BALB/c)^75^, and capturing shifts in mouse sociality induced by experimental context. Developing these methods further promises a standardized way to probe the landscape of social interactions, how this is shifted in ASD and other neurodevelopmental disorders, and which interventions may affect these shifts.

### Strategies to further enhance 3D social tracking

In the future, s-DANNCE’s 3D pose tracking precision and efficiency could be further refined by incorporating recent advances in machine learning and computer vision. For example, s- DANNCE could utilize temporal information, which promotes occlusion robustness in single-animal recordings^16,19,33,76^, although new temporal modeling strategies with reduced computational costs will likely be needed for multi-animal contexts. Tracking might also be refined by modeling animal pairs jointly, for instance by sharing network-extracted features between subjects^77–80^ or discouraging keypoint overlap^81,82^. A more comprehensive, physics-guided approach to tracking, for instance via integration of the biomechanical model supporting touch quantification^83,84^, might also better resolve the most occlusive behaviors, such as rapid tumbling.

### Further development of behavioral analyses and feature sets

To explore complex phenotypes with unknown types and degrees of behavioral presentation, we characterized interaction using a large set of 3D kinematic features spanning a wide range of possible behavior-defining movements. However, not all studies necessarily require the same resolution or feature sets. What is best for a specific social analysis will depend on a range of factors, including the behavioral types of interest, dataset size, and experimenter resources. For example, the relative simplicity of traditional approaches that measure a small number of social variables using either manual annotation or sensor-equipped behavioral chambers makes them an appealing choice when tracking a small number of known and pre-prescribed behaviors at coarser resolution. Features derived from fewer keypoints in reduced tracking configurations, such 2D poses or 3D poses without limbs, could also be used when coarser resolution is sufficient and when social gesturing or contact with limbs is not a primary research focus. Conversely, studies interested in subtle body language may require keypoint and feature sets expanded from what we present here. While we utilized an unsupervised wavelet-derived feature embedding to flexibly capture behaviors on different timescales, there are several alternative unsupervised behavioral identification approaches that are effective on single-animal pose datasets^85,86^. Our dataset offers a testbed for adapting these approaches, which might capture different aspects of social behavior, to 3D multi-animal data.

In the future, we envision bolstering our method’s flexibility and interpretability via support for a broader range of resolutions and the detection of predefined social behaviors. Predefined behavior detectors and their quantitative feature formulations could be shared with the community and applied to track these specific behaviors in new studies, including in real-time and in closed loop with behavior or neural manipulations. In parallel, unsupervised analysis will continue to drive discovery of new phenotypic descriptions using expanded datasets from new experimental contexts, postural tracking resolutions, and measurement modalities.

### A foundational repository for high-resolution behavioral analysis

To facilitate continued development of social behavioral quantification and analysis approaches, we are making s-DANNCE available to the community as an open-source python package and will publicly release our dataset and behavioral maps from over 500 million frames of 6-camera lone and social recordings collected over 80 animals. As a large repository of rich, high-resolution social behavior which includes wild-type rats and mice, rat amphetamine dosing experiments, and a comprehensive within-litter comparison of multiple rat ASD genotypes, we expect that this resource will foster discovery of new properties of neurodevelopmental disorders, stimulate new hypotheses of genotype-phenotype mechanisms testable in subsequent experiments, and serve as a reference behavioral atlas to which future quantitative behavioral studies can be compared.

## Methods

### Animals and husbandry

The care and experimental manipulations of all animals were reviewed and approved by Harvard University Faculty of Arts and Sciences Animal Care and Use Committee. We used 6 male Long-Evans rats (Charles-Rivers, strain 006), aged 9-14 weeks for the wild-type and amphetamine dosing experiments. Monogenic rat autism models were ordered from The Medical College of Wisconsin in litter- and age-matched cohorts of 6-8 male rats and recorded between 12 and 20 weeks of age. Ear punches were genotyped after birth and experimenters were not blinded to genotype. All knockout animals and littermates were genotyped from tail clippings after all experiments were concluded. Animals were kept on a normal 12/12 light/dark cycle at a temperature of 22°C and humidity of 30-70% and were housed in ventilated cages with ad libitum food and water. Animals were housed with littermates and isolated at least 48 hours prior to social experiments.

### Recording apparatus

All recordings were performed in a custom-built elevated cylindrical arena of 1 meter diameter, where animals could move freely. The circular arena base was water-cut from green HDPE (high-density polyethylene) and the arena wall was constructed from 1mm-thick 60 cm-tall clear polycarbonate sheet to contain animals and prevent escape. The arena was illuminated from above by two white LED arrays (Genaray SP-E-500B, Impact LS-6B stands) and surrounded by a commercial fabric green screen for background consistency and contrast to animals. Six high-speed 2MP Basler Ace-2 Basic cameras equipped with 8mm lenses were synchronized with an Arduino IDE 50 Hz hardware trigger called using campy (https://github.com/ksseverson57/campy). A windows PC with 64 GB of RAM equipped with two GPUs (NVIDIA Quadro P4000, NVIDIA GeForce GTX 1660 SUPER, and NVIDIA Quadro RTX 4000 were tested) enabled acquisition from six cameras simultaneously.

### Camera calibration

Intrinsic calibration was performed using the built-in matlab Camera Calibration app. Extrinsic calibration was performed with openCV. Calibration parameters were saved in each experiment folder. The entire arena was surrounded with six cameras on tripods which were calibrated prior to experimental recordings. Each day, the current calibration parameters were checked using Label3D (https://github.com/diegoaldarondo/Label3D) prior to performing any recordings. In order to check calibration, a short video with a stationary object was captured from all six cameras using campy (https://github.com/ksseverson57/campy) and the six views from a single frame were loaded into Label3D. Triangulation from two views was performed for a single point on the object and, if the calibration was still accurate, would correctly identify the chosen point from all remaining views. The calibration was stable enough such that videos could be recorded using the same calibration for weeks, and a new calibration was saved whenever a camera was moved. The pixel resolution is approximately 0.5 mm per pixel on average across camera views.

### Behavioral recordings

#### Experimental protocol details

Before social recordings, all rats were recorded individually in the arena for five consecutive days (except in the case of the *FMR1* cohort, where there were only 3 lone recordings per rat). Animals were then separated into individual housing and kept isolated for at least 48 hours before the start of social recordings. All experiments were completed between 9am and 6pm, however each individual cohort was recorded during the same four-hour time window across days. The arena was cleaned with 70% ethanol each night after recordings were completed and left to dry. Between recordings, a small amount of 70% ethanol was used to wipe the floor and walls of the arena and allowed to dry before the next recording. Rats were randomly paired in an all-to-all round robin format within each experimental cohort. After the complete round robin, additional recordings were performed of rats in random pairings (see **Supp. Table 1** for a list of each pairing for each rat in every cohort). Animals were all housed and maintained in a facility where they were given food and water ad libitum and monitored by veterinary staff. Cages were cleaned and bedding was replaced each week or as needed. Lights were on from 6am to 6pm.

#### Lone Recordings

Animals were placed directly from the home cage into the acquisition rig for 30 minutes (1800 seconds) on each of five consecutive days with recording starting immediately. The arena was ethanol-wiped and allowed to evaporate dry between recording sessions. All animals from the same cohort were recorded across the same days and each animal was recorded within a consistent 4-hour window across days.

#### Social Recordings

Prior to social recordings, all animals underwent lone recordings in the acquisition rig as described above. Animals were then separated into individual lone cages for at least 48 hours. Animals were recorded in all possible pairs across the given cohort (with their littermates and possibly other animals from the age-matched cohort in cases of small litters). For each social recording, the selected pair of animals were marked with diluted food coloring in order to simplify center-of-mass (COM) tracking during post processing steps. Animals were placed in the arena in quick succession with allowance for less than 5 seconds of interaction before recording began. Each animal was only recorded in one social session per day except in the case of *CNTNAP2* and *FMR1* cohorts, where multiple recordings could occur in a single day.

#### Pharmacology Recordings

Prior to lone and social experiments with amphetamine administration, animals were briefly anesthetized using isoflurane, weighed, and injected with 1.25mg/kg amphetamine. After waiting 20 minutes for recovery and acclimation to the injection animals were recorded as previously described. Animals were never dosed with amphetamine or paired with an amphetamine partner in consecutive recordings.

#### Long Evans rat cohort

We ordered six Long Evans males from Charles River (Strain Code: 006) to be used for wild-type and amphetamine behavioral recordings. Animals were group-housed upon arrival to our animal facilities.

#### ASD model rat cohorts

We ordered *Arid1b*, *Chd8*, *cntnap2*, *Grin2b*, *Fmr1*, *Nrxn1* and *Scn2a* knockouts (only heterozygous except for cntnap2 and Fmr1) from the Medical College of Wisconsin (MCW) which produced and maintains these strains in order to facilitate study of rat models of autism (https://www.sfari.org/resource/rat-models/). All knockouts are maintained on a Long-Evans background. Animals arrived housed in groups of 3 or 4 and were allowed to acclimate in home cages for at least 72 hours after arrival. Animals were handled after arrival for weighing and exposure to experimenters. The first set of behavioral recordings for each animal was a set of daily 30-minute lone recordings for five consecutive days (3 in the case of *FMR1*) in the behavioral arena. Each day, animals were removed from the home-cage and placed in the rig by hand after which the experimenter left the room until the recording was complete. Animals were given no prior exposures to the arena for habituation, opting instead to capture the habituation across days. Following the lone recordings, animals were separated into lone housing in home cages and after at least 48 hours social recordings would commence. Each animal was only exposed to one social interaction recording per day (except in the case of *FMR1*, where multiple recordings could be performed on any given day) and all possible pairings within a cohort were completed before any repeated pairings.

#### Rat triad recordings

To test the extension of s-DANNCE to >2 animals, we recorded six additional movies of tryads from the CHD8 group several days after all lone and dyadic interaction recordings were complete. The three animals were marked using small patches of different colors of food coloring (blue, green, and red) prior to recording and placed into the arena from separate cages immediately prior to the start of recording.

#### Bedding recordings

To test the extension of s-DANNCE to an arena with bedding, we recorded six additional movies of pairs of Long Evans female rats for 30 minutes each. The animals were marked using small patches of blue and red food coloring prior to recording and placed into the arena, which contained a layer of bedding approximately 1 cm thick.

#### Mouse cohorts

While we developed our method in rats to study their complex social behavior, mice are commonly used in studies of social behavior and as models for ASD symptoms. We compared the lone and social behavior of males from two commonly used lab strains, C57BL/6 and BALB/c, as well as recorded mixed social dyads composed of one mouse from each strain.BALB/c mice are an albino strain often used for immunology research, and have been described as a model of low sociability^75,87^.

We obtained eight C57BL/6 (Strain Code: 027) and eight BALB/c (Strain Code: 028) male mice from Charles River and allowed mice to remain group-housed for the duration of behavioral experiments. Mice were housed in cages of four same-strain animals. Each mouse underwent two days of lone recording (10 minute experiments) in the behavioral arena after acclimation to the animal facility. Following lone recordings, animals were recorded in pairs for ten minutes each, with each mouse appearing in several recordings in a given day. The white BALB/c animals were marked with a small patch of food coloring on their sides as with the rats to track animal identity. The black C57BL/6 animals required lightening of the fur so half of the animals had small patches of fur on their sides bleached using commercial hair bleach. Bleaching was performed by swabbing a small amount of bleach mixture onto the fur while gently holding the mouse. The bleach was then wiped off and mice were returned to their home cages. This process was repeated across two days for a stronger bleaching effect.

### Estimating 2D top-down and bottom-up occlusion

Three sets of social recordings (“SCN2A”) and the corresponding s-DANNCE predictions were used for estimating the average occlusion from the top-down and bottom-up view. All 3D points were rotated into the camera coordinate system of the virtual top-down/bottom-up camera and were sorted by their distances to the camera center in descending/ascending order within each separate frame. To account for occlusion by body soft tissue, we made a simplification that a keypoint is considered as occluded if it falls within the occlusion region formed by all keypoints above it in a top-down camera view and vice versa for a bottom-up camera view (**Supp.** Fig. 1a). We qualitatively and quantitatively examined the changes in occlusion rates with different occlusion thresholds around each keypoint (5, 10, 15, 20, 25 mm) (**Supp.** Fig. 1b-d). For all numbers reported in **Results,** we adopted a threshold of 15 mm.

### Posture annotation

To acquire 3D posture labels for model training, one human labeler (‘Labeler 1’, **Fig. 2f**) selectively annotated n = 423 unpaired 3D poses from n = 7 social recordings of CNTNAP rats and annotated one social recording held out from the training set solely for model performance benchmarking purpose (n = 1373 frames, both animals, **Fig. 2i, j**). To validate the model’s generalizability to different rat strains, another social recording of rats with FragileX syndrome was similarly annotated (n = 1058 frames, both animals, **Supp.** Fig. 5e-h). For the inter- and intra-observer error analysis (**Fig. 2f-h**), n = 40 frames with close interaction were evenly drawn from two SCN2A social recordings, where three human labelers annotated both animals in these frames for two rounds with permuted frame orders. The total number of 3D poses annotated by each human labeler was n = 160. All 3D pose annotation was performed using Label3D which triangulates multi-view 2D annotations into 3D. We annotated with a skeleton consisting of 23 body keypoints, as coarsely grouped into 4 body part regions: head (*Snout, EarL, EarR, SpineF*), trunk (*SpineM, SpineL, TailBase, ShoulderL, ShoulderR, HipL, HipR*), forelimbs (*ElbowL, WristL, HandL,ElbowR, WristR, HandR*) and forelimbs (*KneeL, AnkleL, FootL, KneeR, AnkleR, FootR*).

### Multi-animal 3D centroid localization

We tracked 3D kinematics in interacting animals by first training a multi-instance COM network to roughly locate different individuals and then applying s-DANNCE to infer a keypoint-based skeleton per instance in each frame. Specifically, we construct volumetric representations from multi-camera images, which are anchored at each animal’s estimated 3D centroid in the world coordinate system for maximizing the estimation resolution. To determine the animals’ center-of-mass (COM) positions, we used a 2D U-Net architecture similar to Dunn et al. ^17^. We configured and trained different COM networks for lone and social recordings by setting the number of output channels as the number of animals present in the scene. For training, model outputs were optimized against 2D Gaussian-shaped heat maps generated from human annotations of COM positions using a mean-squared Euclidean loss. The 2D COM positions were extracted as the maximally activated position in the 2D heat maps during inference in each camera view and the corresponding 3D COM position was reconstructed as the median across all possible camera pairs.

For each set of experiments, for example, all lone or social recordings from a specific cohort of animals, we trained a separate COM network. We labeled approximately 200-500 frames for training each network, making sure to include samples from each animal that appeared in the set of recordings. For social recordings, training frames always included multiple COM labels which were ordered by color (blue, green, red) in order to preserve animal identity. If different sets of colors were used in the same dataset, a separate COM network would be trained for each color pairing.

### Graph neural network pose refinement module

For the first stage, we adopted a 3D encoder-decoder architecture to process the 3D volumetric inputs and output the initial estimation of probability distributions for each body marker. For each 3D heatmap associated with a specific keypoint, we performed a differentiable integration trick^88^ to locate the maximally activated location, or center of mass of this 3D volume, as the predicted position of that keypoint. With the initial 3D poses obtained from the regression stage, we defined an undirected graph *g = (v, ε)* where the nodes were associated with the relative voxel coordinates of body markers *v* = {*ν*_1_,*ν*_1_, …*ν_N_*} and the edges *ε* = {*e_1_*, *e_2_*,…,*e_M_*} linked *M* pairs of keypoints based on anatomical connectivity of the animal. Accordingly, we defined a *N×N* adjacency matrix *A* where 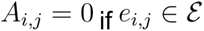 else 1. We refer to this graphical representation as a skeletal graph.

We sampled using bilinear interpolation at the maximal activation positions from features maps of the three deconvolution layers. The multi-scale features were concatenated with each keypoint’s predicted positions to form each graph node’s input features. We then adopted a graph neural network (GNN) to further refine the initial marker localization results. The basic building block of the GNN is a graph convolution module (**Supp.** Fig. 4a). Given a generalized graph based input defined as above, each graph convolution block outputs 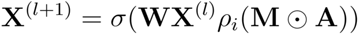 where M acts as an additional weighting parameter that adjusts the contribution of neighboring nodes. The GNN module generates spatial offsets to the initial marker positions in the voxel space. The final marker locations in the world coordinate system were obtained after being scaled by the voxel resolution and translated by the world coordinates of the volume center.

### Physical body constraint

We leveraged prior knowledge of rodent body anatomy to constrain the posture solution space. Given all sets of connected markers *ν_i_* and *ν_j_*, the lower and upper bound of their expected spatial distance [*d^l^*, *d^u^*], we defined the corresponding anatomical plausible range for the lengths of 23 body segments. For better scalability to variably sized animals within and across different rat strains, we reformulated the bone constraints from absolute distances to relative ratios across different segments. In practice, we scaled each segment with the distance between the animal’s left ear and right ear landmark. The expected ratios of body parts were pre-computed from ground-truth pose labels and imposed on both male and female individuals as regularization. We imposed body constraints in the form of L2 loss during training.

### Semi-supervised learning scheme

The previously described body constraint loss was applied on a mixture of labeled and unlabeled timestamps (n = 423 labeled, n = 910 unlabeled). To alleviate the tracked marker ambiguity in occluded social scenarios, we further adopted a strong data augmentation scheme that captured the spatial invariability in 3D metric space. During training, we formed each training data batch with augmented copies of one single input volume and imposed batch-wise consistency over all marker location predictions, using a L2 loss *L_c_*.

The overall optimization objective is given by 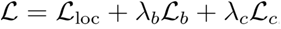, where we define *L_loc_* as L1 distance between model keypoint predictions and ground truth 3D positions, if available for the current frames, and λ_b_, λ_c_ are hyperparameters that balance different loss components during training, which are subject to tuning in finding the best-performing combinations.

### Model implementation and training procedure

The s-DANNCE framework is implemented in PyTorch. The GitHub repository is made available at https://github.com/tqxli/sdannce with codes and detailed instructions for model training, inference and visualization of the tracking results.

For estimating the animals’ 2D centroids as the prior step for pose estimation, we trained 2D U-Net COM networks with skip connections. The numbers of channels used in each layer are [32, 32, 64, 64, 128, 128, 256, 256, 512, 512, 256, 256, 128, 128, 64, 64, 32, 32, c = the number of animals present in the scene], where c = 1, 2 or 3. We did not explore more sophisticated tracking strategies as we only focused on animal interactions in dyads and triplets, but anticipate that this step can be conveniently replaced with other existing centroid/identity tracking methods.

For estimating the 3D poses in rats using s-DANNCE, we adopted the same 3D U-Net architecture as Dunn et al. except that we halved the numbers of feature channels in each layer to reduce the model memory footprint. We pretrained the 3D encoder-decoder (i.e., baseline DANNCE) on a subset of the RAT7M motion capture dataset (n = 88194) for 20 epochs with an initial learning rate of 0.0001, which was decayed at epoch 10 and 15 by a factor of 10. To compensate for the fine-scale body markers at limb ends missing from the RAT7M dataset, we fine-tuned the 3D encoder-decoder backbone with n = 1009 labels with 23 body markers selected from lone rat recordings. The training was done for 450 epochs with a constant learning rate of 0.0001. Note that this adaptation step can be removed without significantly affecting the final performance except for slower convergence. Lastly, we included the GNN module (**Supp.** Fig. 4a) and trained the full s-DANNCE model using n = 423 3D poses collected from social recordings, together with n = 910 unlabeled frames for semi-supervised learning as previously described. The model was trained for 100 epochs, with an initial learning rate of 0.0001 decayed at epoch 50, 70 and 80 with a factor of 5, and optimized using the objective function described above, where hyperparameters λ_b_ = 0.5 and λ_c_ = 0.1. The comparison baseline DANNCE model was trained following the same schedule except for the differences in architecture and optimization objective. All 3D rat pose predictions used for analyses in this paper were obtained from the same s-DANNCE model, including all lone, dyad and triplet recordings.

For generalizing to paired mice and to rats in arenas with bedding, we respectively finetuned the primary rat s-DANNCE model using n = 111 3D mouse pose labels and n = 211 3D rat labels in bedding frames, combining with equal amount of unlabeled samples randomly sampled from frames with inter-animal distances no greater than 120 mm. We did not freeze model parameters during the finetuning and retained the s-DANNCE optimization objective described in section “Semi-supervised learning scheme”. Each model was trained for 40 epochs, with an initial learning rate of 0.0001 decayed at epoch 20 and 35 with a factor of 5.

### Comparison with Han et al. 2024

Han et al. described a multi-animal 2D tracking method, which performs 2D pose estimation (DeepLabCut, or DLC) on masked frames from video instance segmentation. To obtain the instance masks required by training and inference of the image masking method, a human labeler annotated n = 3912 identity-preserving instance masks in n = 1980 frames (severely occluded instances were not annotated), including the entire dyad dataset used for training the baseline DANNCE and s-DANNCE model. These ground-truth (GT) instance masks and corresponding frames were used to train a 2D DLC pose estimation model used for quantitative analyses in **Supp.** Fig. 7. The DLC model was trained following the default training settings for a ResNet50 backbone for 50 epochs until convergence. The 2D pose predictions yielded by the DLC model were triangulated into 3D after taking the median among all possible camera pairings for each timestamp. For the evaluation, we separately evaluated the image masking method’s performance (1) with “GT masks”, where the same human annotator annotated animal masks in all test frames, and (2) with “predicted masks”, where the training GT masks were used to train a Roboflow 3.0 Instance Segmentation model on the Roboflow platform (starting from the v12 public checkpoint pretrained on MS COCO) and the instance segmentation model was used to annotate the test set.

### Individual (animal-autonomous) behavioral mapping

The inferred 3D landmark position trajectories for each recording were used to create a behavioral map which could be used to assign a behavioral label at each time point for each animal across all recordings. We used methods described in several previous works^34,35^ and altered preprocessing, feature selection, and clustering metrics for better performance on the specific data used. All code for deriving the classifications used throughout the results, as well as other examples, will be shared in the s-DANNCE GitHub repository upon publication. Here we briefly describe the method to transform a large dataset of kinematic recordings to behavioral timeseries or ethograms.

All keypoint data were inferred from movies taken at 50 Hz using the same base skeleton (23 keypoints, detailed information saved in ‘skeletons/rat23.mat’) using s-DANNCE, and preprocessed by median filtering with a 3-frame window followed by a smoothing with a 3-frame window in order to reduce any tracking jitter. First, to normalize for the varying animal sizes across cohorts, the size of each animal was estimated by sampling the distance between two virtual markers (the snout and the tail base) and finding the 97.5th percentile. Inferred points were scaled by this scalar value. Each time point was represented as an all-to-all distance of 23 body marker positions, producing a rotationally and translationally invariant description of the posture of the animal. We performed PCA decomposition on this representation across all recordings to find the top 15 postural principal components which represent the instantaneous posture. To incorporate multi-scale temporal information, we computed the power of each of 25 dyadically spaced frequencies for each of these postural projections. The final representative high-dimensional dataset for each movie incorporated a wavelet decomposition of the PCs (15 modes, 25 frequencies each) as previously described as well as a scaled representation of the height of each joint relative to the arena floor (23 keypoints, scaled by 1/10), and speed for each keypoint (23 keypoints, scaled by 1/2). We found that introducing these features was especially helpful in distinguishing rears (as egocentric alignment removed raw height data) and small movements. The final high-dimensional representation consisted of a 421-dimensional vector at each timepoint for each individual animal, whether in a lone or social context.

In order to enable behavioral comparisons across all conditions and experimental cohorts, a subset of high-dimensional postural-temporal representations was used to generate a behavioral map which was then used to re-embed each kinematic trajectory. We sampled frames uniformly in time and performed a t-distributed stochastic neighbor embedding (tSNE) for each movie and sampled temples from these initial embeddings to build the final training subspace. The final set of training samples (∼45,000 samples) was embedded into a 2D space using tSNE, and each frame from all movies was then embedded into this space to create a density map of behavior across all experiments. Applying a watershed transformation to this space (sigma = 1) resulted in 163 spatial clusters along this 2D density which represent fine-grained behavioral clusters which we refer to as LLACs (low-level action clusters). These clusters were further grouped into nine coarse categories which we refer to as HLACs or high level action clusters (*idle*, *sniff*, *groom*, *scrunched*, *reared*, *active crouch*, *explore*, *locomotion*, and *fast*). The fine- and coarse-grained cluster labels were applied to behavioral recordings taken from both lone and paired animals.

### Behavioral synchrony calculation

Synchrony analysis was performed by considering the HLACs assigned simultaneously to paired animals across each recording as described in ^11^. To identify co-occurrence patterns of HLACs in interacting animals, we calculated a 9x9 synchrony matrix (S) for each paired time-series using the equation

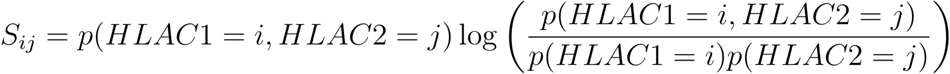

where HLAC1 and HLAC2 refer to the fractional occupancies of each of the nine HLACs (high-level action clusters) for animal 1 and animal 2. Each entry in S captures how likely the pair of behaviors are to occur together above chance. The underlying behavioral distributions were calculated independently for each recording. We report the average for S across experimental groups (**Fig. 3f, Supp.** Figs. 18-25).

### Social behavior mapping and perspective embedding

We extended the previously described methods for unsupervised individual behavioral mapping to produce behavioral clusters capturing the postures of two interacting animals, where features capture the relative posture and dynamics. Several changes were necessary to perform multi-animal social embedding and capture social motifs. First, since the identity of an animal matters within a social context, each social cluster is from the perspective of one of the animals. In practice, this means that features are calculated twice for each interaction recording, with the ordering of keypoints flipped for capturing the perspective of the second rat. We used fewer principal components (6) to describe the shared posture and positioning of interacting animals than in the single animal case, as the emphasis was in capturing stereotyped instances of shared movement instead of a combinatorial description of the movements of each animal. The time scale used for the joint wavelet decomposition was also slower in order to accommodate slower and less precisely coupled movements than in the case of single-animal limb coordination. Shared and individual-animal features were combined to produce the final high-dimensional representation. This combination of features capture aspects of the posture and movement of each animal and emphasizes the shared measures such as distance and orientation between animals as well as the dynamics of shared movement.

To perform the joint embedding, wee first calculated the all-to-all distances between selected 3D keypoints (*snout*, *spineF*, *spineM*, *spineL*, *tailBase*, *shoulderL*, *shoulderR*, *handL*, *shoulderR*, *handR*, *hipL*, *footL*, *hipR*, *footR*) on each animal and performed online-PCA to determine six basis vectors to capture the shared postural representation across all movies. Wavelet decomposition at dyadically spaced frequencies ranging from .2-5Hz was performed on the resulting projections obtained from representation for each social recording. Additional features including distances between specific body parts and angles of heading deflection between animals in 3D were derived from the inferred keypoints and added to the high-dimensional time-series describing the shared dynamics.

In order to sample across all contexts and balance the final embedding in consideration for rare motifs, templates were retrieved from each social recording and pooled to form a training set for embedding using t-SNE, as described for the animal-autonomous mapping. The resulting training set and 2D t-SNE map were then used to re-embed each recording, resulting in a final representation that was clustered using the watershed algorithm. The corresponding embedded values were used to assign each frame a cluster number. These social clusters capture possible animal configurations incorporating not only static features such as distance and orientation, but also properties of these shared features across a span of time scales. Each of the resulting 156 clusters (five clusters could be eliminated because of a lack of samples) was assigned a low-level description based on viewing randomly selected snippets of frames which were assigned the given label (**Supp. Video 4**). The movies contained both animals but were described from the perspective of the focal animal, giving rise to categories that aim to capture how engaged the focal rat and partner are, leading to a description of how engaged each animal is and how symmetric this engagement appears to the human eye. While many clusters (76 of 156 total) were classified as ‘No Interaction’, these clusters still contain valuable information about the interaction between animals and range from low-level descriptions like *‘laying on opposite sides of the arena with little to no activity*’ to ‘*both performing active rears while less than one animal- length apart*’. While there does not appear to be active inter-animal engagement in case of the latter description, this cluster captures a synchronized movement.

Importantly, each animal within a social pairing is assigned its own social cluster label at every frame, as the social embedding retains the identity and perspective of the given animal. For example, one animal may correspond to a cluster where, from that animal’s perspective, it is “chasing” the other animal, whereas the partner will instead correspond to a cluster that captures “being chased”. These ‘perspectives’ can be resolved to a single interaction-level annotation based on an agreement matrix derived from the human-annotated descriptions. This intermediate description can be used to evaluate how much one-sided or mutual interaction occurs throughout a social recording and how interaction progresses over time (**Supp. Video 5**).

### Manual annotation of clusters

Two human annotators independently viewed movies of reconstructed 3D skeletons randomly sampled from each behavioral cluster from individual and joint mapping. Sampled clips were picked at random from all experimental movies, from time points where the given label was assigned for at least 25 subsequent frames or 500ms of consecutive kinematics. For individual (animal-autonomous) clustering, each cluster was assigned a short description and a coarse category (**Supp. Table 2, Supp. Movie 3**). For joint clustering, each cluster was assigned a description and the interaction was given a coarse social category (*no interaction*, *lightly engaged*, *engaged*, *partner lightly engaged*, *light mutual interaction*, and *mutual interaction*) intended to capture the level of asymmetry and intensity of the sample movies (**Supp. Table 3, Supp. Movie 3**). Any clusters where annotators were in disagreement were reviewed together until a consensus was reached to produce the final set of coarse behavioral cluster descriptions presented here.

### Estimating social contact with deformable mesh models

To estimate points of contact between socially interacting rats, we registered a biomechanical model paired with a deformable mesh model to s-DANNCE keypoints and quantified intersections between meshes. Here, we will describe the biomechanical model and accompanying deformable mesh model, the registration process, and the algorithms used to determine contacts.

#### Biomechanical model

We used a biomechanical model paired with a deformable mesh model as a tool to estimate the volumes of interacting animals at each frame by registering the biomechanical model to the keypoints inferred with s-DANNCE. The biomechanical model of the rat was developed as part of a previous study^50^ to resemble the bone length and body mass distribution of Long Evans rats in the laboratory and work with the MuJoCo physics simulator. The pose of the biomechanical model has 74 degrees of freedom (DoF). These include the global Cartesian position of the model’s root (3 DoF), the quaternion defining the orientation of the root (4 DoF), and a set of transformations denoting the joint angles of each body part relative to its parent in an acyclic tree that denotes the relationships between connected body parts (67 DoF).

To account for differences in animal sizes, we isometrically scaled the biomechanical model according to the segment lengths estimated with s-DANNCE. For every session, we calculated the median segment length for each animal and each session, *seg_as_*, and computed the median segment length across all sessions, *seg_med_*. The scaling factor for a particular animal and session, *s_as_*, was then defined as the ratio 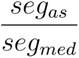.

#### Deformable Mesh

The biomechanical model described above includes a deformable mesh model that was created to approximate the shape of a rat. The deformable mesh model is implemented using the MuJoCo skin asset. Briefly, the deformable mesh is composed of a set of 6880 vertices connected to form a set of triangular 10752 faces that together approximate the shape of a rat. The position of each vertex and the orientation of its normal is influenced by the positions of one or more bones in the biomechanical model. Each bone contributes to the position of a set of vertices with a weight that specifies how much the position and orientation of the bone contribute to the position of the vertices. The 3D position of each vertex is calculated by weighting the influence of its corresponding bones. Normal vectors for the mesh are computed from the 3D vertex positions and the faces derived from connections of vertices.

To better approximate the volume of animals of different sizes, we specify an inflation parameter for each animal which expands or contracts the deformable mesh along the normal vectors by a given distance. Similar to the scaling of the biomechanical model, we used the distribution of bone segment lengths estimated from s-DANNCE to determine the inflation parameter. The inflation parameter for a particular animal and session, *i_as_*, was calculated as follows

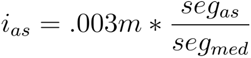

The value of .003 m was determined through visual inspection by overlaying the deformable mesh models of different inflation parameters atop the multicamera images of the animal with the median segment length. The inflation parameter allowed us to scale the size of the deformable mesh model to account for variability in the animal sizes, or growth that occurred throughout the collection of the dataset.

#### Registration

Our biomechanical model was designed to approximate the general skeletal structure of the rat, and not skin, fat, muscle or cartilaginous tissue. This presented a problem when trying to register the biomechanical model to s-DANNCE keypoints, as several keypoints correspond to bodily landmarks that were not present in our biomechanical model. To address this problem and register the biomechanical model to the s-DANNCE keypoints, we used a custom implementation of simultaneous tracking and calibration (STAC)^89^. STAC iteratively optimizes two quantities in an alternating fashion to calculate the biomechanical model poses that best explain the keypoint positions: a set of learned offsets that relate points on the biomechanical model to s-DANNCE keypoints (m-phase) and the joint angles of the biomechanical model (q-phase) (**Supp.** Fig. 14).

In the m-phase, STAC optimizes a set of offsets that each relate a reference point on the biomechanical model to an s-DANNCE keypoint. Specifically, the m-phase uses L-BFGS-B to minimize the mean squared error between s-DANNCE keypoints and ‘fictive keypoints’ defined by applying the set of offsets to the set of reference points. The reference point for a given s- DANNCE keypoint was defined as the location of the parent of the body closest to the keypoint. The offset was defined as a 3D vector extending from the reference point in the reference frame of the parent body. This optimization occurs over a small training dataset that will be described in greater detail below. At each step of the optimization, we adjust the set of offsets by an epsilon step, compute the new fictive keypoints using MuJoCo’s forward kinematics, and compute the empirical gradient to determine the direction of the offset update.

In the q-phase, STAC optimizes the joint angles (expressed as a set of quaternions) via least squares optimization to minimize the mean squared error between the s-DANNCE keypoints and the fictive keypoints. At each step of the optimization, we adjusted the joint angles by an epsilon step, computed the new fictive keypoints using MuJoCo’s forward kinematics, and computed the empirical gradient to determine the direction of joint angle updates. Importantly, the offsets were fixed during the q-phase, enabling the optimization of joint angles for a given set of offsets and frames. We applied this procedure to each frame serially, and initialized the pose of the biomechanical model using the pose from the previous contiguous frame. In the first frame, the biomechanical model’s base pose was used to initialize the optimization.

Applying STAC involves two steps: calibrating the biomechanical model to s-DANNCE keypoints, and registering the calibrated biomechanical model to new data. To calibrate the biomechanical model for a given animal and session, we created a training dataset from the initial 10 seconds of data (500 frames) and optimized the offsets and joint angles through three alternating iterations of the q- and m-phases. For each q-phase, we optimized each frame in the 500-frame trajectory, while for each m-phase, we randomly selected 50 frames from the 500 frame training set. To register the calibrated biomechanical model to all of the data from a given session, we reused the offsets from the calibration step and performed a single q-phase optimization over all frames.

#### Contact determination

To estimate the points of contact between two animals, we automatically identified frames in which the mesh models for two interacting animals intersected. For every frame, we posed the biomechanical models representing both animals in MuJoCo according to the poses registered with STAC and logged the positions of the vertices comprising the mesh models. To detect intersections between meshes, we constructed a bounded volume hierarchy from the vertices and faces comprising one animal’s mesh and queried whether any vertices comprising the other animal’s mesh were contained within the volume. When a vertex was found to be inside of the volume, we logged the identity of the penetrating vertex, the identity of the face closest to the penetrating vertex, the identity of the vertex on that face closest to the point that was the shortest distance between the face and the penetrating vertex, and the frame number. For the purposes of analyses, contacts were defined by these pairs of vertices and the frame number.

In general, the procedure described above is asymmetric such that the contacts estimated by constructing a bounded volume hierarchy from one animal of a pair does not equal the contacts estimated by constructing a bounded volume hierarchy from the other animal. We symmetrized it by performing the procedure twice, swapping the identity of the animal used to construct the bounded volume hierarchy, and combining the collection contacts resulting from both iterations. When applied over a recording, this process generates a collection of vertex-vertex pairs encoding the spatial location of contacts on both animals and the times at which they occurred.

#### Contact validation

To validate the performance of our automatic contact estimation method, we compared its predictions to those of human annotators on a set of frames in which two rats were in close proximity. Using a collection of six 30-minute sessions sampled evenly across the genetic models of ASD, we randomly sampled 1100 timepoints in which the centers of mass of both animals were less than 150 mm apart. We then used Label3D to view the animals from all six camera angles simultaneously. We defined seven regions on the rat’s body to consider in our contact validation (Head, Left forelimb, Left hindlimb, Right forelimb, Right hindlimb, Upper body, and Lower body). The boundaries between each of the body parts was defined by grouping bones of the biomechanical model into each of the seven categories, noting the bones with the highest weight for each vertex, and assigning each vertex to one of the seven categories according to the category of its maximum-weight bone. For each frame in the dataset, three human labelers manually labeled contacts between each of the body part pairs from which we estimated the interlabeler variability in manual annotation. To estimate the intralabeler variability, we had two labelers repeat their annotations for a subset of 500 timepoints.

To validate automated contact estimates, we applied the contact estimation method to the timepoints in the manually labeled dataset. For each timepoint, we organized the resulting contacts into the seven body part categories, denoting a body part contact pair if there existed a vertex contact pair with vertices corresponding to each body part. Because contact events are relatively sparse, we used balanced accuracy as a metric to quantify the performance of our contact estimation method, equally weighting the accuracy of touches and non-touches.

We considered three baselines against which to compare the deformable mesh model’s performance: a ‘keypoints’ baseline in which we derived a contact-detection model directly from s-DANNCE keypoints, the interlabeler variability, and the intralabeler variability. For the keypoints baseline, we aimed to estimate how well one could estimate social contacts by using only the pairwise distance between keypoints on social partners. We defined a contact as any time in which two keypoints were within a threshold of 30 mm apart, and assigned each keypoint to one of the seven coarse regions to compare to manually labeled contacts. We estimated the optimal threshold value with a line search. For the interlabeler variability, we compared the contact estimates across the three individuals, computing the average balanced accuracy across all labeler pairs. For the intralabeler variability, we compared the contact estimates from individuals on one day to those from the same individuals on the same data, labeled on a different day, and computed the average balanced accuracy across individuals.

### Touch PCA

To produce a low-dimensional representation of touch within each low-level joint cluster (LLJC), we isolated frames when animals were in each LLJC and found the fraction of time each of the body zones (head, limbs, upper front, and upper rear) were in contact with the other animal for the focal rat and the partner rat (**Fig. 6**). We concatenated these profiles into one 14- dimensional vector and normalized this representation to produce a touch profile for each LLJC where touch occurred for at least 10% of total frames (84 of 156 LLJCs). We performed PCA on the set of normalized touch profiles and found that the first two principal components captured over 75% of the variance (**Supp.** Fig. 12). We used this 2-dimensional space as a complementary way to visualize behavioral changes in ASD knockout models (**Fig. 7**).

### Statistical methods

After reducing each behavioral experiment to a ‘behavioral profile’ or probability vector that captures the usage of each of the fine-grained behavioral clusters (only Action Clusters in the case of lone recordings, both Action and Joint Clusters in the case of social dyads), we compared these profiles between experimental groups. Because there are multiple observations from the same individual when they are paired with different partner subjects within an experimental group, we fit a hierarchical mixed model to perform statistical comparisons^90,91^. This prevented a false assumption of independence between samples from the same individual. Specifically, we fit a linear mixed effects model (LME, Matlab function *fitlme()*) to observations (e.g., the usage of a specific cluster in each recording session) given a fixed effect for experimental group (e.g., ASD KO vs. WT littermate) and a random effect for intercept grouped by subject identity.

We performed statistical comparisons at two levels: one at the level of the behavioral profile for each group and another at the level of usage for each low-level behavioral cluster. In the first case, we compared a behavioral profile from two groups (for example an ASD knockout model and the corresponding WT littermates) by calculating the Jensen-Shannon Divergence (JSD) for profiles measured in each session to the average from one of the groups. These JSD values were the observations used to fit the LME model accounting for subject identity, as described above^90^. At the level of individual clusters between groups, for each cluster we performed the same LME fit, with cluster usages as observations, and corrected the resulting p-values for multiple comparisons using the Benjamini-Hochberg False Discovery Rate (BHFDR) method (MATLAB function *mafdr(…, ’BHFDR’, ’true’)*).

We reported the p-value and JSD between the mean of the two groups for each set of comparisons as well as the number of behavioral clusters from each coarse group (HLAC and HLJC) that was up- or down-regulated in the cohort of interest (**Supp. Table 4, Supp.** Fig. 16). We wrote Matlab wrapper functions *testGroupLme* and *findSigBeh* to identify the group and cluster level significance, respectively, and have included them in the SocialMotionMapper code repository.

### Data availability

All of the kinematic tracks and clustering results presented in this work will be made publicly available for easy access upon publication. For each recording, we will provide the 6-camera video data, as well as the 3D keypoint predictions obtained using s-DANNCE, individual and social classes produced in our analysis (LLACs, HLACs, LLJCs, and HLJCs), and metadata describing the recording (experimental context, experimental group, date of recording, rat ID, amphetamine status, and partner genotype) detailed in **Supplementary Table 1**.

## Author Contributions

UK and DA built the acquisition rig. UK and JFA performed all behavioral recordings. TL built the s-DANNCE model and performed training, inference, and benchmarking of markerless tracking. DA performed fitting to the volumetric model and validation of touch detection. UK performed all behavioral analyses. TWD and BPO supervised and funded this work. UK, TL, DA, BPO and TD wrote the manuscript with input from all authors.

## Acknowledgements

This work was supported by grants to UK (SFARI BTI Fellowship), DA (NIH D-SPAN Award 1F99NS125834-01A1), BPO (SFARI Rat Models Consortium Award #899348, NIH R01GM136972), TD (NIH R01GM136972, McKnight Foundation Technology Award). We also acknowledge helpful feedback on the manuscript from Vikram Chandra.

---

**Supplementary Video 1**: s-DANNCE acquisition, tracking, and volumetric fitting. Top left, ample representative clip of social interaction in rat dyad recordings. Top right, 2D reprojection of s-DANNCE tracked keypoints on single-camera view. Bottom left, 3D keypoint representation for movie snippet. Bottom right, deformable mesh model overlay visualized for corresponding movie frames.

**Supplementary Video 2**: Tracked keypoints for animals from five sets of experimental recordings obtained using s-DANNCE. First, tracked keypoints from six lone wild-type Long Evans rats across five days, day 4 indicates acute dosing with 1.25mg/kg amphetamine twenty minutes prior to recording. Second, WT-WT social pairings across wild-type rats. Third, acute amphetamine dosing experiments from the same wild-type cohort where one animal was dosed with 1.25 mg/kg amphetamine twenty minutes prior to recording. Fourth, tracked keypoints from six littermates from the *SCN2A* cohort, where knockout animals are indicated. Fifth, *SCN2A* knockout and wild-type littermates during social interaction.

**Supplementary Video 3**: Visualization of multi-scale behavioral mapping and clustering introduced in this manuscript. For a rat dyad recording, kinematic tracks from each individual are independently embedded into the individual or animal autonomous map to label each frame with an LLAC (low-level action class). Separately, the kinematics and shared features of the interacting animals are embedded from the perspective of each animal to label each interacting with a LLJC (low-level joint class).

**Supplementary Video 4**: Examples from the catalog of low-level action classes (LLACs) and low-level joint classes (LLJCs). First, eight examples of individual LLACs are shown, along with their corresponding high-level labels (HLACs). 16 3D reconstructions of tracked rat skeletons are translationally and rotationally aligned to best show the movements captured by each class. Next, examples from seven LLJCs are shown, one example from each of the high-level joint classes (HLJCs). For each of the nine instances from each social class shown, the principal or ‘perspective’ rat is shown with colored markers, and the partner with black markers.

**Supplementary Video 5:** A sample clip of social interaction with corresponding behavioral labels for both individual animal action classes (HLACs) and the joint social interaction classes (HLJCs).

**Supplementary Video 6**: Examples of the extension of 3D kinematics tracking by s-DANNCE to rat triplets, mouse dyads, and in dyads placed in an arena with bedding.

**Supplementary Figure 1:**
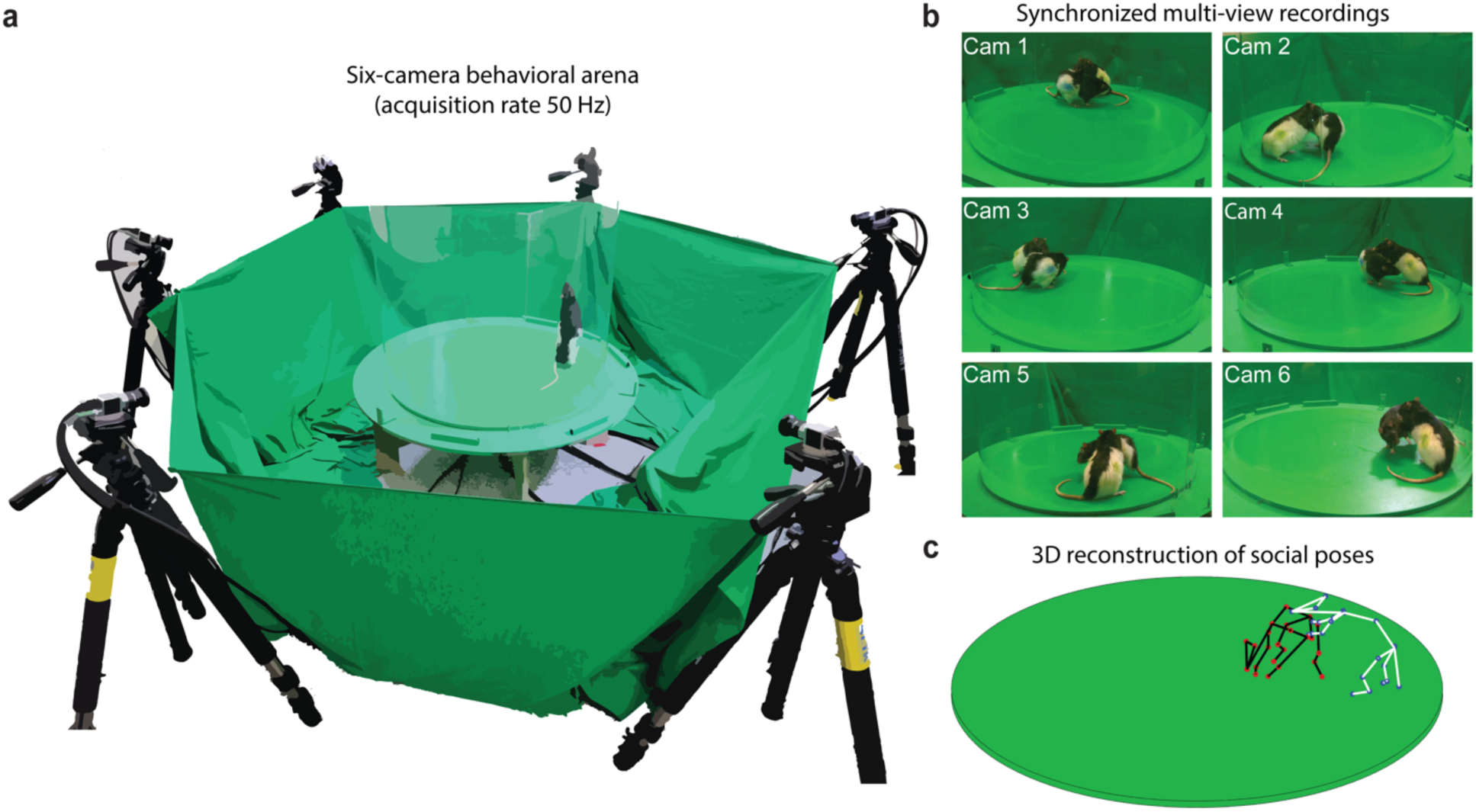
Multi-camera behavioral arena. **a**, Circular behavioral arena surrounded by six synchronized 50 Hz color cameras for recording rat dyadic interactions. **b**, Single timepoint capture of six-camera acquisition from the behavioral arena. **c**, Corresponding 3D social poses inferred by s-DANNCE from frames in **b**.

**Supplementary Figure 2:**
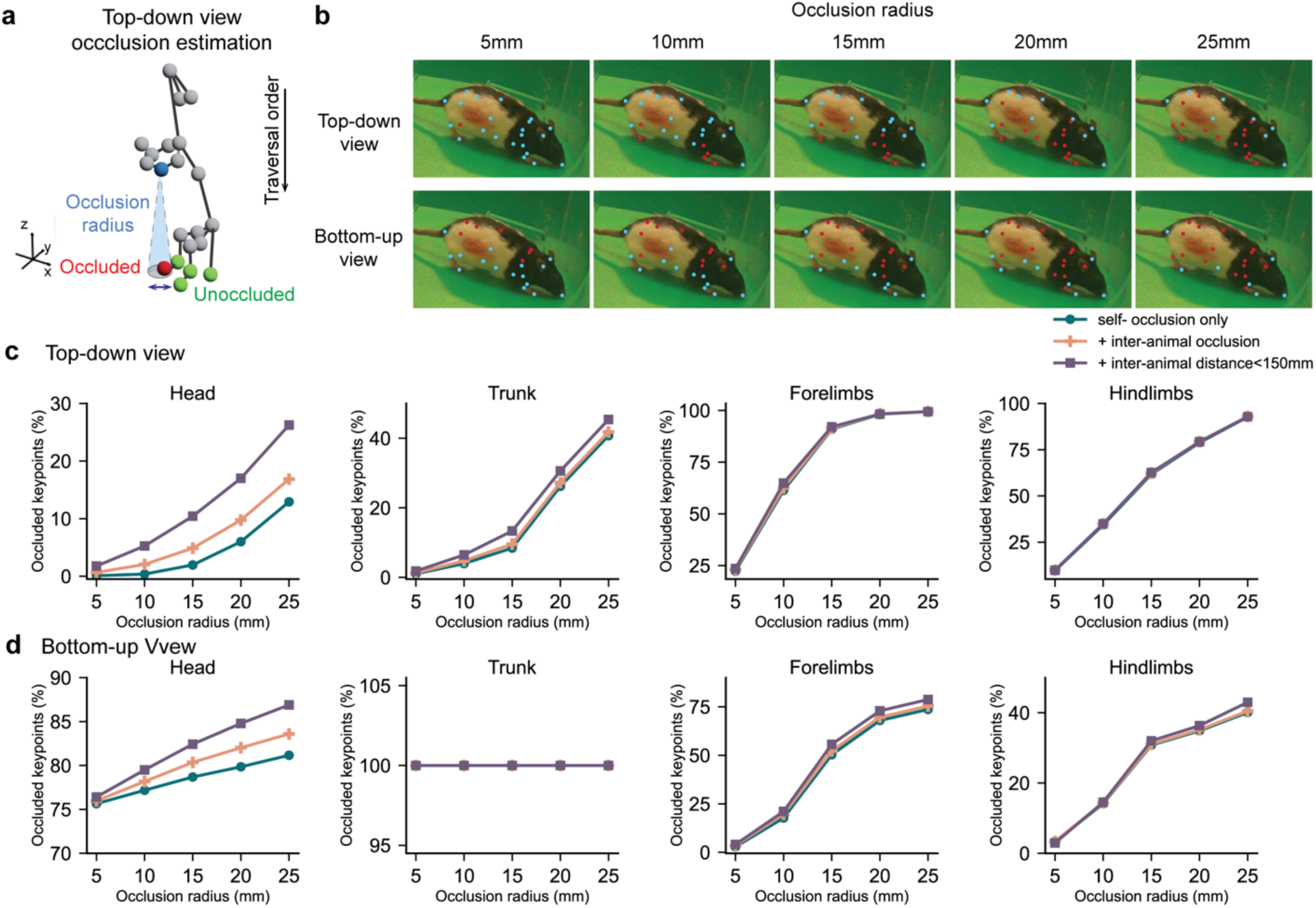
Occlusion analysis in social animals. **a**, Schematic demonstrating how occlusion between keypoints was estimated in an imaginary top-down camera view. Blue color indicates the keypoint of interest and its affected region following the traversal order. Similar procedure was applied for estimating bottom-up occlusion. **b**, Qualitative examples of how occlusion varies with different occlusion radii. Red indicates that the keypoint is assumed to be occluded. **c-d**, Quantitative comparisons of the proportions of occluded keypoints in the animals’ head, trunk, forelimbs and hindlimbs, respectively from the top-down and bottom-up view. The occlusion rates are independently computed for different occlusion radii (5, 10, 15, 20, 25 mm) and except for self-occlusion from the animal’s own body (blue), we optionally consider occlusion from social interactions (orange, purple).

**Supplemental Figure 3.**
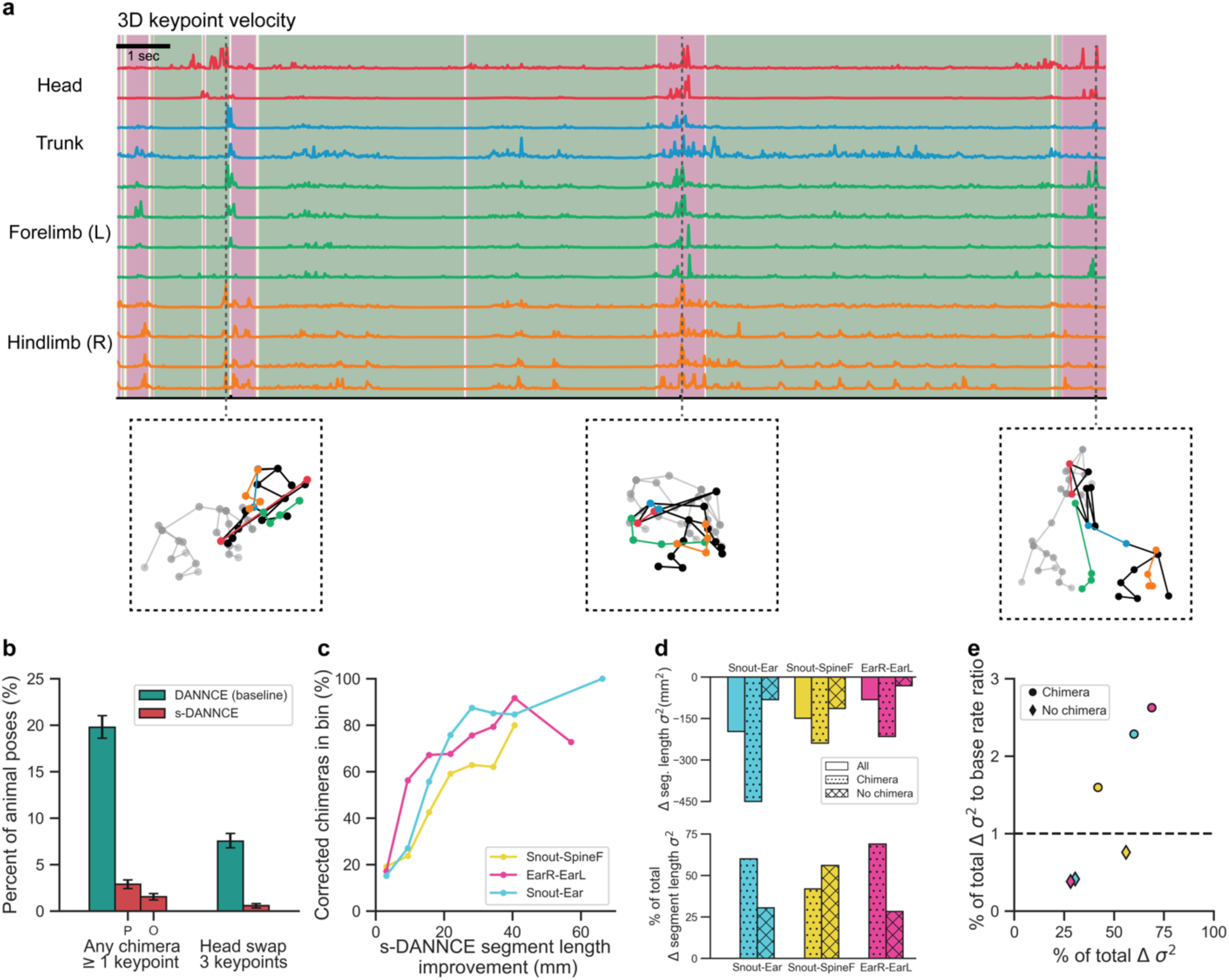
s-DANNCE corrects chimeric errors, driving improved anatomical stability. **a,** Failure cases of baseline DANNCE in interacting animals. Top, first-order velocity traces of 12 selected body markers in one animal engaging in social interactions (t = 20 seconds). Background colors indicate the dynamics in inter-animal distances (pink: < 150 mm). Bottom, qualitative examples sampled from timestamps with sudden velocity spikes. Strong correlations between close inter-animal distances implying social activities and chimeric errors in DANNCE predictions are observed. **b,** Percentage of poses with chimeric errors in close interaction frames (inter-animal distance 150 mm or less), for both baseline DANNCE and s-DANNCE. The overall s-DANNCE chimera rate includes partially fixed chimeras (P; chimeras with fewer keypoints than baseline DANNCE) and all others (O). Error bars are 95% confidence intervals. **c,** Plot showing the rate of s-DANNCE chimeric error correction as a function of s-DANNCE segment length improvement magnitude (the amount of reduced segment length deviation from the mean). Larger s-DANNCE segment length improvements were predominantly from frames where s-DANNCE corrected chimeras, illustrating the impact of chimeric error correction on segment length stability metrics. **d,** Top, change in segment length variance (σ^2^) from baseline DANNCE to s-DANNCE for the indicated body segments, shown for all frames (n=6500) and frames with (n=1706) and without (n=4794) baseline DANNCE chimeras. s-DANNCE variance reduction (improved stability) was larger in frames with baseline DANNCE chimeric errors. Bottom, variance reduction in frames with and without baseline DANNCE chimeras, expressed as percentage of the variance reduction in all frames. **e,** Scatter plot showing the base rate-relative share of segment length variance reduction for frames with and without baseline DANNCE chimeras. Variance reduction in chimera frames has a disproportionately large effect on overall segment length stability.

**Supplementary Figure 4:**
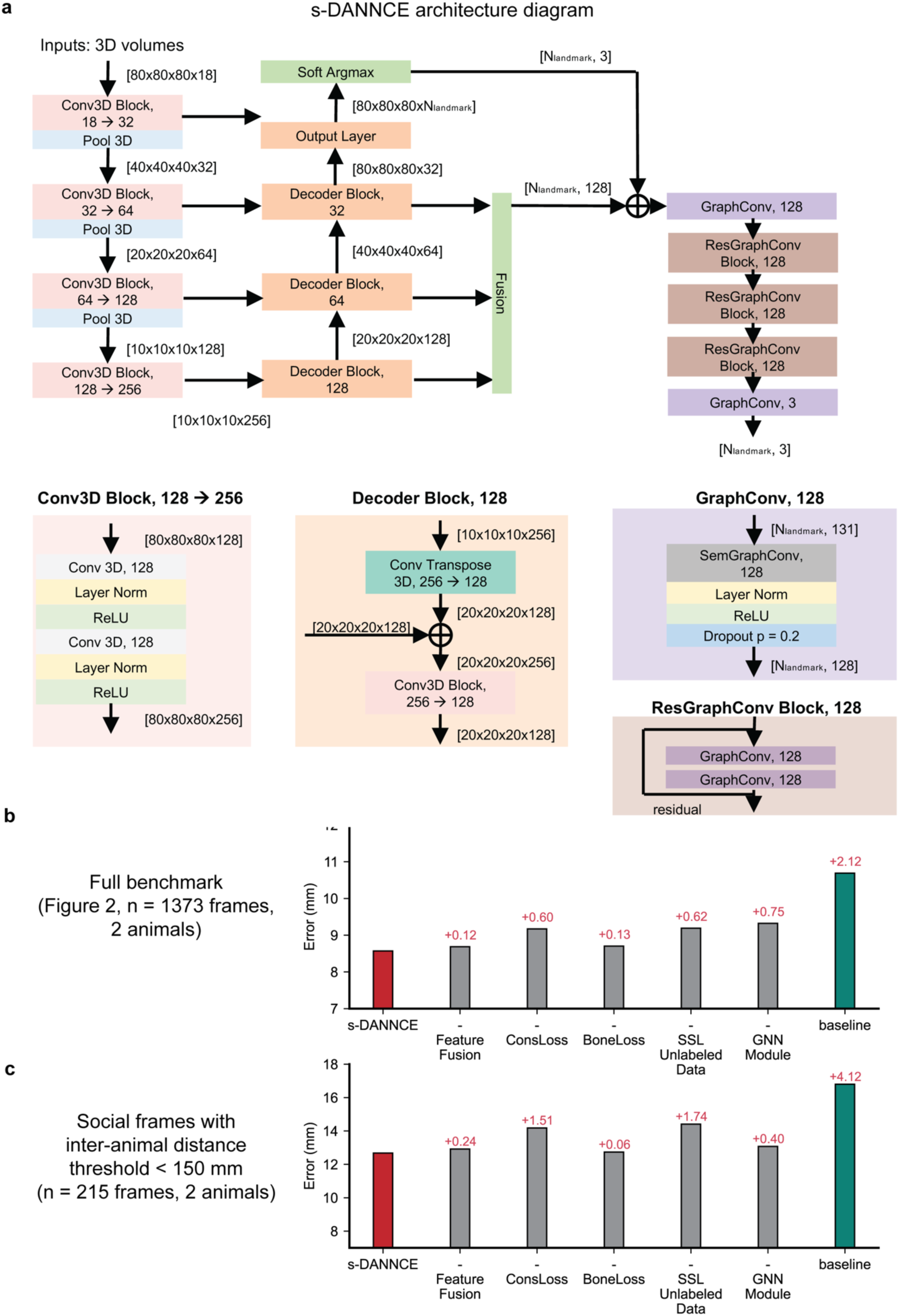
s-DANNCE model architecture. **a**. Detailed schematic of the s- DANNCE neural network architecture, as supplement to **Fig. 2a**. **b-c**, Ablation studies of different modeling components in s-DANNCE. The (-) symbol refers to the ablation of that specific component from the best-performing s-DANNCE model. For the complete descriptions of each modeling component, refer to **Methods**.

**Supplementary Figure 5:**
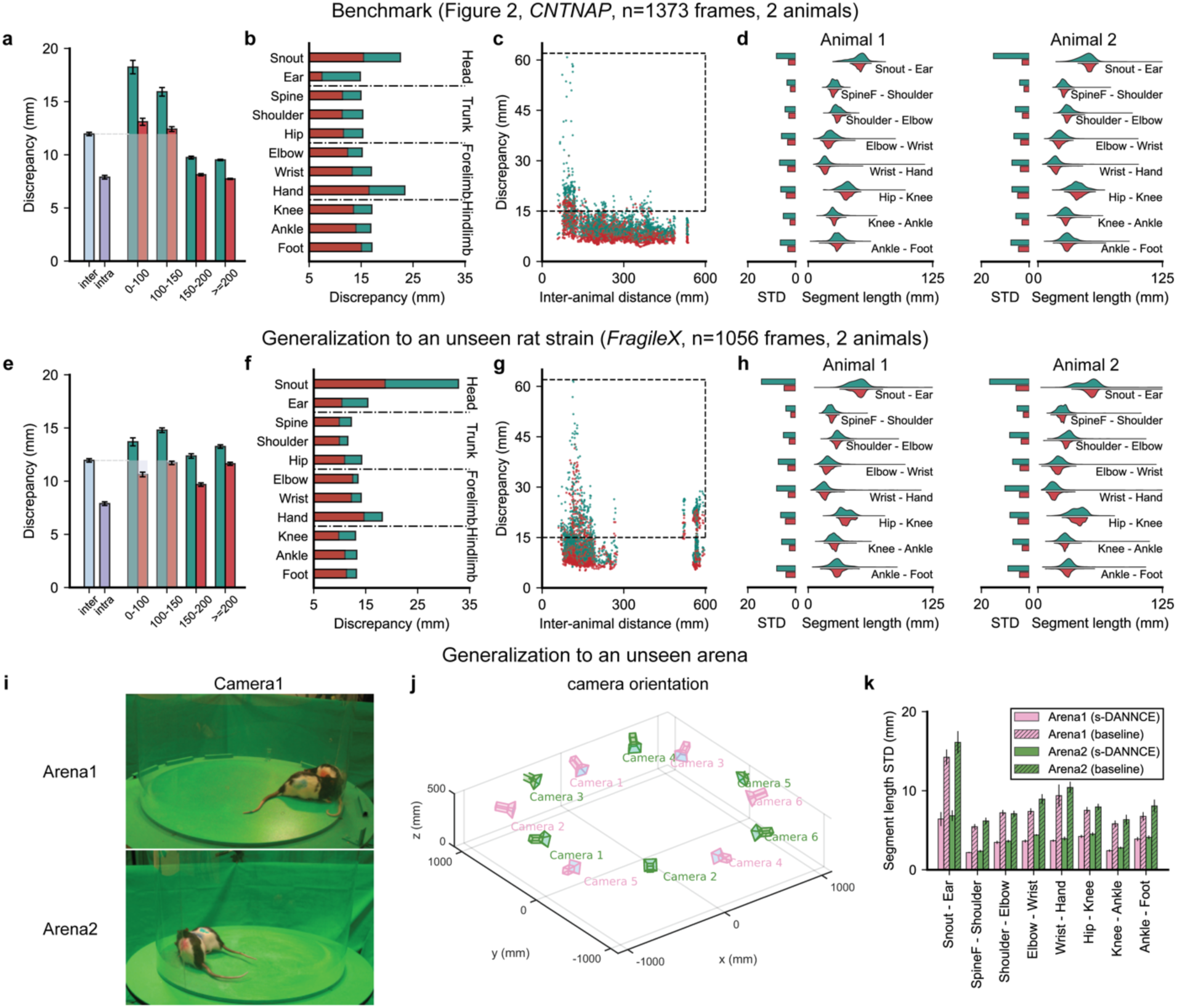
s-DANNCE tracks rats across strains and arena setups. **a-d** are reproduced from **Fig. 2i-j** for comparison: **a**, Bar plots of landmark localization discrepancy over a fully annotated dataset (*CNTNAP,* n = 1373 frames, 2 animals). Frames are grouped by the inter-animal distances (0-100, 100-150, 150-200, >=200 mm). Error bars are 95% CI. The average intra- and inter- discrepancies of the three human labelers (light blue and purple bars, respectively) are computed from **Fig. 2f,g**. **b**, Mean errors with respect to different body landmark positions (n = 215 frames within 150 mm inter-animal distance). **c**, Scatter plot of landmark localization discrepancy as a function of inter-animal distance. **d**, Distributions of body segment lengths as derived from different model predictions in each animal in the recording. Symmetric body segments are merged for conciseness. **e-h,** The same types of analyses as panels **a-d** but performed on a different recording of *FMR1 KO (FragileX)* rats (n = 1056 frames, 2 animals; n = 504 frames within 150mm inter-animal distance). The average intra- and inter- discrepancies of the three human labelers (light blue and purple bars, respectively) in panel **e** are computed from **Fig. 2f,g**. **i-j**, Visualization of two arenas with different camera orientations. Pose labels collected in Arena 1 were used to train the rat models and we examined the shifts in tracking performance when directly generalizing the model to the unseen Arena 2. **k**, Standard deviations in segment length derived from predictions of s- DANNCE and baseline DANNCE respectively in each arena. n = 6 social recordings of male Long Evans SCN2A knockouts were sampled to represent tracking performance in Arena 1 and n = 6 social recordings of male Long Evans with amphetamine injections were sampled for Arena 2. Frames were evenly balanced by the corresponding inter-animal distances with a bin size of 100 mm to accommodate for shifts in behavioral profiles. In contrast to baseline DANNCE, the s-DANNCE model demonstrated better robustness against shifts in camera positioning.

**Supplementary Figure 6:**
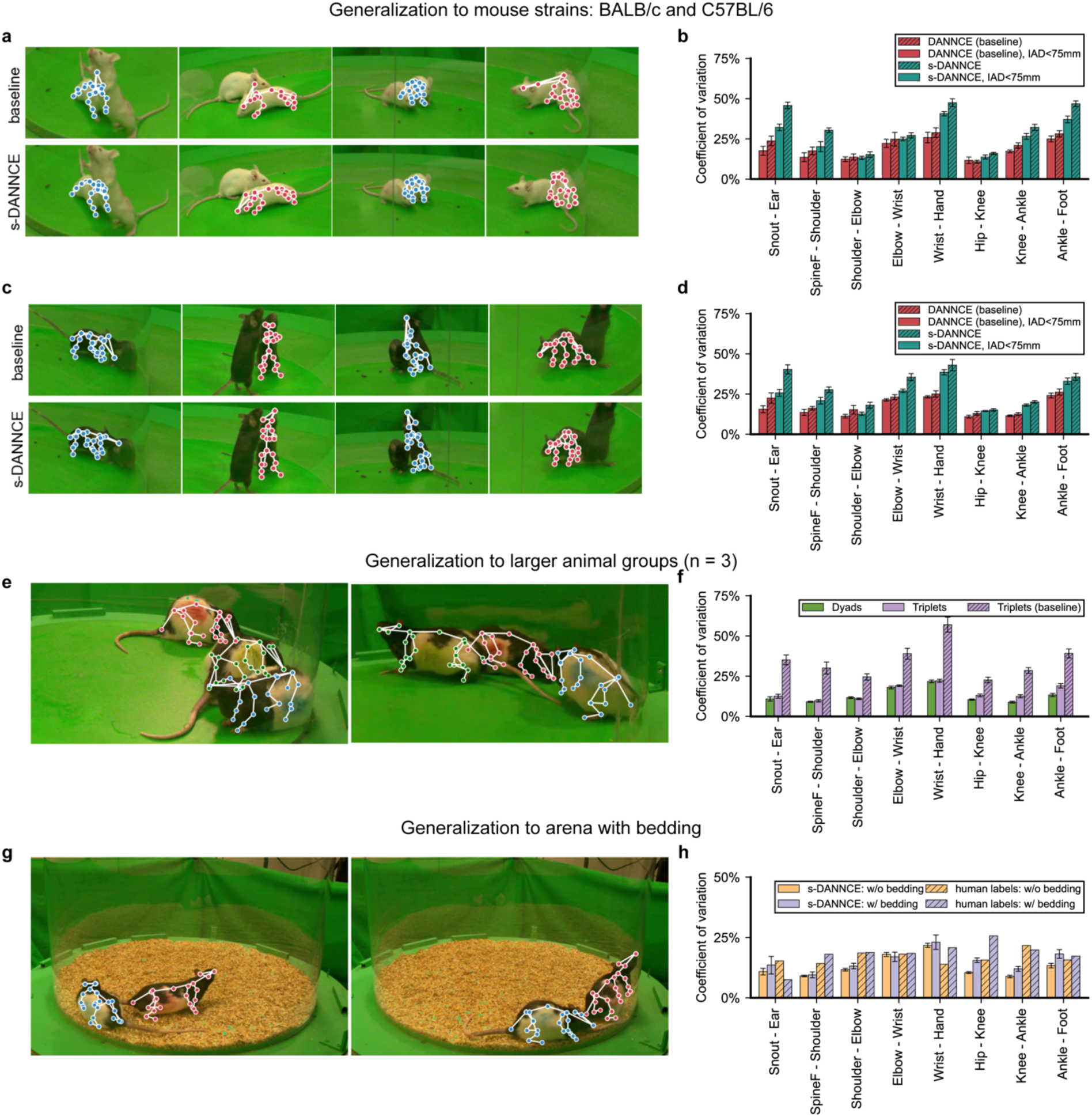
s-DANNCE generalizes across species and experimental conditions. **a-d,** Generalization to mice: Using the pretrained rat models as initializations, we respectively trained baseline DANNCE and s-DANNCE models using n = 111 additional 3D pose labels collected in paired mice. **a**, Qualitative tracking performance comparison between baseline DANNCE and s-DANNCE in BALB/c mice. **b**, Coefficient of variation (%) in segment lengths derived from different model predictions in n = 6 dyad recordings, shown for all frames (n = 180,000) and for frames with inter-animal distance (“IAD”) no greater than 75 mm (n = 39476). **c,d**, Same analyses repeated for C57BL/6 mice. Out of a total of n = 180,000 frames, n = 27052 frames have inter-animal distance no greater than 75 mm. **e-f,** Generalization to larger sized rat groups: **e**, Qualitative tracking examples from a triplet of Long Evans rats when directly applying the s-DANNCE rat model. **f**, Tracking stability in terms of coefficient of variation (%) in segment lengths, as the number of simultaneously tracked animals in the arena scales (n = 2, 3). For the n = 4 triplet recordings (each with n = 90000 frames), we sampled n = 6 dyad and n = 12 lone recordings of SCN2A knockouts, each with a total of n = 12 instances for comparison. **g-h,** Generalization to arena with bedding: All predictions were made by a model finetuned from the pretrained rat model using n = 211 additional 3D pose labels collected in an arena with bedding. **g**, Qualitative tracking examples of female Long Evans rats within an arena with additional bedding. **h**, Coefficient of variation (%) of segment lengths derived from model predictions and human annotations, with and without bedding in the arena. For model performance, we compared n = 6 dyad recordings of Long Evans female rats in the bedding arena with n = 6 dyad recordings of male SCN2A knockouts. For human performance, we compared the n = 211 pose labels used for training the bedding model to the n = 160 3D poses annotated by Labeler 1 as used in **Fig. 2f-h**.

**Supplementary Figure 7:**
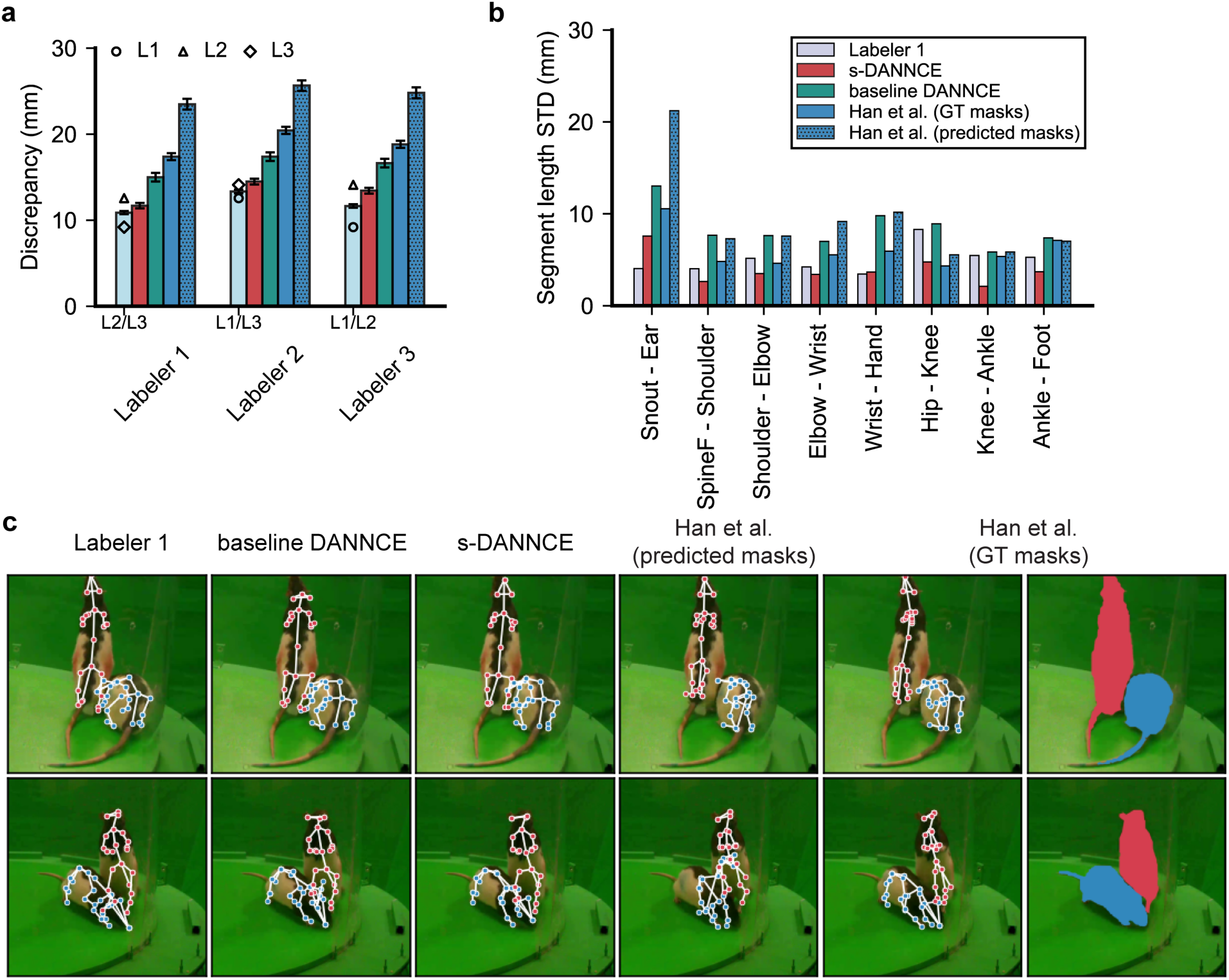
Comparison with 2D instance-masking pose estimation (Han et al. 2024). 3D pose estimation performance of 2D keypoint triangulation when animals are first segmented out of images with 2D instance masks, as proposed in Han et al. 2024. **a,** Bar plots of inter-labeling discrepancies between human and model annotations. Human, baseline DANNCE, and s-DANNCE data are reproduced here from **Fig. 2g**. For the 2D instance masks, we report results from using predicted masks and from using ground-truth (GT) masks. For GT masks, a human labeler annotated n = 3912 identity-preserving instance masks in n = 1980 dyad images, inclusive on the entire dyad dataset used for training s-DANNCE. These GT masks and frames were used for training the 2D pose estimation model (DLC) used for all subsequent evaluations. The labeler also annotated GT masks in all test frames, which were used for the “GT masks” evaluation. For the “predicted masks” benchmark, the training GT masks were used to train a Roboflow 3.0 Instance Segmentation model on the Roboflow platform (starting from the v12 public checkpoint pretrained on MS COCO). This model was then used to predict masks on the test set. For the 2D pose predictions yielded by the Han et al. model, we adopted a triangulation protocol that took the median of 3D reconstruction among all possible camera pairings. **b**, Standard deviations in segment length. Human, baseline DANNCE, and s-DANNCE data are reproduced here from **Fig. 2h**. **c,** Qualitative comparison between human ground truth pose labels (“Labeler 1”), baseline DANNCE, s-DANNCE and instance masking methods. GT instance masks were never used for baseline DANNCE and s- DANNCE. Compared to s-DANNCE, the instance mask method exhibited reduced precision for landmark localization, despite having nearly perfect input masking for “GT masks”.

**Supplementary Figure 8:**
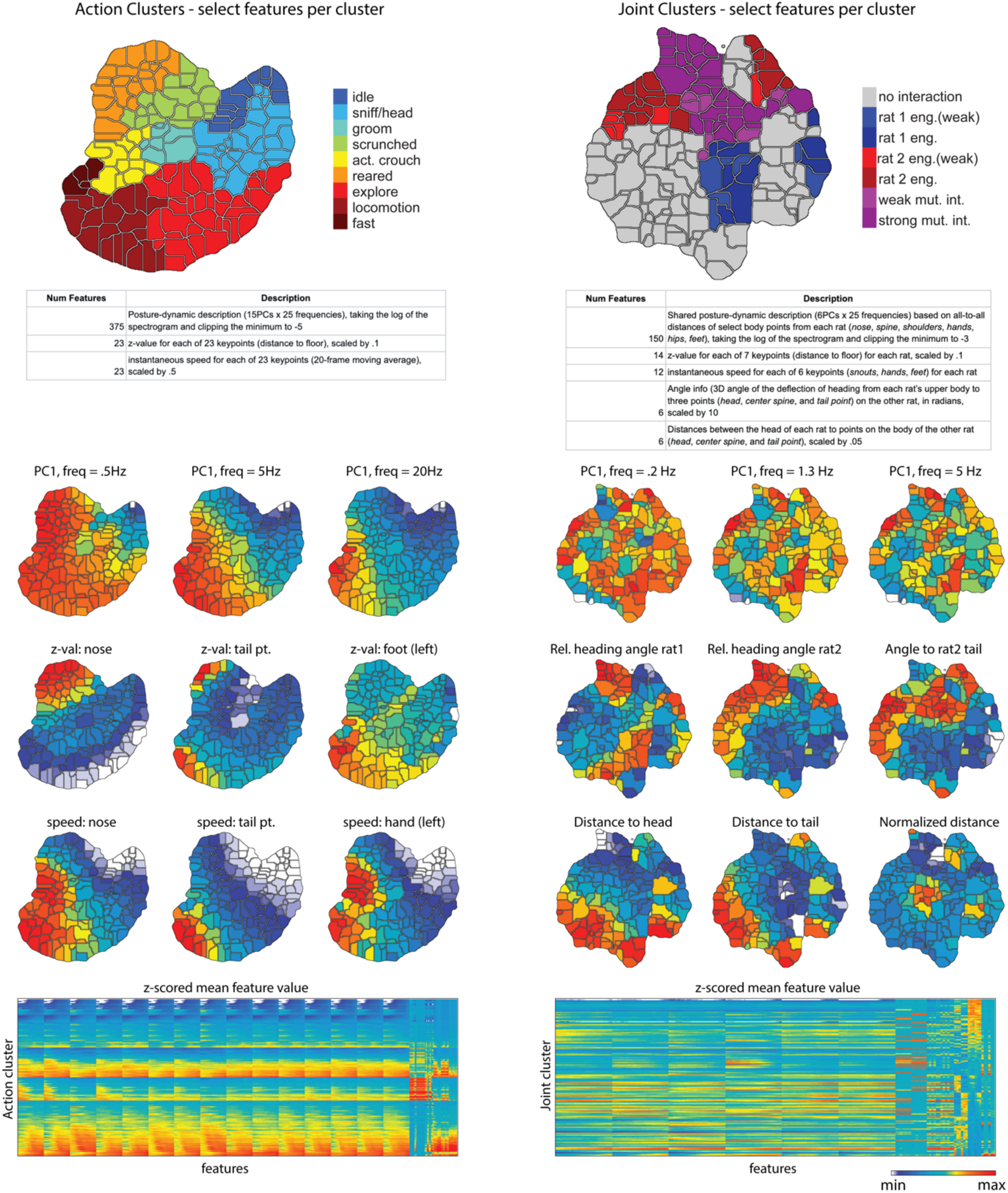
Select features across action and joint clusters. The features used for action clustering (left) and joint clustering (right) are listed below cluster map summaries. The mean value computed across all movies for each action cluster (left) and joint cluster (right) is shown for select features. All mean feature values for action (left) and joint (right) clusters are visualized below and reported in the individual cluster description table (**Supp. Tables 2 and 3**).

**Supplementary Figure 9:**
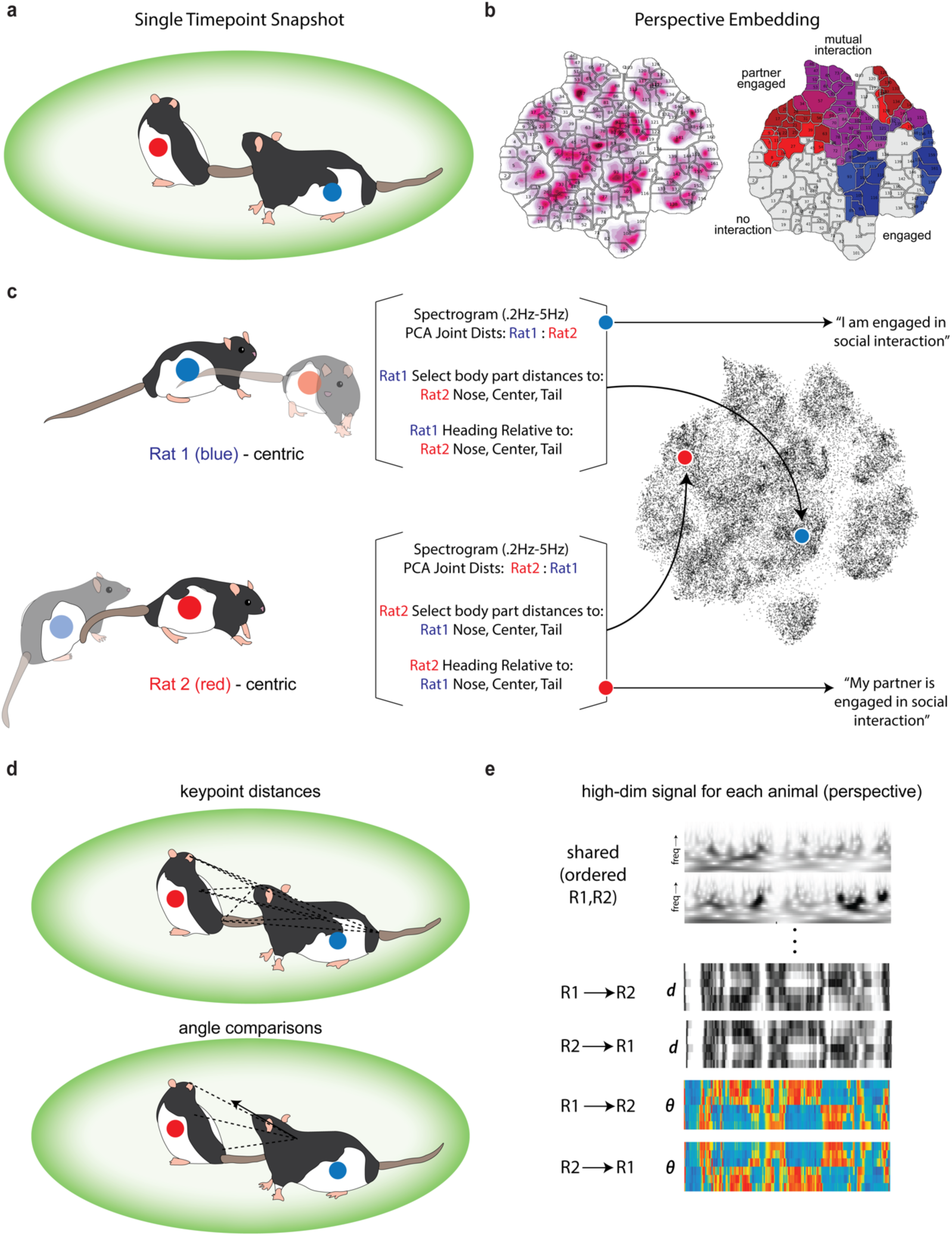
Visualization of dyadic embedding. a, Each point in the social recording contains two animals with identities maintained. b, The shared postural and dynamic information are embedded into a joint map to parse joint behavior motifs. c, The embedding from the perspective of each animal is processed individually and the order of shared kinematic features maintains the ‘perspective’ for each individual. This results in a corresponding point in behavioral space to each animal at each point in time. Behavioral labels then describe the shared interaction from the perspective of that animal. d, Examples of features from the perspective of one animal (blue). e, Construction of the shared high-dimensional feature vector from the perspective of one animal includes information about both animals but maintains ordering with the focal animal first.

**Supplementary Figure 10:**
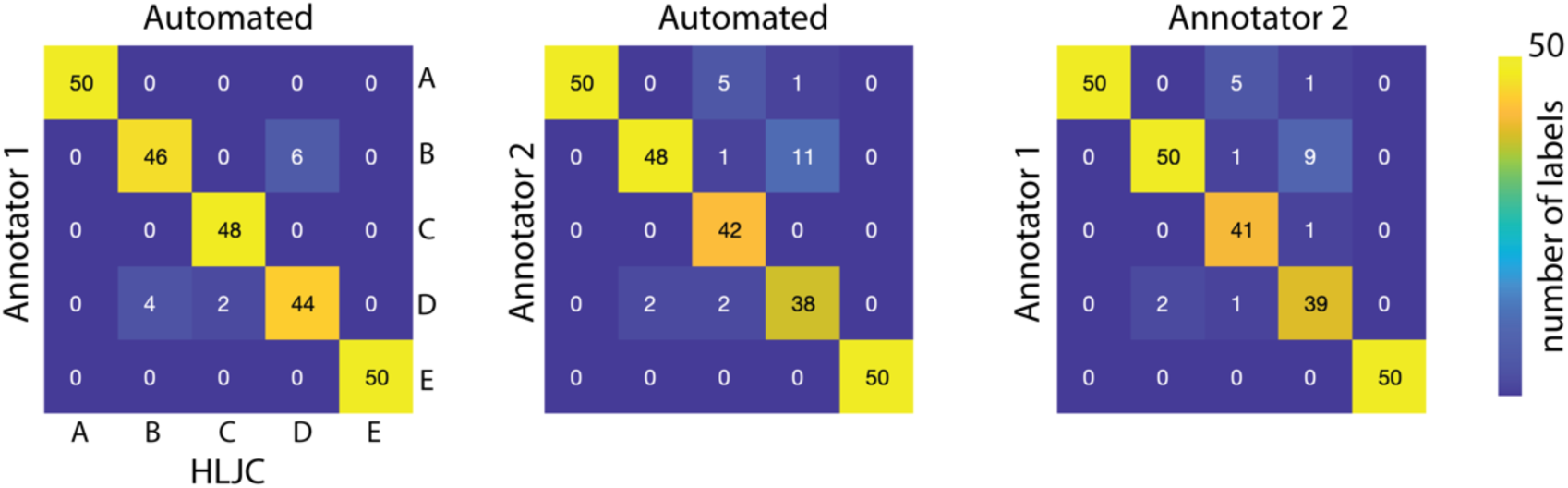
Assessing accuracy of dyadic embedding derived clusters. We compared manual annotation of 250 snippets of social data consisting of 50 snippets each from five low-level joint clusters derived from our unsupervised social clustering. Two annotators were given instructions to assign one of five behavioral groups to each cluster: *no interaction* (far apart, little movement), *rat1 engaged* (rat1 following rat2, rat2 avoiding)*, rat2 engaged* (rat 2 investigating rat1 from behind), *mutual interaction* (close head-to-head, allogrooming), and *no interaction* (mutual rearing). The annotators had 95.2% and 91.2% agreement with the original cluster labels, and there was 92.0% agreement between the labelers.

**Supplementary Figure 11:**
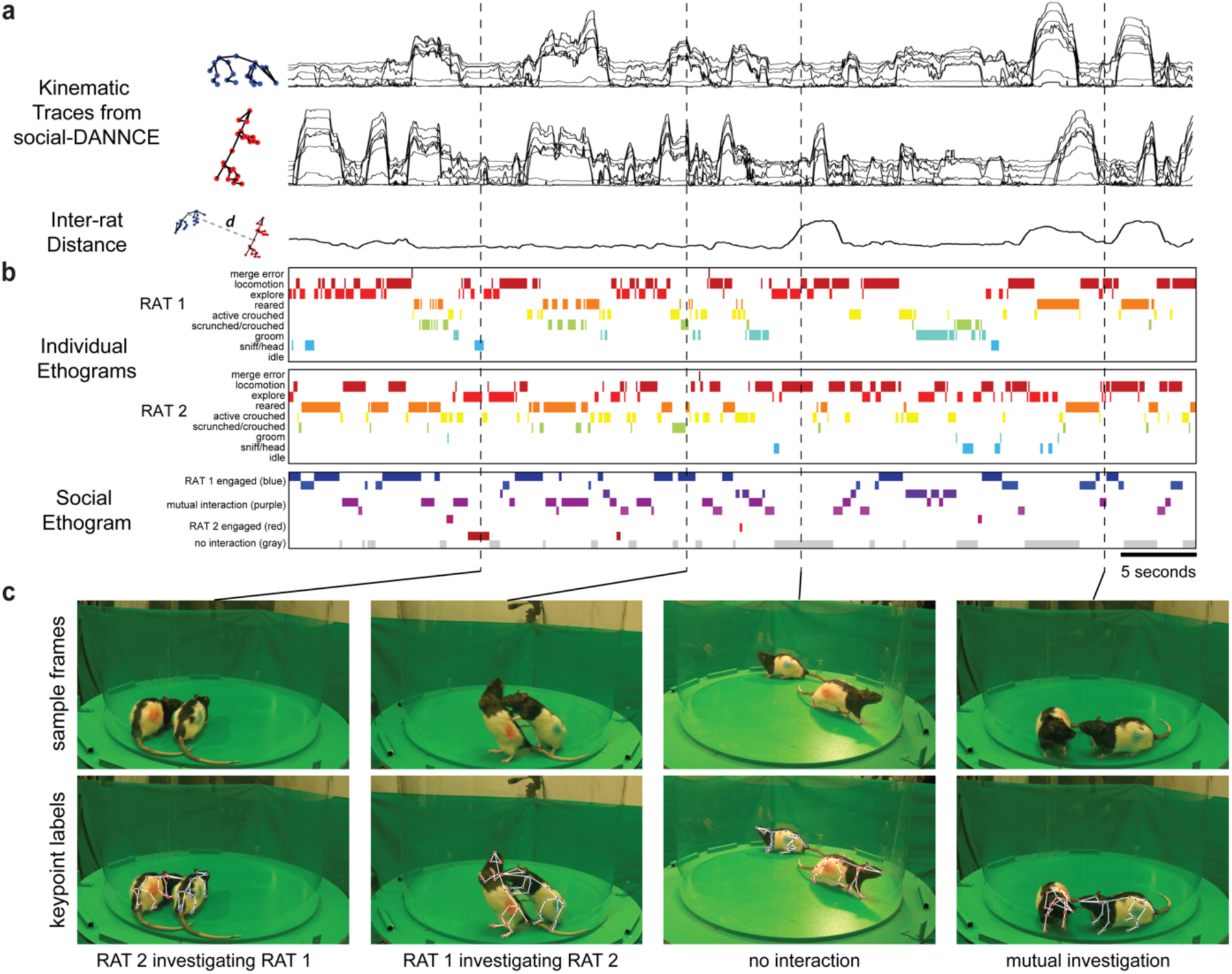
Behavioral analysis of rat dyad recordings. **a**, Sample kinematic traces and distance between animals is plotted for a 60-second bout of social interaction. **b**, Classical embedding labels for each animal and the joint social embedding are plotted as time-series ethograms with corresponding behavioral labels as defined in Figure 2. **c**, Snapshots from recorded sample interaction are shown for four different social behaviors overlaid with skeletons inferred from s-DANNCE.

**Supplementary Figure 12:**
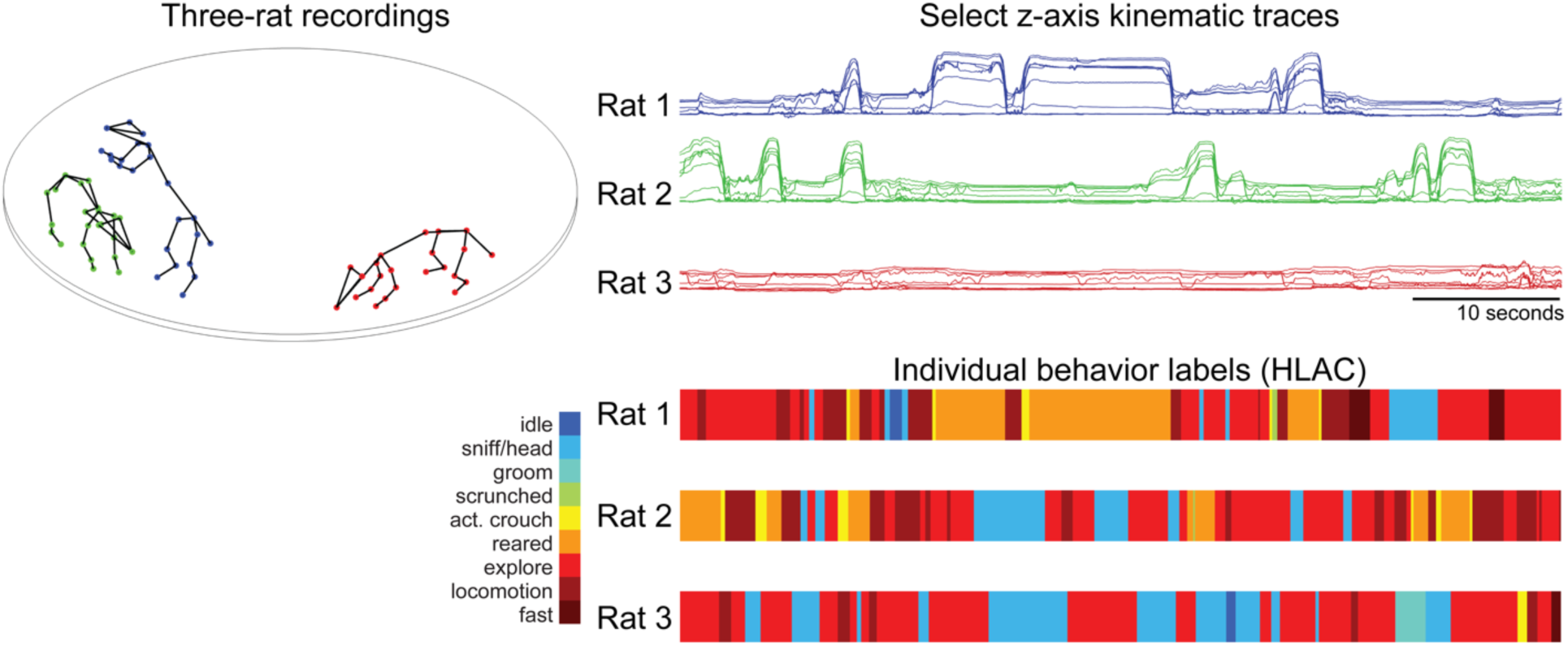
Behavioral analysis for rat triad recordings. s-DANNCE-inferred keypoint trajectories from rat triads were processed individually to produce individual level behavioral labels for each rat. A snapshot of postures (left), z-traces of select keypoints over one minute of recordings (top right), and ethograms representing high-level action clusters (HLACs) for the corresponding behaviors (bottom right) are shown.

**Supplementary Figure 13:**
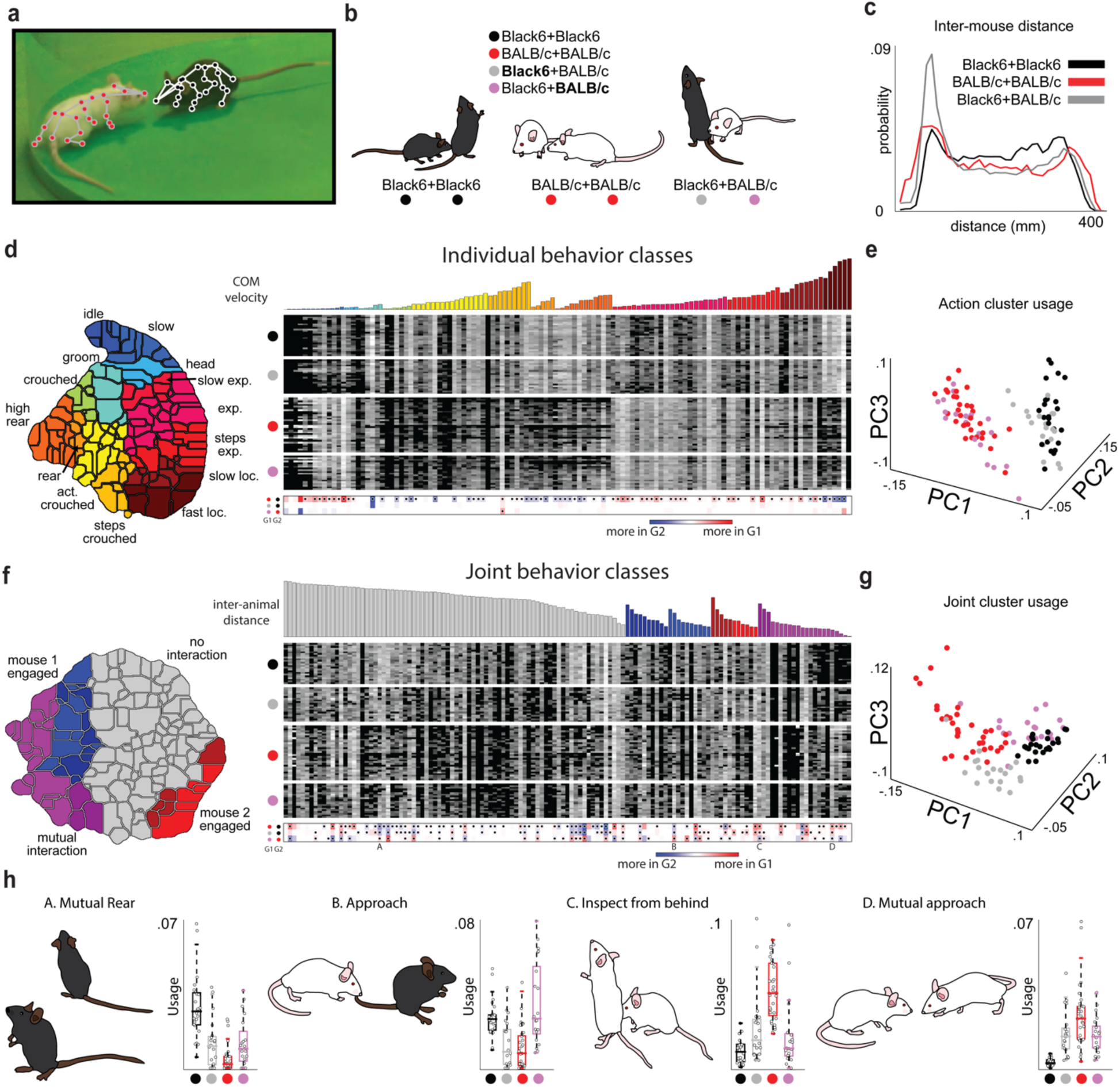
Social phenotyping in C57BL/6 (Black6) and BALB/c mouse strains. **a**, An s-DANNCE model was used to track 3D keypoints in mouse pairs during social interaction. **b**, Black6+Black6, BALB/c+BALB/c, and mixed Black6+BALB/c pairs were recorded in a behavioral arena. **c**, Histograms of inter-animal distances across the three types of pairings. **d**, Postural and kinematic features were sampled from all recordings and used to generate a behavioral map, which was then split into clusters using a watershed transformation. Manual annotation of the map was performed by viewing movies corresponding to a given cluster randomly sampled from all recordings. Action class clustering applied to each mouse individually was used to generate behavioral profiles for each animal within each pairing. Below, differences between mean expressions of three pairs of groups is shown. G1 and G2 refer to the groups specified to the left of each row of the colored comparison heatmap at the bottom of the panel. For example, in the first row, G1 is BALB/c+BALB/c pairings (symbolized by a red dot as in **b**) and G2 is C57BL/6+C57BL/6 (symbolized by a black dot as in **b**). **e**, PCA performed on individual action profiles during social interaction. **f**, Joint class clustering applied from the perspective of each mouse was used to generate joint behavioral profiles, with differences between groups presented as in **d**. **g**, PCA performed on joint profiles during social interaction. **h**, Individual occupancy differences across the four groups are shown for four example joint behaviors.

**Supplementary Figure 14.**
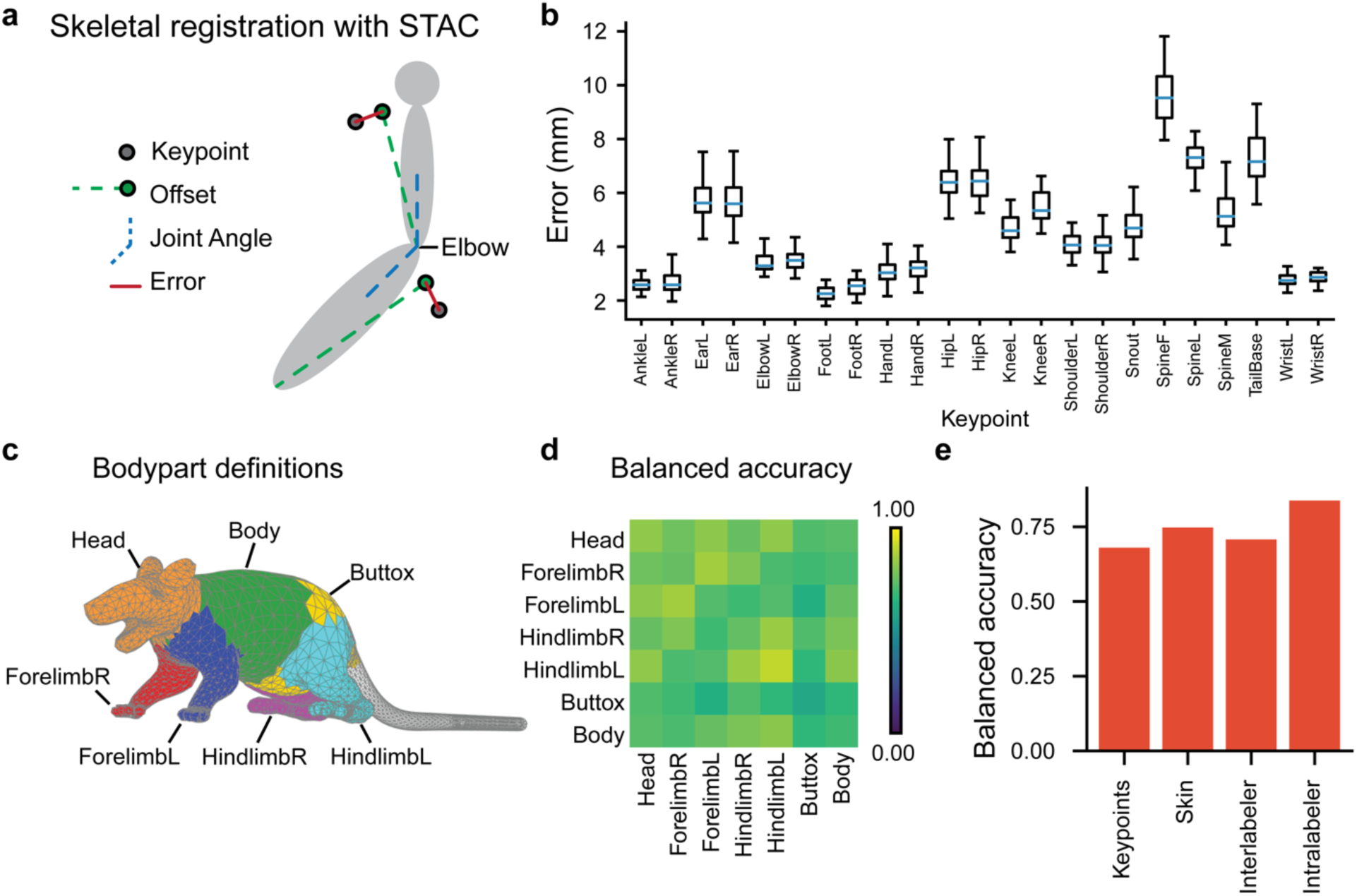
Estimating social touch with skeletal registration and deformable mesh models. A) Schematic of skeletal registration. STAC iteratively optimizes a set of learned offsets that relate points on a skeletal model to keypoints and the joint angles of the skeletal model that best explain the keypoint positions. B) The distribution of average registration errors for all keypoints across sessions. Blue lines indicate the median, box limits indicate the 25th and 75th percentiles, whiskers indicate the maximal point up to 1.5 times the interquartile range from the nearest box limits. C) Schematic depicting the definitions of body parts used for manual contact labeling. D) Balanced accuracy of contacts estimated with the deformable mesh relative to manual contact labeling for all body part pairs. E) Average balanced accuracy of contact quantification methods and human labelers. Balanced accuracy is the average of the true positive rate (the fraction of contacts that were correctly identified) and the true negative rate (the fraction of non-contacts that were correctly identified). The keypoints method defined contacts as any time in which two keypoints on social partners were within 30 mm of one another (see methods). Interlabeler balanced accuracy was the average accuracy between all pairs of distinct human labelers. Intralabeler accuracy was the average balanced accuracy of human labelers labeling the same data on two different days. The skin model achieves a balanced accuracy greater than both the keypoints approach and the accuracy between labelers.

**Supplementary Figure 15:**
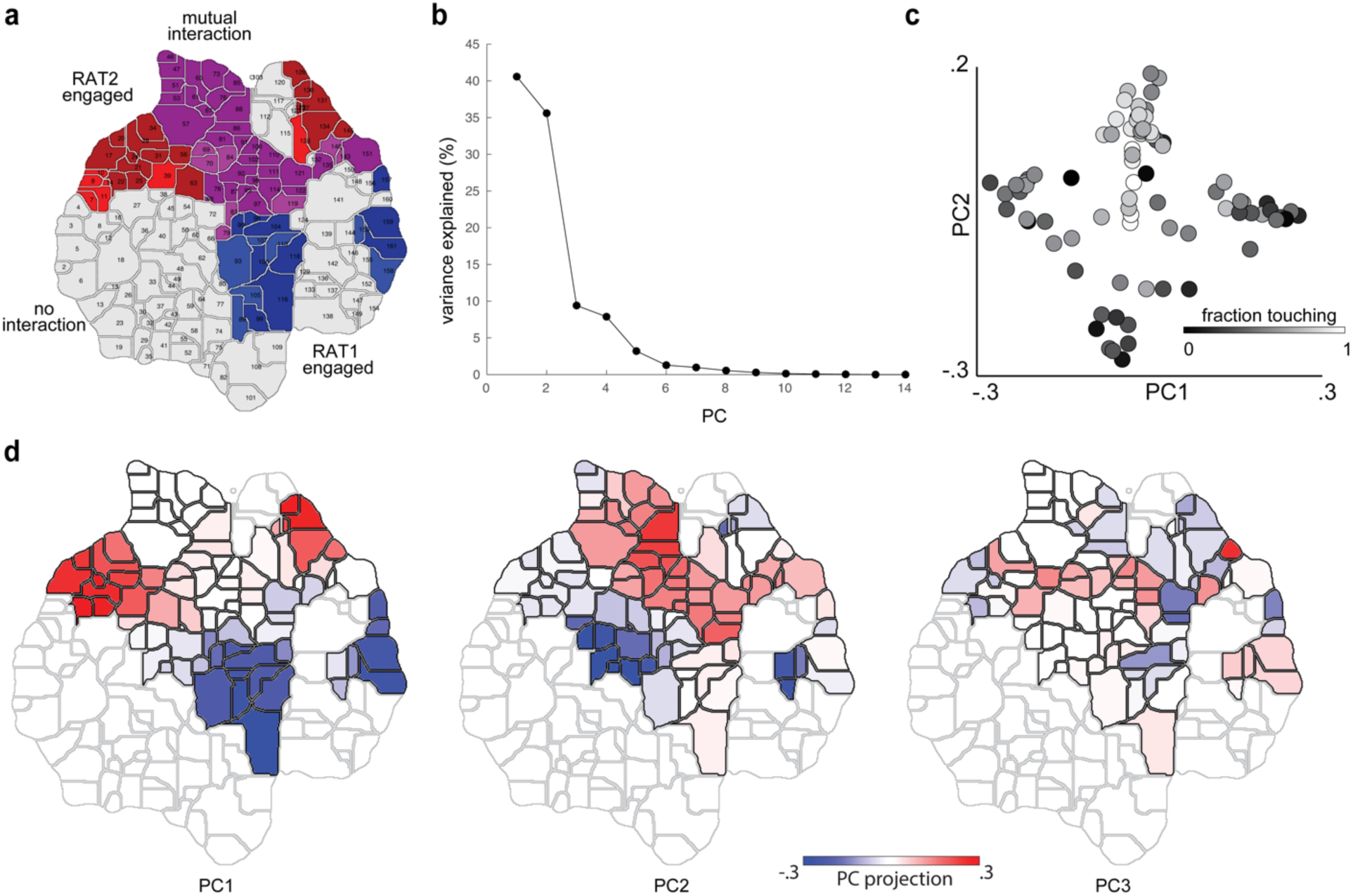
Touch profiles by social cluster. **a**, Joint clustering with high-level category labels. **b**, Variance explained per PC on 14-dimensional (coarse/reduced body map) normalized touch densities for animal1/animal2. **c**, Fraction of time touching for likely-touch (touching >= 10% of the time) clusters on PC axes (refer to **Fig. 5**). **d**, Loadings onto first 3 PC axes for social clusters (non-touch clusters outlined in gray).

**Supplementary Figure 16:**
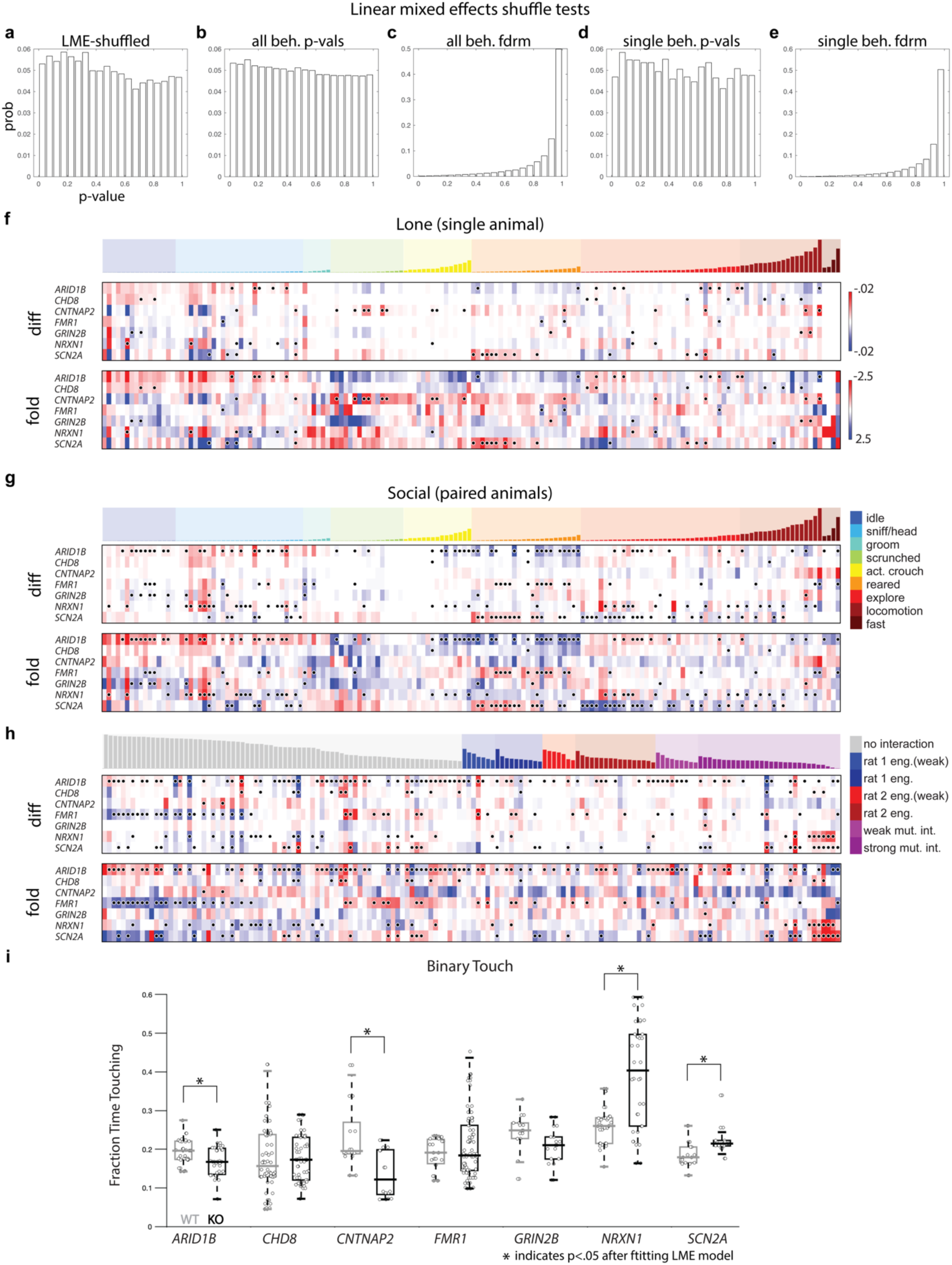
Summary of comparisons for ASD model rat cohorts. **a**, LME model fits were performed on condition- and identity- shuffled data from the WT and amphetamine social experiments. 10000 trials were run with different permutations of group assignment to action cluster usage profiles and the histogram of resulting p-values is reported. **b**, Individual behavior usage was calculated for data shuffled as in **a** and the pooled p-values across all behaviors are reported. **c**, p-values for **b** following a Benjamini-Hochberg False Discovery Rate (BHFDR) correction. **d**, p-value histogram for a single behavior from the shuffled comparison described in **b** is plotted. **e**, p-values for **e** following a BHFDR correction. **f**, Difference and fold change for action cluster usage for each ASD knockout model as compared to wild-type littermates for lone behavior. **g**, Difference and fold change for action cluster usage for each pair of knockout animals as compared to wild-type littermates during social interaction. **h**, Difference and fold change for joint cluster usage for each ASD knockout model as compared to wild-type littermates during social interaction. **i**, Fraction of time with any contact between fitted deformable mesh bodies during interaction across ASD model strains. Significance is indicated after fitting a LME model for each comparison, p < 0.05.

**Supplementary Figure 17:**
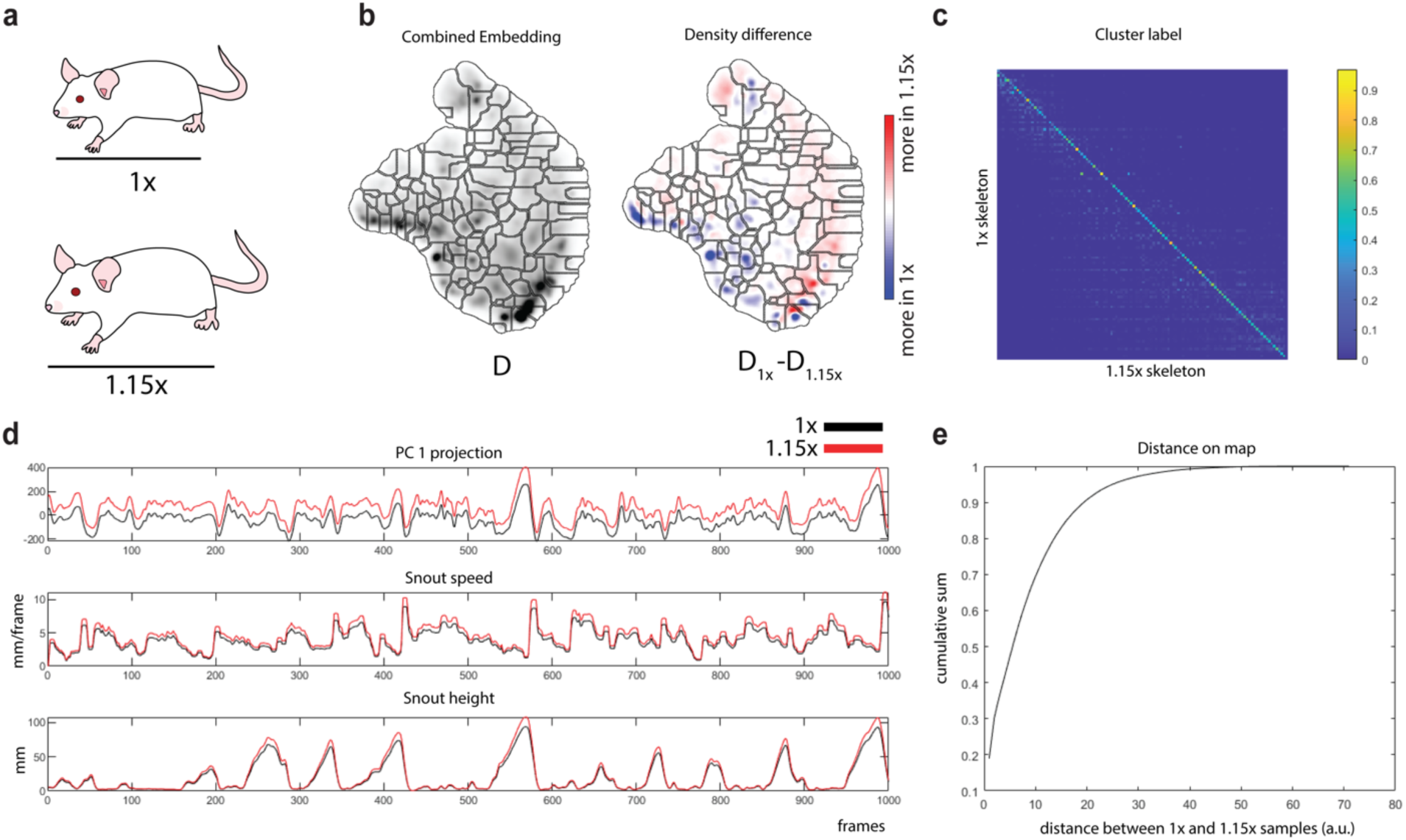
Body-scaled embedding comparison. **a,** A dataset was constructed from all lone mouse recordings, plus a copy with the distance between keypoints scaled up by 15% to test the effect of un-normalized body scale differences on behavioral mapping. **b,** A single behavior map was produced using this dataset, and the total density (left) and difference between the unscaled and scaled densities is shown. **c,** Matrix quantifying the fraction of samples in the 1.15x dataset that are assigned to the same clusters as original 1x samples. Color bar represents the fraction of 1x cluster samples. **d,** Example traces of three different features, showing how they are affected by body scale. **e,** Cumulative distribution function of paired distances between each 1x sample and its 1.15x scaled version.

**Supplementary Figure 18:**
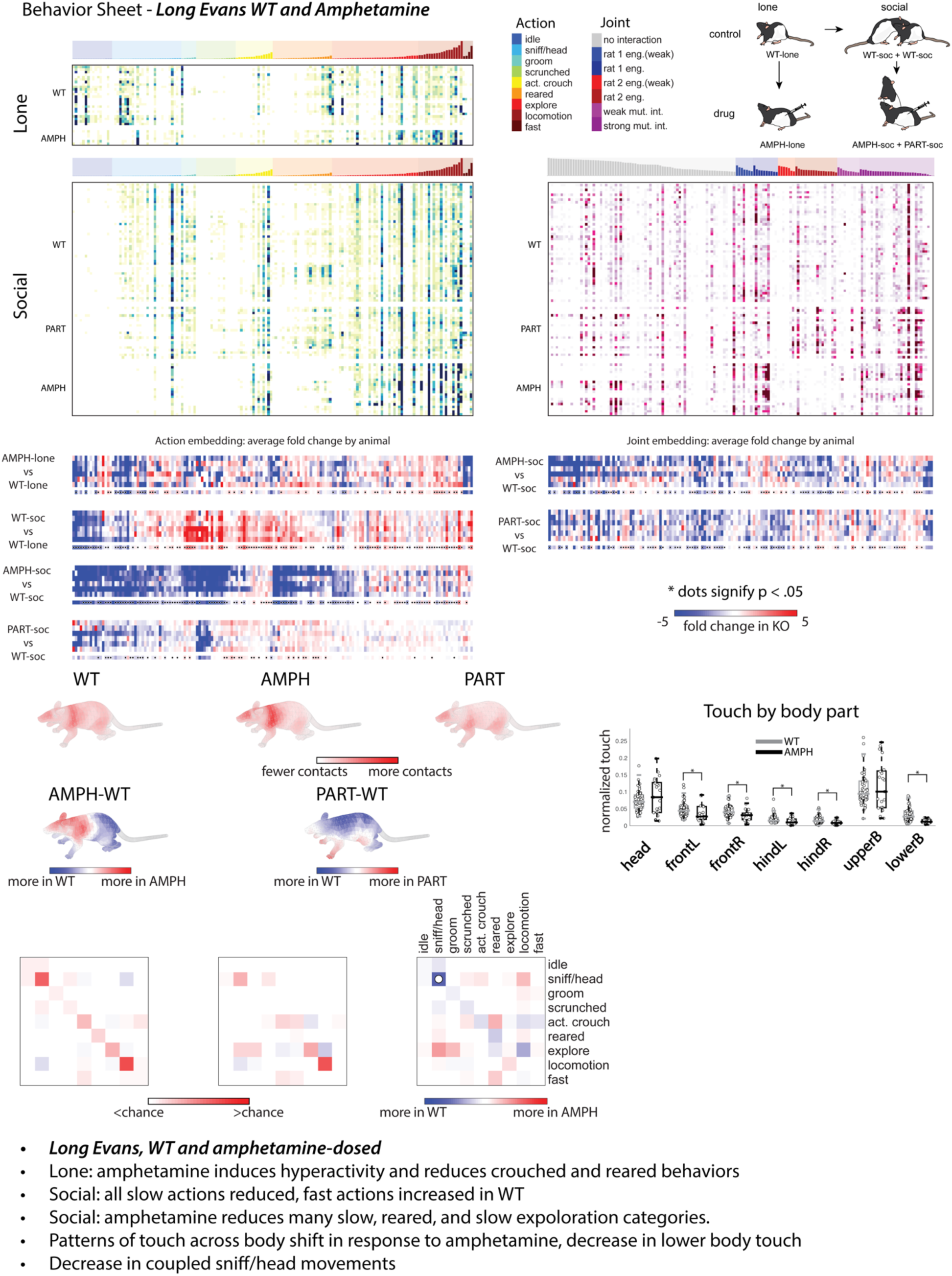
Long Evans wild-type and amphetamine detailed behavior sheet. All raw usage for action and joint clusters is reported. Average differences from mean wild type behavior for each animal are shown for each context. Raw touch for wild-type (WT), amphetamine (AMPH), and partner (PART) are shown, along with differences from average wild-type usage along the body for AMPH and PART trials. The difference in total touch for amphetamine-dosed and wild-type recordings for each body part is shown and significance is indicated with an asterisk. Mutual information calculated for WT/WT and AMPH/PARTNER pairings and differences between the two are shown. Summary of findings is reported.

**Supplementary Figure 19:**
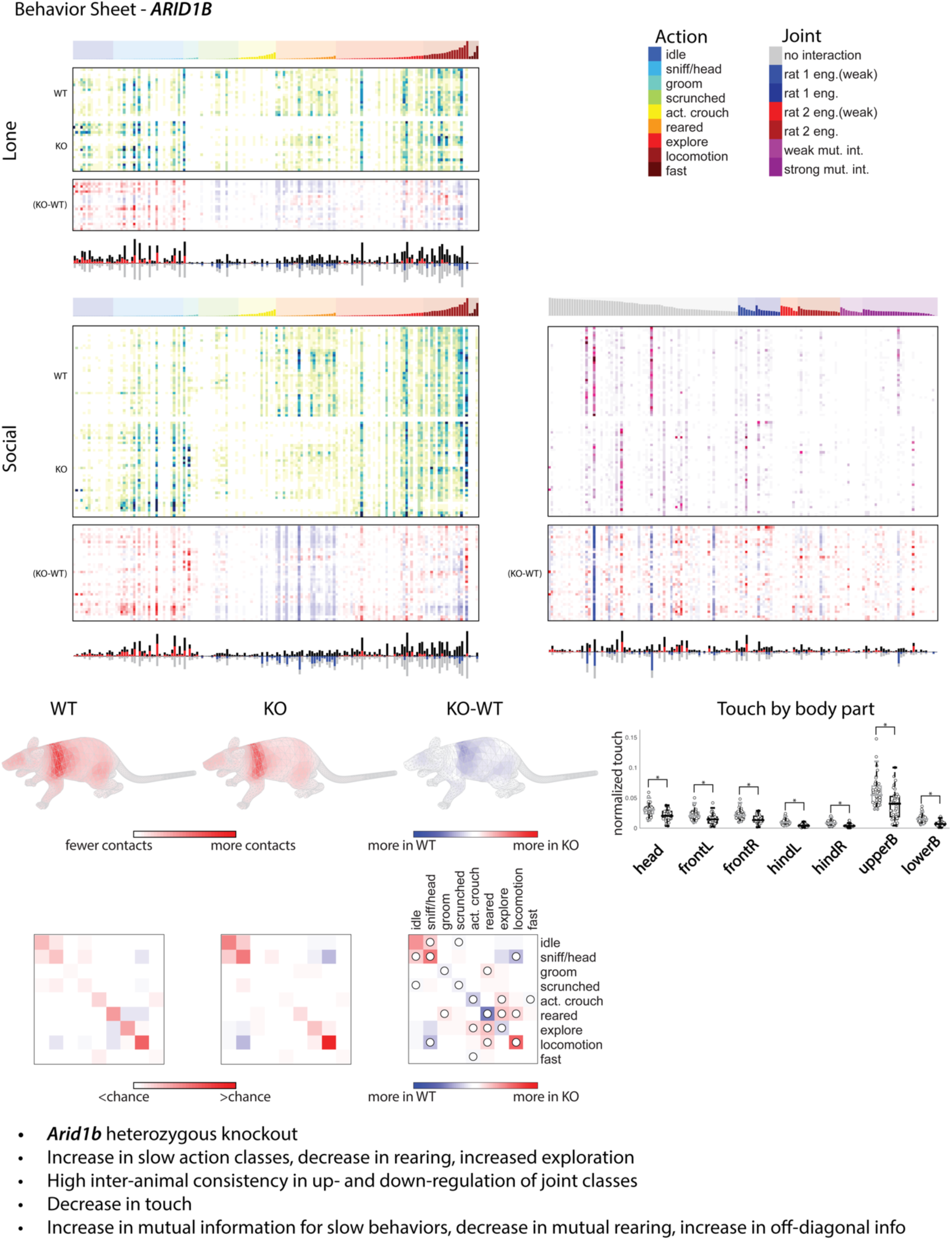
*ARID1B* and wild-type littermate detailed behavior sheet. All raw usage for action and joint clusters is reported. Average differences from mean wild type behavior for each recording are shown. Raw touch and difference for wild-type (WT) and knockout (KO) are shown. The difference in total touch for WT and KO recordings for each body part is shown and significance is indicated with an asterisk. Mutual information calculated for WT/WT and KO/KO pairings and differences between the two are shown. Summary of findings is reported.

**Supplementary Figure 20:**
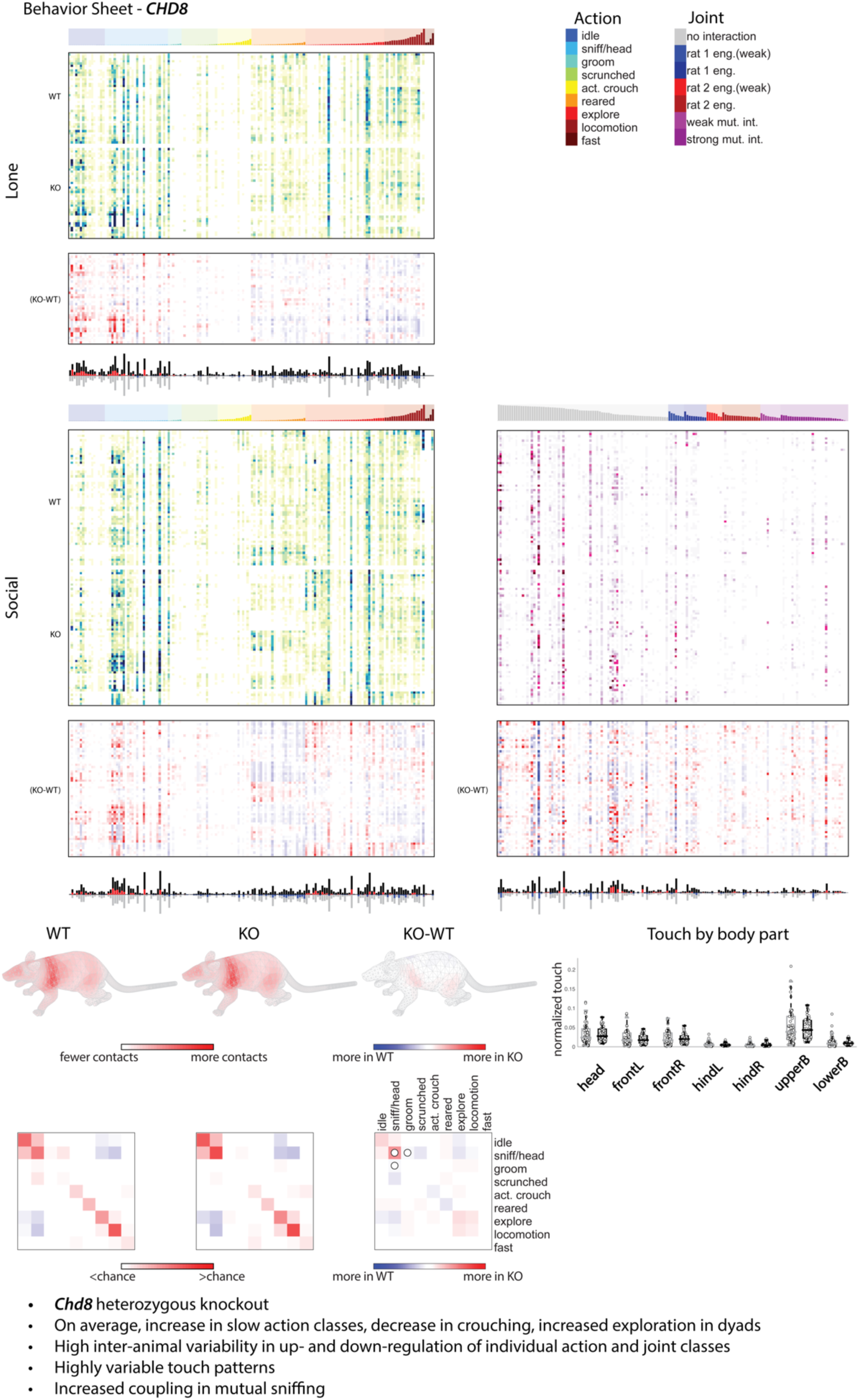
*CHD8* and wild-type littermate detailed behavior sheet. All raw usage for action and joint clusters is reported. Average differences from mean wild type behavior for each recording are shown. Raw touch and difference for wild-type (WT) and knockout (KO) are shown. The difference in total touch for WT and KO recordings for each body part is shown and significance is indicated with an asterisk. Mutual information calculated for WT/WT and KO/KO pairings and differences between the two are shown. Summary of findings is reported.

**Supplementary Figure 21:**
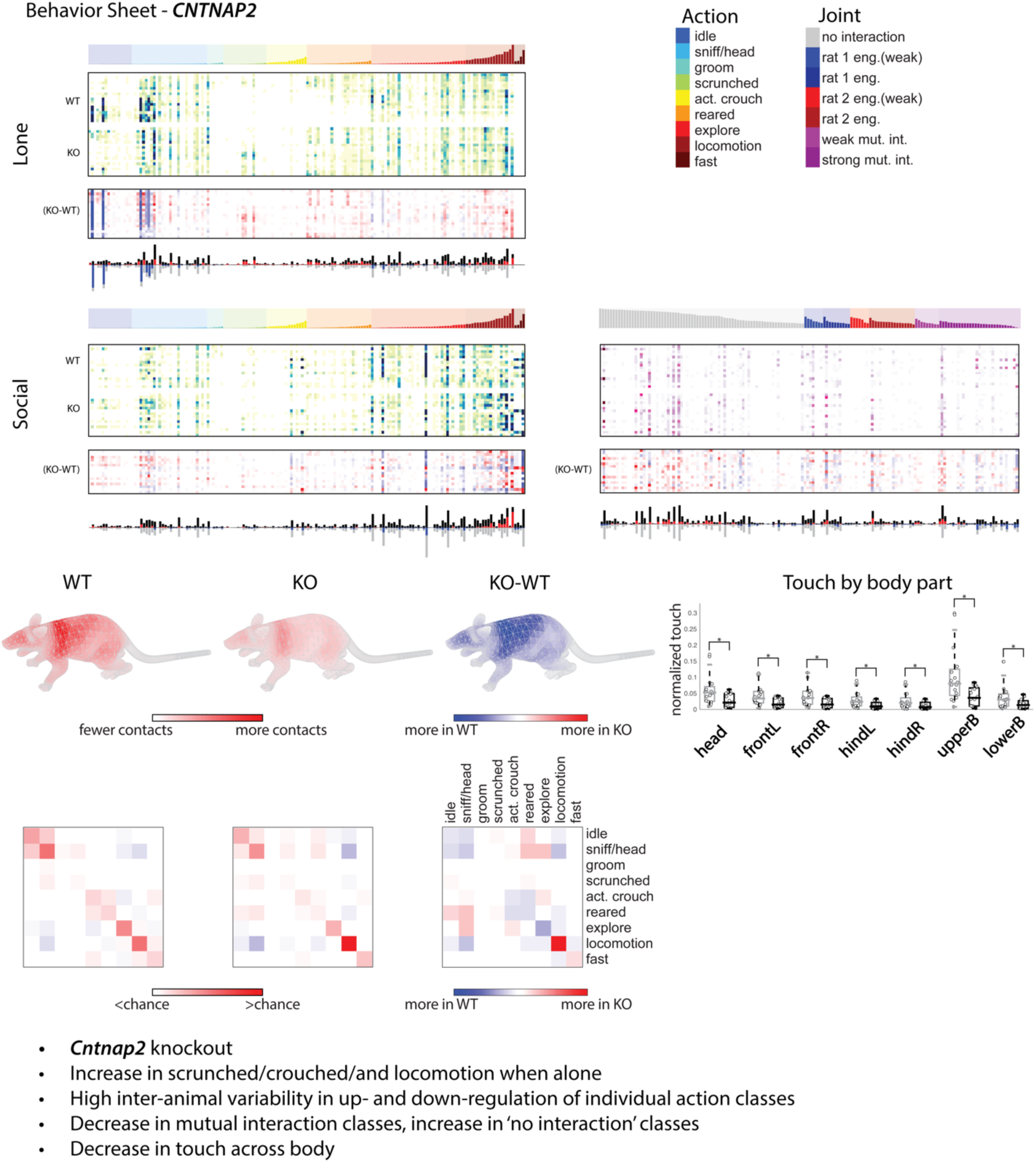
*CNTNAP* and wild-type littermate detailed behavior sheet. All raw usage for action and joint clusters is reported. Average differences from mean wild type behavior for each recording are shown. Raw touch and difference for wild-type (WT) and knockout (KO) are shown. The difference in total touch for WT and KO recordings for each body part is shown and significance is indicated with an asterisk. Mutual information calculated for WT/WT and KO/KO pairings and differences between the two are shown. Summary of findings is reported.

**Supplementary Figure 22:**
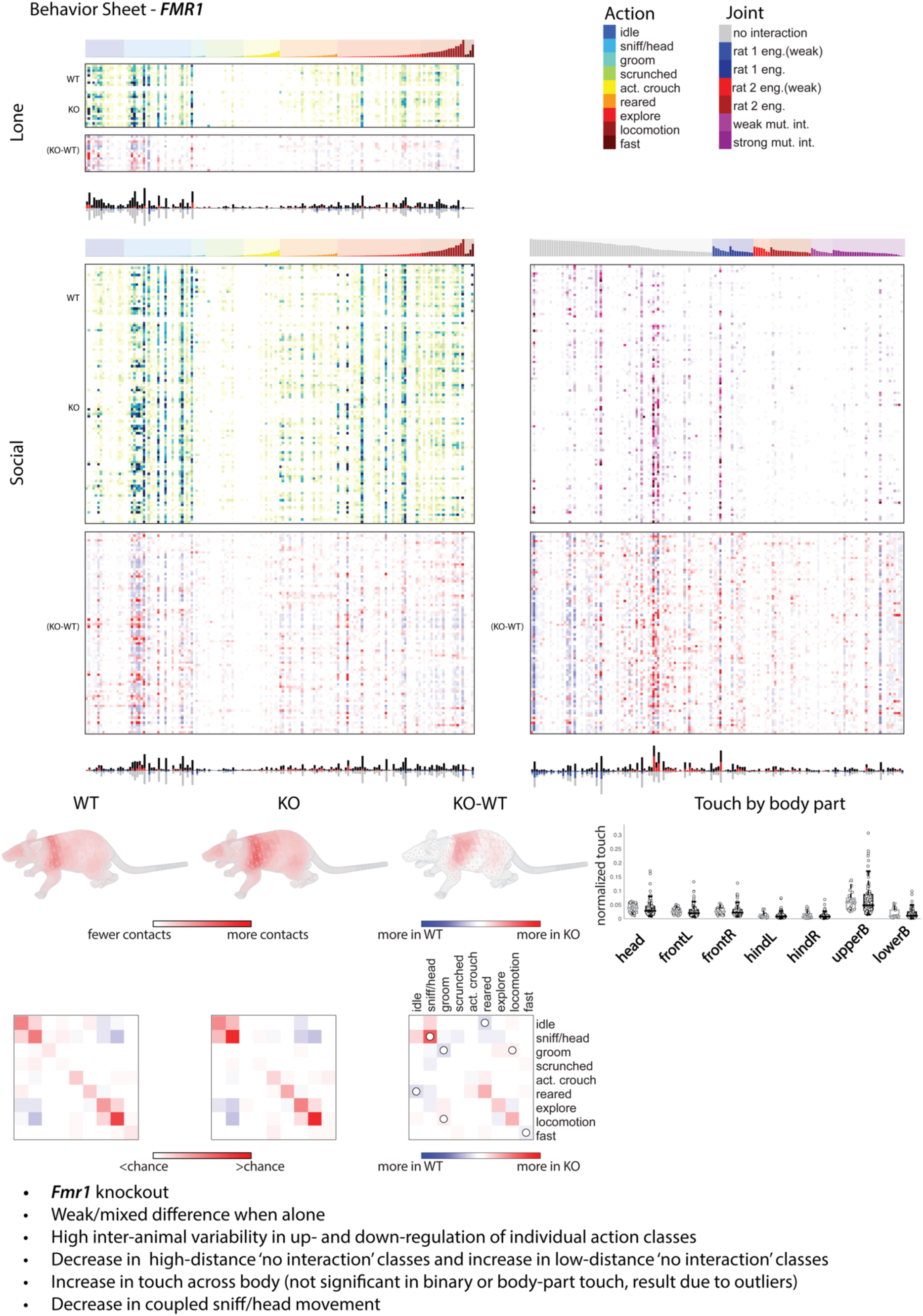
*FMR1* and wild-type littermate detailed behavior sheet. All raw usage for action and joint clusters is reported. Average differences from mean wild type behavior for each recording are shown. Raw touch and difference for wild-type (WT) and knockout (KO) are shown. The difference in total touch for WT and KO recordings for each body part is shown and significance is indicated with an asterisk. Mutual information calculated for WT/WT and KO/KO pairings and differences between the two are shown. Summary of findings is reported.

**Supplementary Figure 23:**
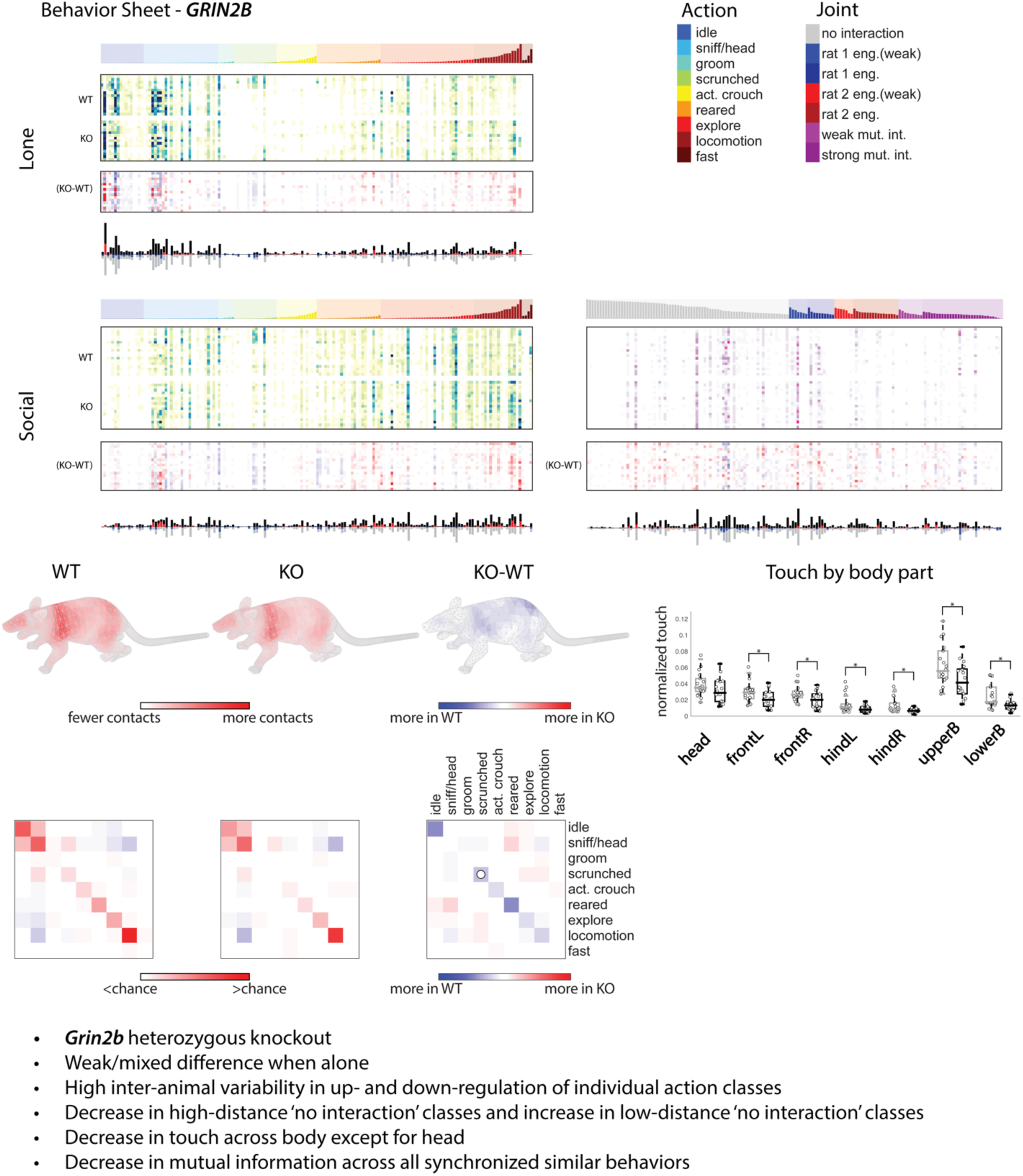
*GRIN2B* and wild-type littermate detailed behavior sheet. All raw usage for action and joint clusters is reported. Average differences from mean wild type behavior for each recording are shown. Raw touch and difference for wild-type (WT) and knockout (KO) are shown. The difference in total touch for WT and KO recordings for each body part is shown and significance is indicated with an asterisk. Mutual information calculated for WT/WT and KO/KO pairings and differences between the two are shown. Summary of findings is reported.

**Supplementary Figure 24:**
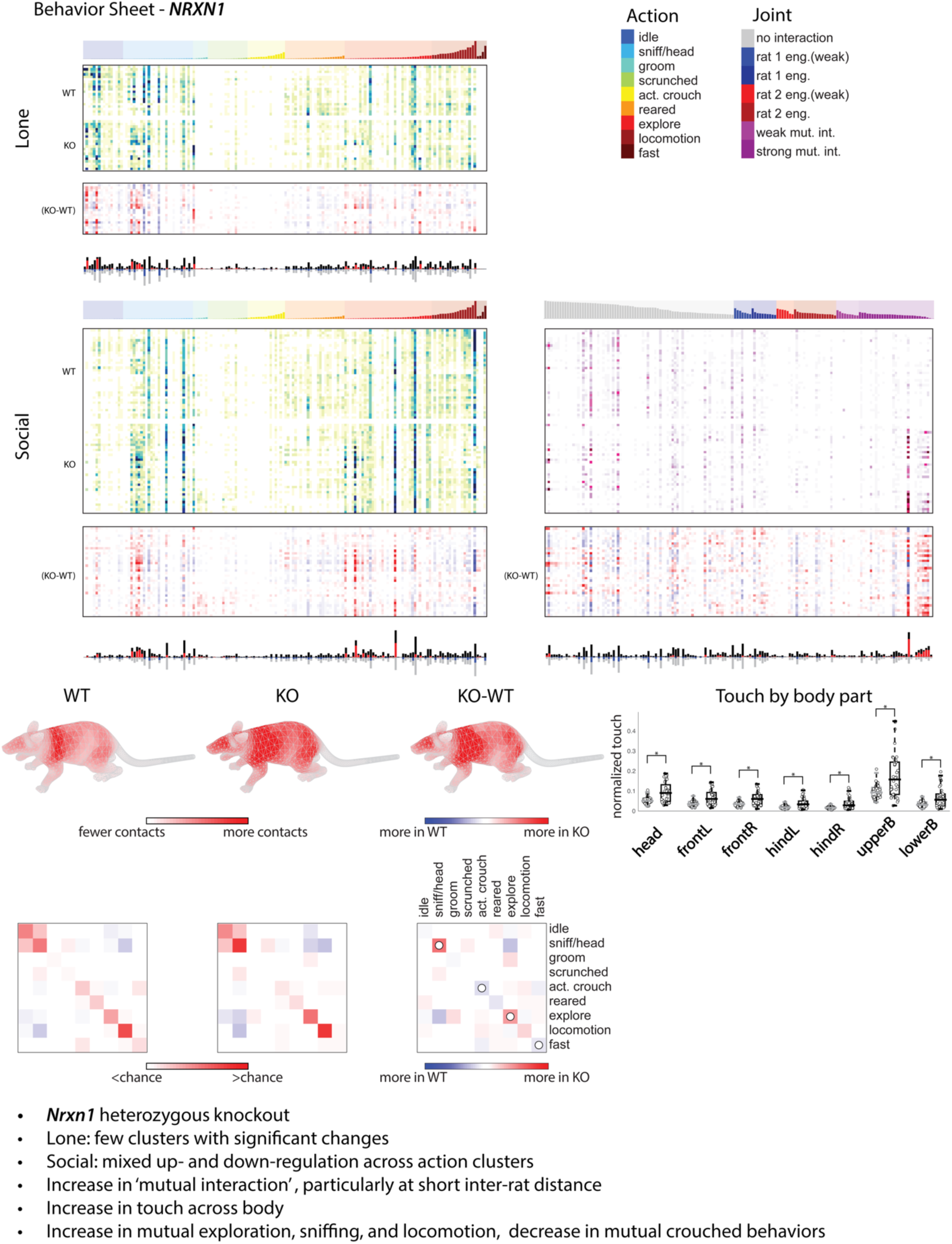
*NRXN1* and wild-type littermate detailed behavior sheet. All raw usage for action and joint clusters is reported. Average differences from mean wild type behavior for each recording are shown. Raw touch and difference for wild-type (WT) and knockout (KO) are shown. The difference in total touch for WT and KO recordings for each body part is shown and significance is indicated with an asterisk. Mutual information calculated for WT/WT and KO/KO pairings and differences between the two are shown. Summary of findings is reported.

**Supplementary Figure 25:**
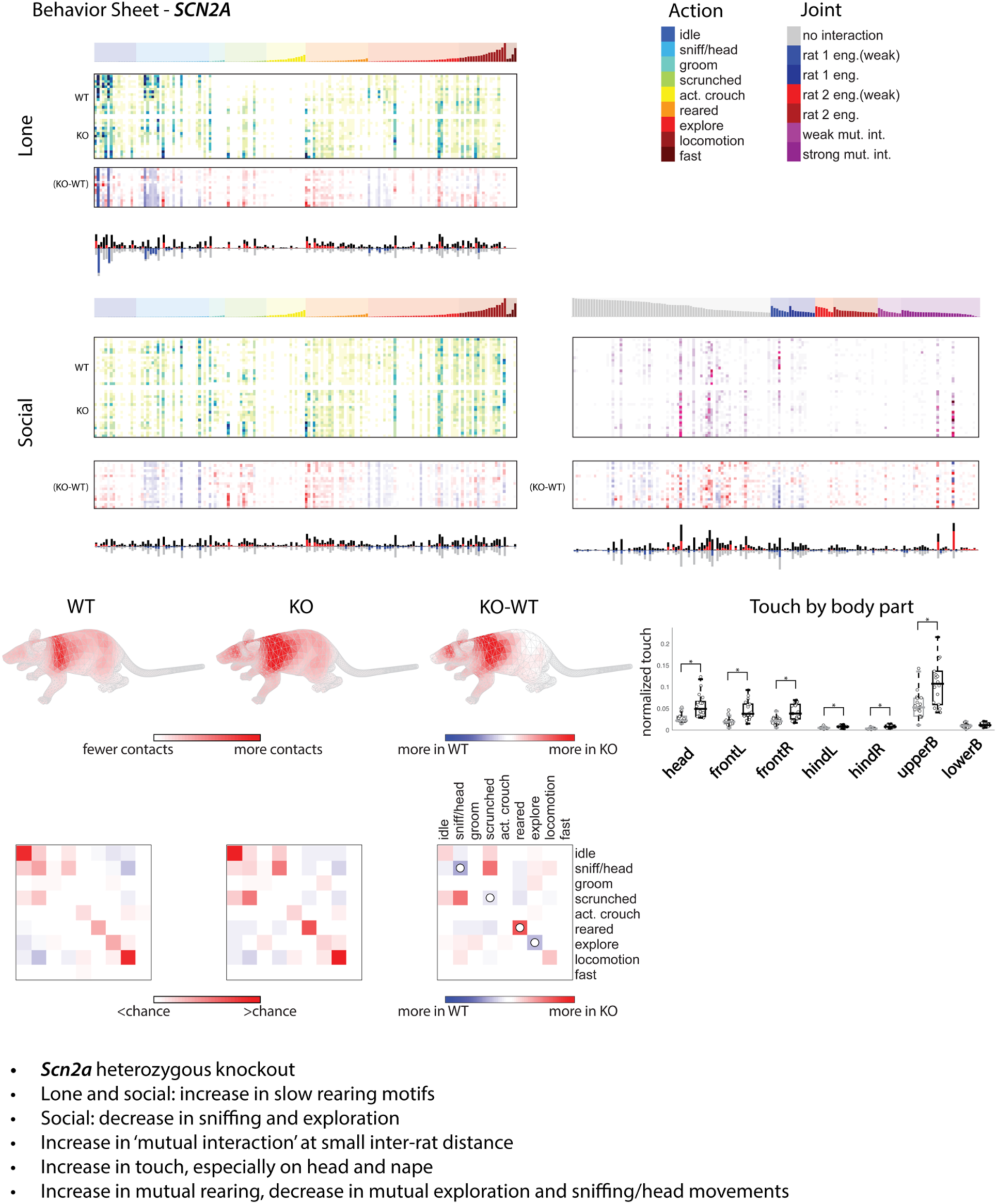
*SCN2A* and wild-type littermate detailed behavior sheet. All raw usage for action and joint clusters is reported. Average differences from mean wild type behavior for each recording are shown. Raw touch and difference for wild-type (WT) and knockout (KO) are shown. The difference in total touch for WT and KO recordings for each body part is shown and significance is indicated with an asterisk. Mutual information calculated for WT/WT and KO/KO pairings and differences between the two are shown. Summary of findings is reported.

